# Light contamination in stable isotope-labelled internal peptide standards is frequent and a potential source of false discovery and quantitation error in proteomics

**DOI:** 10.1101/2021.11.08.467684

**Authors:** Mogjiborahman Salek, Jonas D. Förster, Wolf-Dieter Lehmann, Angelika B. Riemer

## Abstract

In mass spectrometry-based proteomics, heavy internal standards are used to validate target peptide detections and to calibrate peptide quantification. Here we report light contamination present in heavy labelled synthetic peptides of high isotopic enrichment. Application of such peptides as assay-internal standards potentially compromises the detection and quantification especially of low abundant cellular peptides. Therefore, it is important to adopt guidelines to prevent false-positive identifications of endogenous light peptides as well as errors in their quantification from biological samples.

## INTRODUCTION

Stable isotope labelling (SIL) of amino acids in general starts by growing microorganisms (e.g. green algae, cyanobacteria) in the presence of highly ^13^C-enriched CO_2_ and/or highly ^15^N-enriched ammonia (each at >99.5 atom% enrichment) as sole source of carbon and nitrogen, respectively [1,2]. Labelled amino acids obtained in this way are then incorporated into peptides either by chemical synthesis or metabolic labelling [3]. Peptides containing at least one amino acid highly enriched in ^13^C and/or ^15^N are often named “heavy” to set them off against their cognates with natural isotopic abundances, named “light” in this context.

In targeted mass spectrometry (MS)-based proteomics and in particular peptidomics studies, heavy internal standards are regarded as the current gold-standard to validate peptide detection and quantitation. For example, with peptides presented by human leukocyte antigen (HLA) class I molecules, this identification is directly linked to subsequent immunotherapy and anti-cancer vaccine design [4-6]. The challenge in achieving accurate quantification is to effectively eliminate systematic and random errors due to various instrumental parameters or matrix effects. Isotope dilution MS (IDMS) based on heavy internal standards fits this purpose and provides proof of detection and absolute quantification of the highest precision and accuracy fit for clinical validation [7-9]. More recently, internal standards were used to dynamically control signal acquisition for endogenous light peptides via live instrument control [10]. With these broad applications in the field of mass spectrometry and in particular those assays aiming at clinical translation, high isotopic purity of heavy peptides used as internal reference is of key importance.

However, we here report that heavy peptides generated by chemical synthesis frequently contain sufficient levels of light cognates to bias interpretation of MS data, thus wrongly validating the detection of cellular peptides.

## MATERIALS AND METHODS

### Synthetic peptides and sample preparation

Peptide Retention Time Calibration (PRTC) Mixture (88321, Pierce™) was purchased from Thermo Fisher Scientific and is prepared at the same quality standards as “AQUA Ultimate” grade peptides of high chemical purity (>99%). Custom-synthetized heavy peptides were obtained from JPT Peptide Technologies (Berlin, Germany) and Synpeptide Co., Ltd (Shanghai, China) for low chemical purity (>70%) The above vendors are indicated in Table S1 as vendor 1, 2 and 3, respectively. For all vendors, the isotope purity is reported >99 atom% ^13^C and ^15^N, and incorporated heavy amino acids are V (^13^C_5_, ^15^N_1_), L/I (^13^C_6_, ^15^N_1_), K (^13^C_6_, ^15^N_2_) and R (^13^C_6_, ^15^N_4_). Upon arrival, peptides were dissolved in DMSO at 0.5nmol/μl, brought to a final concentration of 5 pmol/μl, dried, and stored at –80°C until LC-MS analysis.

### Immunoprecipitation of HLA-displayed peptides

HLA immunoprecipitation (IP) was performed according to previously published protocols [11,12]. Briefly, 1×10^8^ C33A cells, a HPV-negative cervical cancer cell line, were lysed with a lysis buffer containing 1% N-octyl-β-D glucopyranoside, 0.25% Na-Deoxycholate, protease inhibitor cocktail (Sigma-Aldrich, Mannheim, Germany) /PMSF (Carl Roth, Karlsruhe, Germany) in PBS. After centrifugation at 40,000xg, 4°C, for 30 min, HLA-peptide complexes were immuno-isolated by incubation with BB7.2 mouse anti-human HLA-A2 monoclonal antibody crosslinked to protein G Sepharose beads (Cytiva, Marlborough, MA, USA) for 4h at 4°C under constant mixing on a rotating wheel. Supernatant was discarded after centrifugation at 3200xg, for 3 min at RT. Pelleted HLA-peptide complexes bound to antibody-beads were washed 3 times in each of the following steps: first with ice-cold 20mM Tris-HCl pH 8 containing 150mM NaCl, followed by the same buffer but with 400 NaCl, and finally with 20mM Tris-HCl alone. Peptides were eluted from HLA bound to antibody-beads by 0.3% TFA. Resulting peptides were desalted by reverse-phase purification using a SepPak 96-well plate and dried by vacuum centrifugation (Concentrator plus, Eppendorf, Hamburg, Germany).

### LC-MS

Samples were dissolved in 5% ACN, 0.1% TFA. Synthetic peptides were injected up to 280 fmol per peptide on column. All samples were analysed by liquid chromatography (U-3000, Thermo Fisher Scientific) coupled to Q Exactive HFX or Orbitrap Exploris 480 (Thermo Fisher Scientific). The LC gradient consisted of a first segment going from 3% to 7.5% B (19.9% H_2_O, 0.1% FA, 80% ACN) and 92,5% A (0.1% FA in H_2_O) over 5 min, followed by a second segment reaching 35.6% B in 75 min. Finally, the % B was increased in 2 steps to 95% in 4 min, followed by 5 min wash before 11 min equilibration at 2% B. For quantifying light contamination, Orbitrap Exploris was operated with multiplexed tSIM scans for simultaneous mass measurement of multiple peptide precursor ions using 240k resolution at 200 m/z, 100 ms injection time (IT), 100% AGC target and 2-3 separate quadrupole isolation windows, focused on the heavy and the light precursors. Protonated polycyclodimethylsiloxane (PCM-6, a background ion originating from ambient air) at 445.12 m/z served as a lock mass. MS2 data was acquired with PRM scans using 60k resolution at 200 m/z with a narrow <1 m/z isolation window tuned per precursor. Heavy precursor settings were 30ms IT, 50% AGC target. Light precursor masses were measured with 1000ms IT, 1000% AGC target. These settings are identical to those we use for the analysis of endogenous peptides in a biological sample.

To test the matrix effect, either 10 or 40 ng HeLa Protein Digest Standard (Pierce™, Thermo Scientific™) served as a test matrix which were spiked with 150 fmol (Fig 2 a), or 100 fmol (Fig 2 b c) of heavy target peptides. As a control, an IP sample obtained as described above was spiked with 50 fmol heavy target peptides (Fig 2 d). MS2 data was acquired with PRM scans using 60k resolution at 200 m/z with a narrow < 1 m/z isolation window tuned per precursor. To mirror the conditions of analysis of a real biological sample, the IP sample from C33A cells spiked with heavy peptides was analysed with parameters for heavy and light precursor mass measurements at 150 ms IT and 100% AGC target for the former, 1000 ms and 1000% for the latter. To explore the impact of the matrix effect on the sensitivity of peptide detection, MassPREP™ E.coli digestion standard (MPDS E.coli, Water, Milford, MA, USA) was used in the specified amounts to test the detection of the peptides from vendors 1 and 2 (Fig 2 e-g, Table S1). This data was acquired on a Q Exactive HF-X with an MS scan followed by 8 PRM scans, each with 0.7 m/z isolation window for a maximum of 350 ms IT and 5e5 ions AGC target with 23% normalized collision energy (NCE).

### Data Analysis

MS1 data analysis was performed with R (v. 4.1) [13] with the “tidyverse” suite of packages (v. 1.3.0) [14]. Theoretical isotopic envelopes were calculated with the IsoSpec R package (v. 2.1.3) [15]. For theoretical heavy envelopes, several isotopic purity settings for heavy isotopes ^13^C and ^15^N were probed and 97% isotopic purity was selected for best fits. Theoretical isotopic envelopes and tSIM spectra were centroided and matched with 4 ppm mass tolerance. The matched envelope was then fitted with a linear model, minimizing the sum of squares error.

MS2 data was analysed with the Skyline software (v. 20.2) [16]. Top 4 intensity product ions were extracted with 10 ppm mass tolerance. Detected peaks were manually curated. Light peaks were discarded when peak retention times or shapes did not match with the heavy reference, when the normalized spectral contrast angle (dotp) [17] was low, or when too few transitions were detected. For the complex peptide matrix (CPM) analysis, the “nls.multstart” (v. 1.2.0) R package [18] was used to fit the response of the peptide MS2 intensity (I) to the CPM load with the function I = *a* ([CPM]*b*)10^^^*c*.

## RESULTS AND DISCUSSION

The sequences and extent of light contamination in the synthetic heavy peptides we tested are given in full in supplementary Table S1 and Figure S1. These different sets of peptides were in use at the time in our laboratory and reflect different suppliers and purities. They include 9-15 amino acids long human leukocyte antigen (HLA) binders or tryptic peptides containing 1 or 2 labelled residues.

Representative for the peptides of highest available quality (99.5% isotope purity, >97% chemical purity, concentration precision ±10%, absolute quantification (AQUA) grade), peptide #43 GISNEGQNASI<u>K</u> shows precursor isotopic envelopes at both light (monoisotopic 0) and heavy (monoisotopic +8) m/z (Figure 1a). The strong similarity of the fragmentation patterns between the light and heavy monoisotopic precursor ions in comparison to an external library, as captured by high spectral contrast angle (dotp) values (0.96, 0.98), confirms the sequence identity of the light and heavy peptides (Figure 1b). The correct peptide identification is further supported by the alignment of their retention times, indicating virtually perfect chromatographic co-elution. The light contamination of peptide #43 GISNEGQNASI<u>K</u> is quantified as 219 ppm according to the MS2 peak area of the shown transitions (Figure 1c and Figure S1, for other peptides from this set: #40 (4507 ppm) to #54 (25 ppm)).

**Figure 1.**
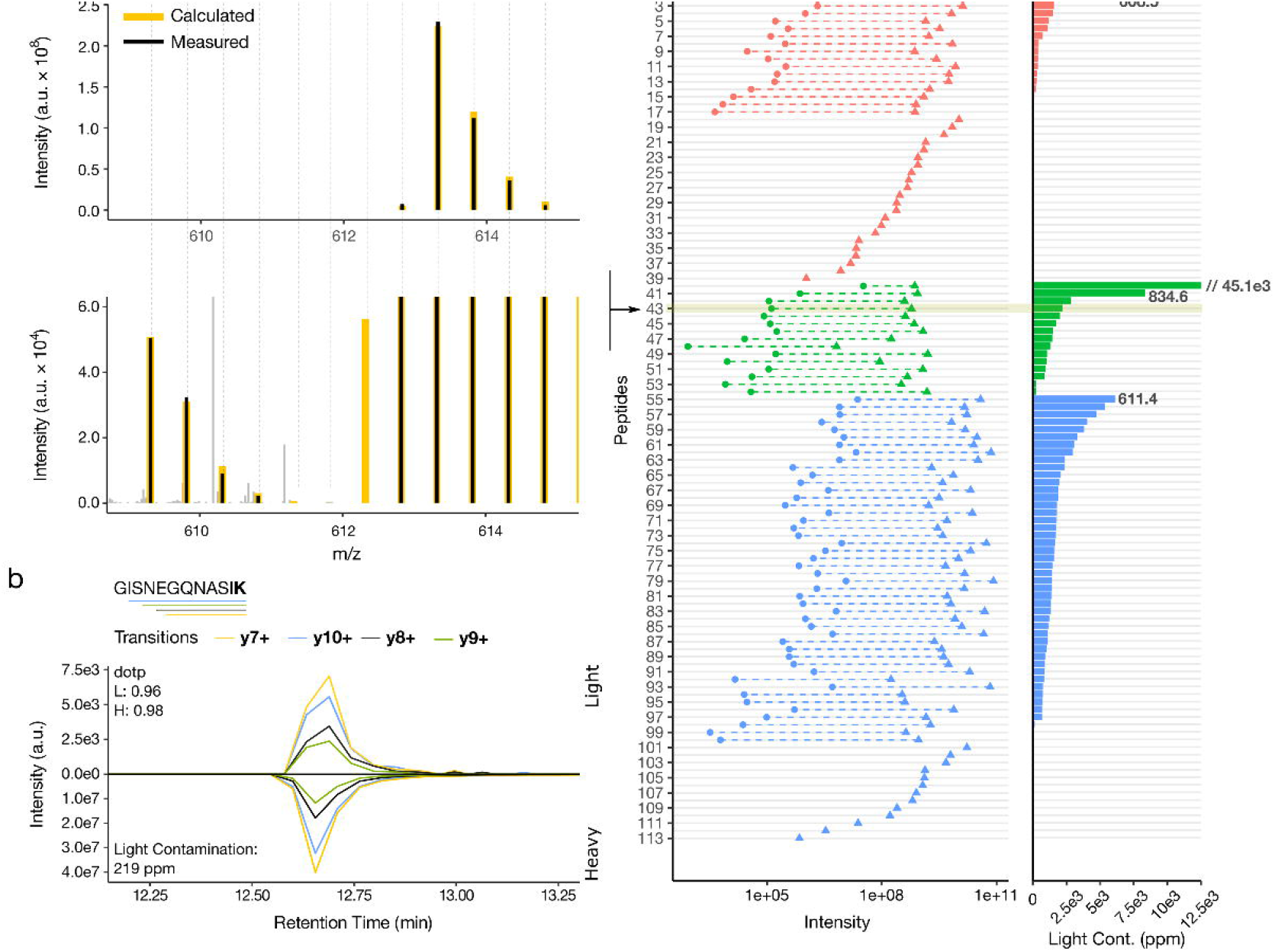
Presence of light contamination in heavy peptides. (a) Upper and lower panels show selected ion monitoring (SIM) data acquisition scans for the light and heavy forms of a representative peptide and its 4000x zoom in, respectively. The calculated (yellow) and experimentally measured (black) isotope patterns of the labelled peptide and its light contamination are displayed. The indicated number of heavy isotopes, “0” and “+8”, corresponds to the light and heavy monoisotopic signals, respectively. Background peaks not matched to the isotope pattern are displayed in grey (b) Extracted fragment-ion chromatograms representing the light and heavy forms of the depicted peptide sequence (the labelled residue, lysine (K), is in bold, for which the sum of incorporated stable isotopes is 8 (^13^C_6_ and ^15^N_2_), as indicated in (a)). The bars underneath the peptide sequence correspond to the detected fragments or transitions and are color-coded accordingly. dotp is the normalized spectral contrast angle that is scoring the similarity of the detected peptide’s fragmentation pattern of light (L) and heavy (H) precursors, compared to a reference library. The identity of the peptide is further confirmed by the co-elution of the light and heavy signals. (c) Extent of light contamination (L/H) for 113 tested heavy peptides, from three vendors (for colour scheme see Table S1). The light contamination for each peptide is shown in the left panel as the summed peak area for the top 4 fragments of the light (dots) and heavy (triangles) peptides. The right panel shows the extent of light contamination as parts per million (ppm). The arrow pinpoints the peptide #43 (219 ppm), for which the details of light contamination are shown in panels (a) and (b).

Although sources of such contamination can be multiple, the absence of a gradual increase in the intensity of intermediate signals between the light (0) and heavy (+8) monoisotopic peaks, excludes the possibility of incomplete incorporation of ^13^C and ^15^N isotopes during the initial metabolic labelling of the amino acids. More likely, the source of contamination would therefore be the introduction of completely unlabelled, *i*.*e*. “light”, amino acids from an external source during amino acid labelling, extraction or peptide synthesis.

In around 50% of all heavy synthetic peptides we analysed as mixtures within separate batches from different vendors, the level of light contamination was higher than 100 ppm (0.01%, Figure 1c). The highest level of contamination measured was 4.5% for peptide #40 TASEFDSAIAQD<u>K</u> belonging to the AQUA grade peptide mixture (Figure 1c and Table S1). Initial measurements indicated exceedingly high contamination of >900 ppm for a total of 6 peptides. When confirming these peptides individually, the contamination assessment could be corrected down by factors of 100 to 1500 for those peptides, where the source of contamination was tracked down to another peptide of related sequence in the same mixture (Table S1, peptides #92, #95, #99, #100, #113). For instance, the peptide #92 RALYVDSLFF initially showed contamination of 32812 ppm but could be corrected down to 84 ppm when measured separately, whereas the source of major contamination was determined to be the peptide #84 RALYVDSLFFL which introduced RALYVDSLFF as incomplete synthesis product. Thus, spike-in of such related sequences in a biological sample would be problematic. Nevertheless, it is important to note that the light contamination reported here cannot be attributed solely to cross-contamination due to mixing of peptides of low chemical purity. The separate analysis of single peptides still shows light contamination (Table S1, #1 K<u>L</u>GEIVTTI). In addition, when analysing three independent batches of the AQUA grade peptides (chemical purity >97%), the contamination remained high (#40: 4.5%, 1.1% or 0.04%, data not shown).

Data acquisition with instrument parameters tailored for high sensitivity inevitably increases the chances of detecting light contaminations as low as 50 ppm (Figure 2a, Table S1, #8 RTLED<u>L</u>LMGT). Similarly, in quantitative MS the well-documented matrix effect [19-21], often attributed to the changes in the efficiency of electrospray ionization, can enhance or suppress a target peptide signal. Indeed, we observed that the analysis of heavy peptides in the presence of a complex peptide matrix (CPM, here: HeLa tryptic digests), enhances both the signal of the heavy peptide and of its light contamination, which might have remained undetected in the absence of the matrix (Figure 2b). Increasing the amount of CPM showed similar effects as increasing instrument sensitivity of detection (Figure 2c). It has been suggested that consistent quantification requires analysis in a fixed amount of matrix [22], implying that in biological experiments negative controls including heavy internal standards need to be performed in the presence of cellular or closely related matrices. In the absence of such a control, peptide detection in a biological sample would lack sufficient level of objective evidence to distinguish the light contamination introduced by the heavy internal standard, from a *bona fide endogenous peptide* (Figure 2d).

**Figure 2.**
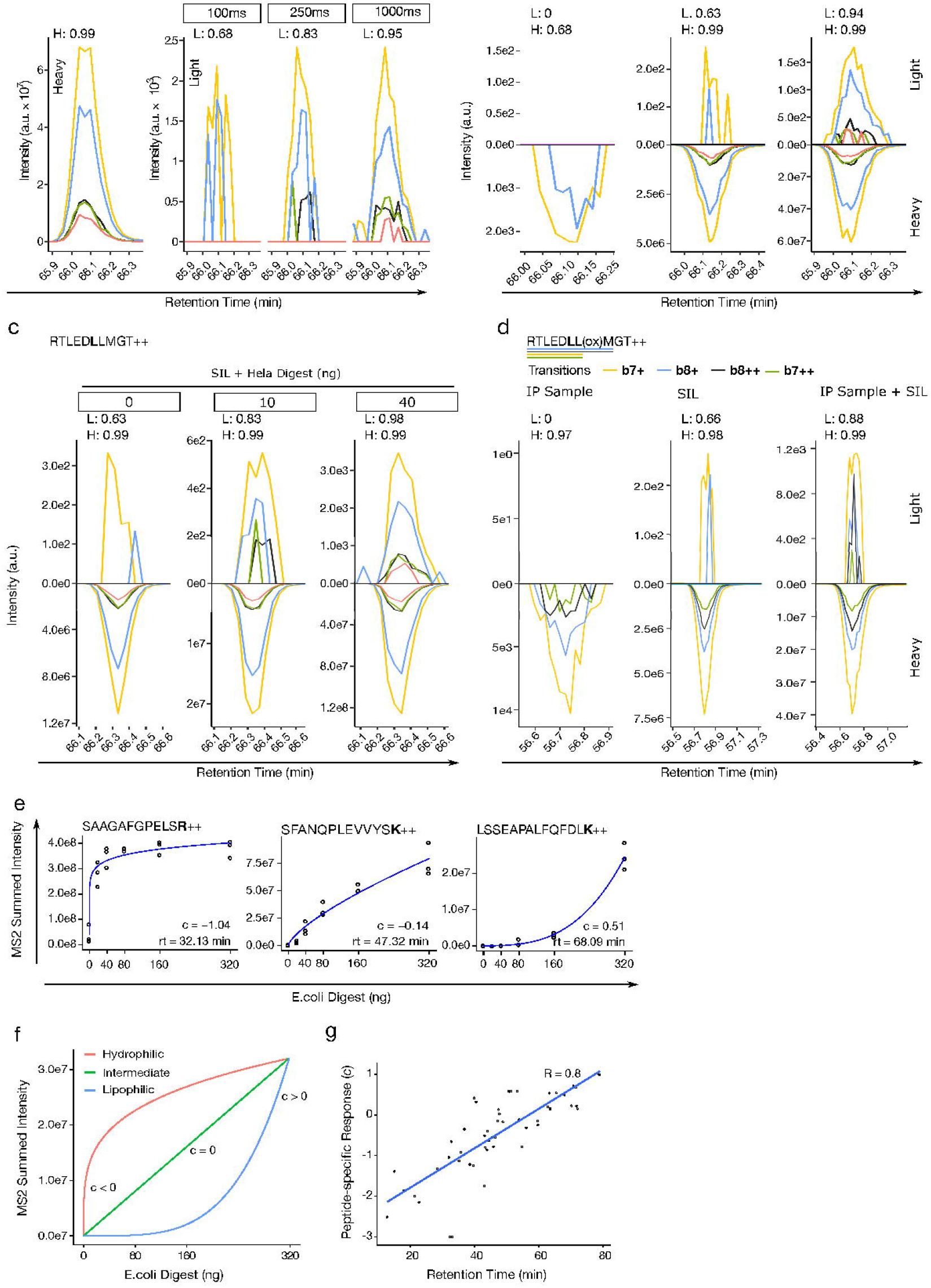
Injection time and matrix effects on the detection of light contamination. (a-d) Extracted ion chromatograms representing the transitions (fragments) corresponding to the depicted heavy peptide sequence and its light contamination detected in different conditions. dotp comparison to reference library is annotated for light (L) and heavy (H) signals. (a) Increasing of injection time (sensitivity); (b) presence or absence of a CPM; (c) increasing amounts of CPM; (d) in the context of anti-HLA immunoprecipitation (IP) sample. Oxidized peptide is targeted in this assay. Heavy reference signal in the IP sample is trace-level carry-over from other experiments on the same LC system. This is common for heavy peptides injected at relatively high concentrations. (e-g) Correlation between the chemical nature of target peptides and the amount of CPM. Three technical replicates per condition. The response of peptide intensity to CPM amount is fitted with I = *a* ([CPM]*b*)10^^^*c*; (e) Three representative peptides (hydrophilic, intermediate, lipophilic) for different behaviour in the presence of CPM. (f) The effect of factor c and the modes that were commonly observed for peptides of different hydrophobicity (g) The factor c shows a linear correlation with peptide retention time (RT) (Pearson correlation coefficient of 0.8).

To further investigate the observed CPM effect, we analysed 54 (Table S1, vendors 1 and 2) heavy peptides covering the entire chromatographic retention time range in the presence of increasing amounts of *E. coli* tryptic digests. The summed intensities of the top 6 fragment ions for three representative peptides with early, intermediate and late retention times indicate that hydrophilic peptides, represented by peptide #49 SAAGAFGPELS<u>R,</u> need much less CPM to reach their maximum intensity than hydrophobic peptides, represented by peptide #48 LSSEAPALFQFDL<u>K</u>. Collectively, the response of the intensity (I) for each of the 54 peptides to the amount of CPM is best captured by I = *a* ([CPM]*b*)10^^^*c*, where c is representing the curvature as negative, null or positive, a and b are scaling and offset factors, respectively. c delineates three responsiveness behaviours to the amount of CPM, as fast, linear and slow (Figure 2f). Indeed, this peptide-specific responsiveness correlates linearly with the chromatographic retention time of the peptides (Figure 2g), suggesting that the MS signal enhancement by CPM in our data is mainly caused by preventing peptide losses via hydrophobic interactions to vessel surfaces.

The consequences of light contamination present in heavy internal standards are even more important for interpretation of LC-MS data in biological contexts, as modern MS instrumentations are sensitive by design. Newly developed sophisticated data acquisition approaches [10,23-25] further increase the basic instrumental sensitivity and thereby the importance of taking into account the unappreciated source of error introduced by light contamination of heavy peptides. For instance, recently introduced intelligent instrument control improves IDMS by dynamic adjustment of detection parameters based on heavy internal standards. A low resolution survey of heavy standards added to the sample at a higher concentration will trigger real time measurement of light endogenous peptides of low abundance at high resolution and sensitivity [10]. Here, light contamination, introduced by the addition of heavy labelled peptides of seemingly high isotopic purity, can be easily mistaken as endogenous light peptides leading to false positive identifications and flawed quantifications.

We therefore recommend the adoption of a few simple and rational guidelines to prevent false-positive detection of endogenous peptides. First, the impact of light contamination in batch analyses with multiple peptides can be controlled by consistent design of the peptide labelling. For instance in the case of KSVLTAFLM<u>L</u>W and KS<u>V</u>LTAFLM, the first peptide contains an unlabelled segment identical to the second, which should be prevented by carefully selecting the site of labelling, and if necessary introducing a second labelling within the peptide sequence. Second, it is paramount to determine systematically that the employed LC-MS method does not produce any light target signal when performed with a suitable negative control matrix spiked with the heavy internal standard peptide of interest. Additionally, in targeted MS workflows, it is still common to use light synthetic peptides, physically indistinguishable from the endogenous targets, for the optimisation of instrument parameters. We strongly recommend that the LC-MS system is not exposed to light synthetic target peptides under any circumstances, as carry-over could again lead to false-positive detection.

## CONCLUSION

We here report that heavy synthetic peptides produced with the highest chemical and isotope purity can still contain substantial amounts of light cognates. The data was initially presented at the HUPO 2019 conference and its HUPO-HIPP satellite meeting in Adelaide [26]. To the best of our knowledge, this under-appreciated source of error is only described in one other publication, as one of several pitfalls that can obscure the analysis of HLA ligandome data [27]. We here provide an in-depth analysis of light contamination by i) testing heavy peptides from multiple providers, ii) measuring the overall extent of light contamination, and iii) describing the matrix effect that enhances the intensity of light contamination that otherwise may remain undetected in quality controls. Given the high sensitivity of modern mass spectrometers, light contamination of internal standards, even as low as 50 ppm, will inevitably lead to a high level of false discovery. In contrast to protein detection relying on multiple peptides, the impact of the neglected source of error reported here is expected to be large on the analysis of single peptides such as low abundance post-translational modifications in cellular signalling [28] and tumor-derived HLA-displayed peptides [5], often identified at the limit of detection. To prevent the generation of false-positive detections by such studies in the future, we recommend easy-to-implement protective measures.

## Acknowledgements

The authors gratefully acknowledge Rebecca Köhler for excellent technical assistance and Dr. Jonas Becker for careful proofreading and editing of the manuscript.

## Figure Legends

**Figure S1. Full set of extracted ion chromatograms acquired for light contamination assessment**.

Extracted ion chromatograms for heavy (bottom) and light (top) precursors. Where appropriate, panels showing peptide chromatograms from single peptide solutions are given next to chromatograms acquired from the full peptide mixture. dotp and ppm assessment of heavy vs light is calculated from the summed peak area of the top 4 transitions of the displayed chromatograms. Where light contamination signal was not selected as identified, ppm and dotp are given as “NA” (not available).

**Table S1. Full list of peptides acquired in this study and the assessed ppm quantification for the light contamination**.

Peptide numbering reflects the position in Figure 1c. Vendor information is given as specified in the experimental section. Light contamination signal is given as fraction of light precursor peak area vs heavy precursor peak area. Separate columns list measurements where peptide contaminations were confirmed individually as opposed to bulk measurement in mixture.

