## Supplementary Figure 1 for "Light contamination in stable isotope-labelled internal peptide standards is frequent and a potential source of false discovery and quantitation error in proteomics"

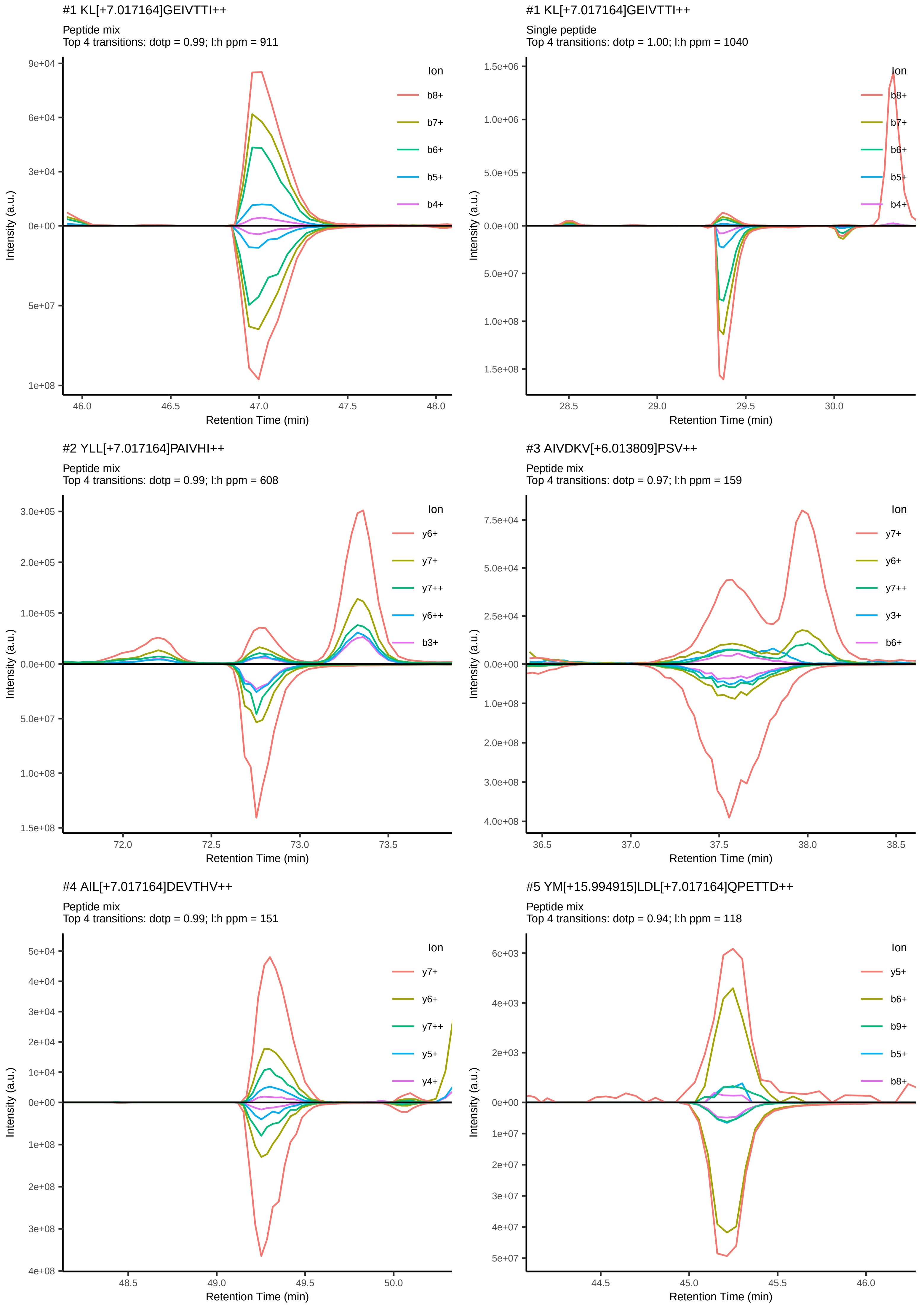
#1 KL[+7.017164]GEIVTTI++
Peptide mix
Top 4 transitions: dotp = 0.99; l:h ppm = 911
9e+04
6e+04
3e+04
0e+00
5e+07
1e+08
Intensity (a.u.)
46.0
46.5
47.0
47.5
48.0
Retention Time (min)
Ion
b8+
b7+
b6+
b5+
b4+
#1 KL[+7.017164]GEIVTTI++
Single peptide
Top 4 transitions: dotp = 1.00; l:h ppm = 1040
1.5e+06
1.0e+06
5.0e+05
0.0e+00
5.0e+07
1.0e+08
1.5e+08
Intensity (a.u.)
28.5
29.0
29.5
30.0
Retention Time (min)
Ion
b8+
b7+
b6+
b5+
b4+
#2 YLL[+7.017164]PAIVHI++
Peptide mix
Top 4 transitions: dotp = 0.99; l:h ppm = 608
3.0e+05
2.0e+05
1.0e+05
0.0e+00
5.0e+07
1.0e+08
1.5e+08
Intensity (a.u.)
72.0
72.5
73.0
73.5
Retention Time (min)
Ion
y6+
y7+
y7++
y6++
b3+
#3 AIVDKV[+6.013809]PSV++
Peptide mix
Top 4 transitions: dotp = 0.97; l:h ppm = 159
7.5e+04
5.0e+04
2.5e+04
0.0e+00
1.0e+08
2.0e+08
3.0e+08
4.0e+08
Intensity (a.u.)
36.5
37.0
37.5
38.0
38.5
Retention Time (min)
Ion
y7+
y6+
y7++
y3+
b6+
#4 AIL[+7.017164]DEVTHV++
Peptide mix
Top 4 transitions: dotp = 0.99; l:h ppm = 151
5e+04
4e+04
3e+04
2e+04
1e+04
0e+00
1e+08
2e+08
3e+08
4e+08
Intensity (a.u.)
48.5
49.0
49.5
50.0
Retention Time (min)
Ion
y7+
y6+
y7++
y5+
y4+
#5 YM[+15.994915]LDL[+7.017164]QPETTD++
Peptide mix
Top 4 transitions: dotp = 0.94; l:h ppm = 118
6e+03
4e+03
2e+03
0e+00
1e+07
2e+07
3e+07
4e+07
5e+07
Intensity (a.u.)
44.5
45.0
45.5
46.0
Retention Time (min)
Ion
y5+
b6+
b9+
b5+
b8+

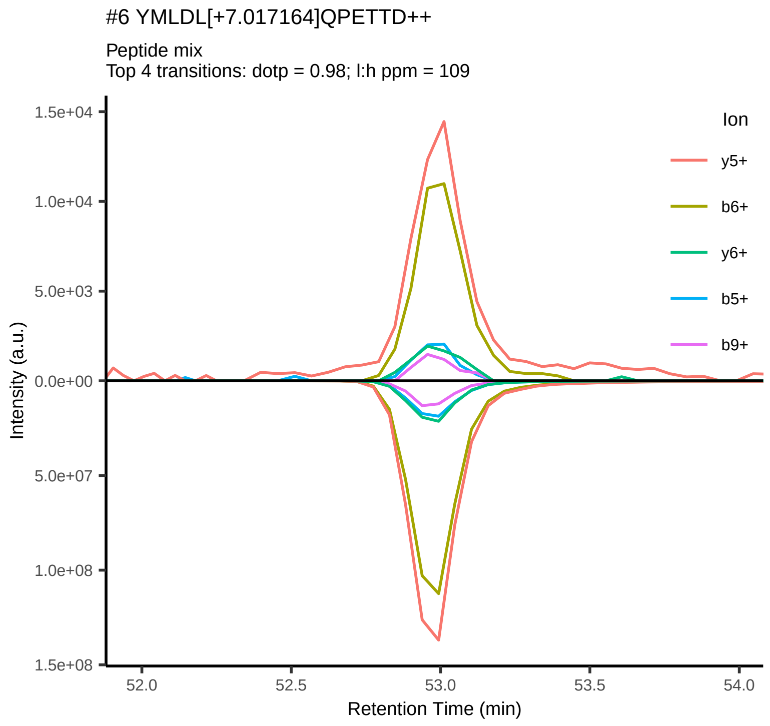
#6 YMLDL[+7.017164]QPETTD++
Peptide mix
Top 4 transitions: dotp = 0.98; l:h ppm = 109
Ion
y5+
b6+
y6+
b5+
b9+
1.5e+04
1.0e+04
5.0e+03
0.0e+00
5.0e+07
1.0e+08
1.5e+08
Intensity (a.u.)
52.0
52.5
53.0
53.5
54.0
Retention Time (min)

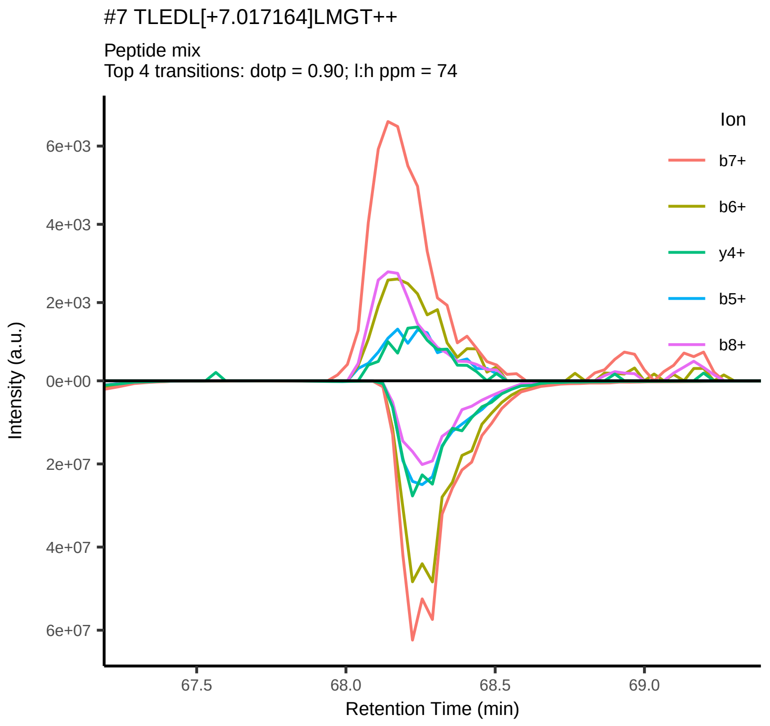
#7 TLEDL[+7.017164]LMGT++
Peptide mix
Top 4 transitions: dotp = 0.90; l:h ppm = 74
Ion
b7+
b6+
y4+
b5+
b8+
6e+03
4e+03
2e+03
0e+00
2e+07
4e+07
6e+07
Intensity (a.u.)
67.5
68.0
68.5
69.0
Retention Time (min)

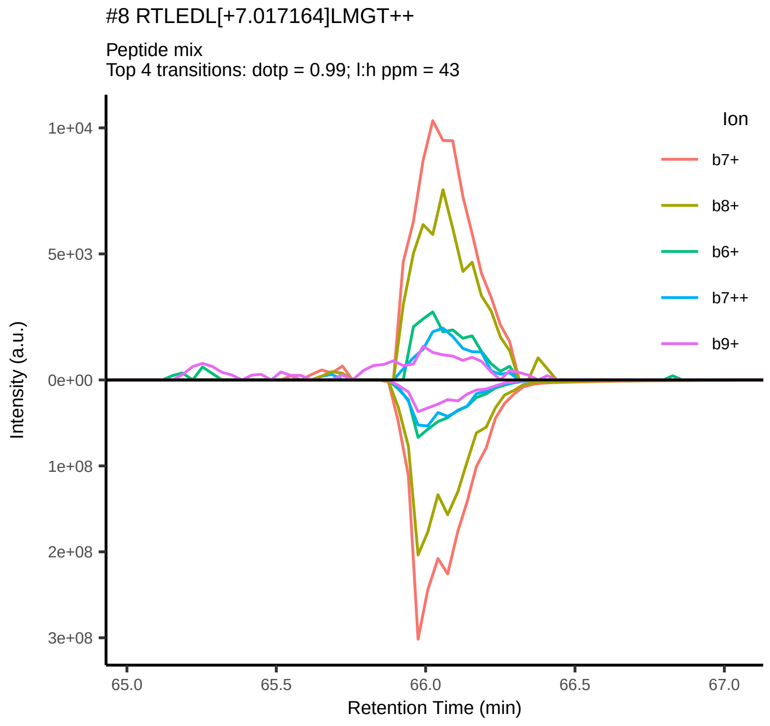
#8 RTLEDL[+7.017164]LMGT++
Peptide mix
Top 4 transitions: dotp = 0.99; l:h ppm = 43
Ion
b7+
b8+
b6+
b7++
b9+
1e+04
5e+03
0e+00
1e+08
2e+08
3e+08
Intensity (a.u.)
65.0
65.5
66.0
66.5
67.0
Retention Time (min)

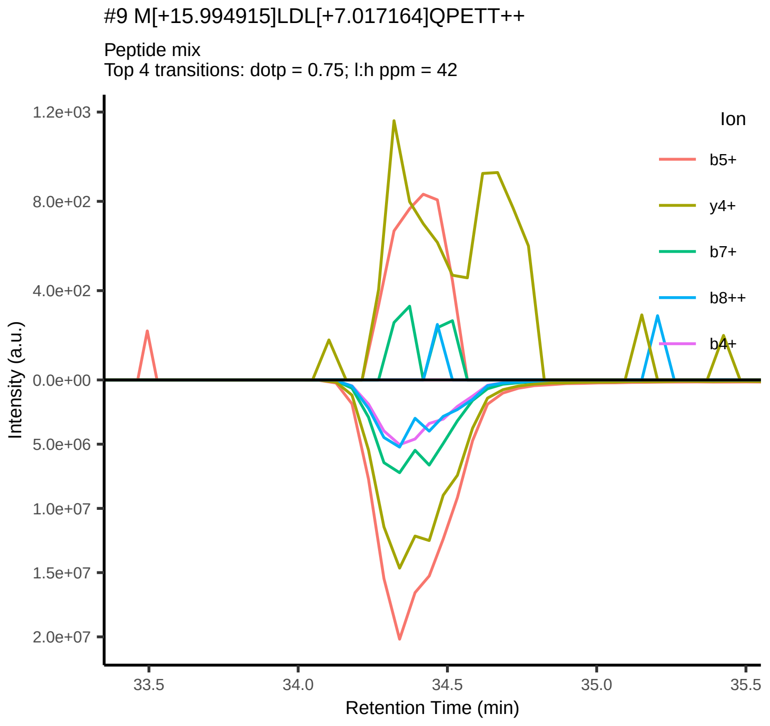
#9 M[+15.994915]LDL[+7.017164]QPETT++
Peptide mix
Top 4 transitions: dotp = 0.75; l:h ppm = 42
Ion
b5+
y4+
b7+
b8++
b4+
1.2e+03
8.0e+02
4.0e+02
0.0e+00
5.0e+06
1.0e+07
1.5e+07
2.0e+07
Intensity (a.u.)
33.5
34.0
34.5
35.0
35.5
Retention Time (min)

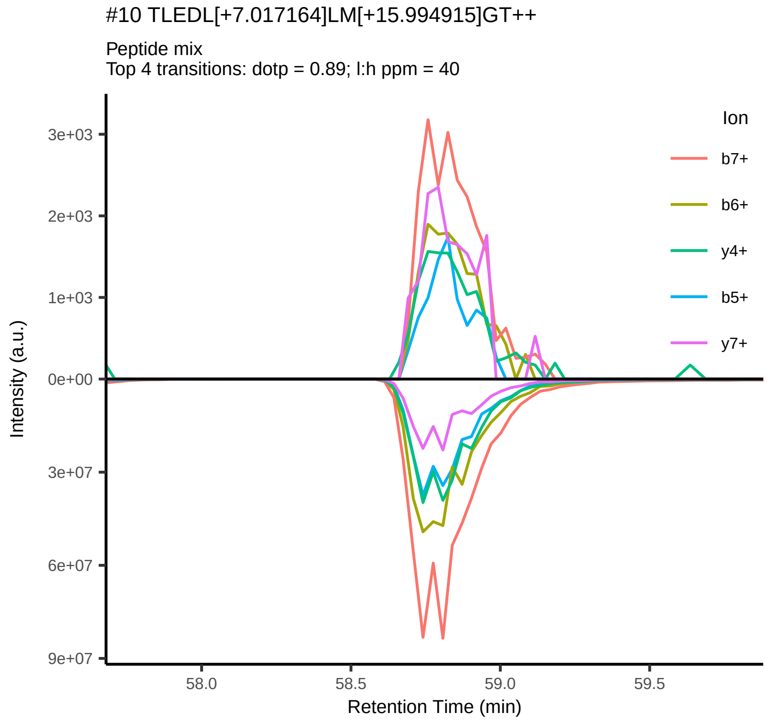
#10 TLEDL[+7.017164]LM[+15.994915]GT++
Peptide mix
Top 4 transitions: dotp = 0.89; l:h ppm = 40
Ion
b7+
b6+
y4+
b5+
y7+
3e+03
2e+03
1e+03
0e+00
3e+07
6e+07
9e+07
Intensity (a.u.)
58.0
58.5
59.0
59.5
Retention Time (min)

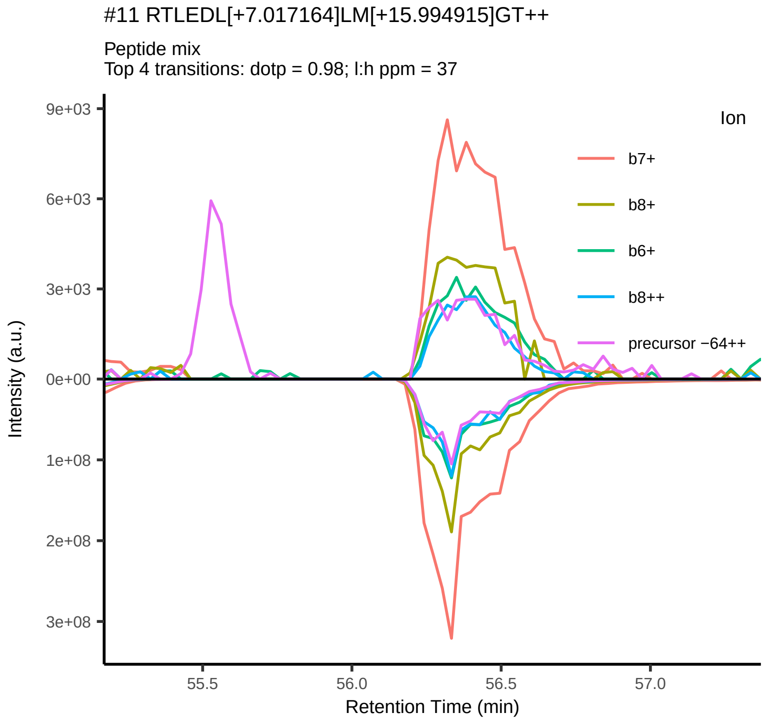
#11 RTLEDL[+7.017164]LM[+15.994915]GT++
Peptide mix
Top 4 transitions: dotp = 0.98; l:h ppm = 37
Ion
b7+
b8+
b6+
b8++
precursor −64++
9e+03
6e+03
3e+03
0e+00
1e+08
2e+08
3e+08
Intensity (a.u.)
55.5
56.0
56.5
57.0
Retention Time (min)

#12 IRTLEDL[+7.017164]LM[+15.994915]GT++

Peptide mix
Top 4 transitions: dotp = 0.98; l:h ppm = 32

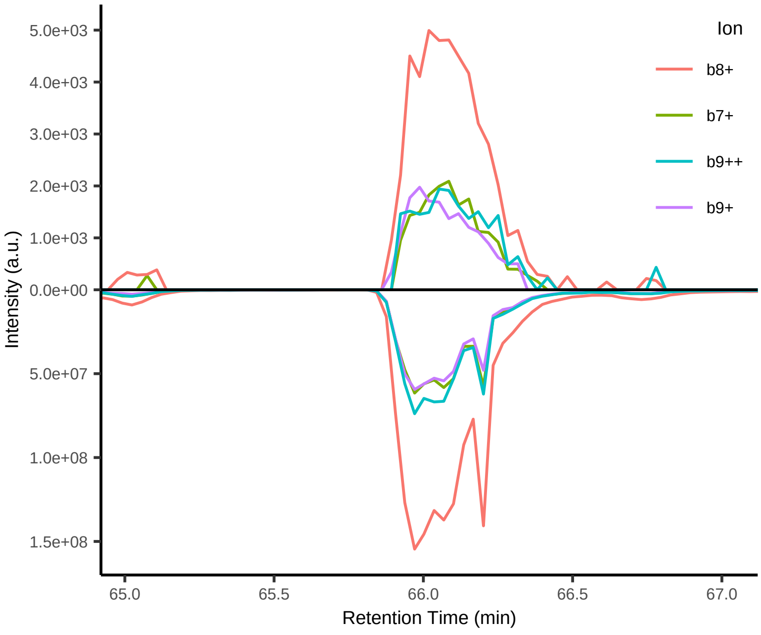


#13 IRTLEDL[+7.017164]LMGT++

Peptide mix
Top 4 transitions: dotp = 0.97; l:h ppm = 28

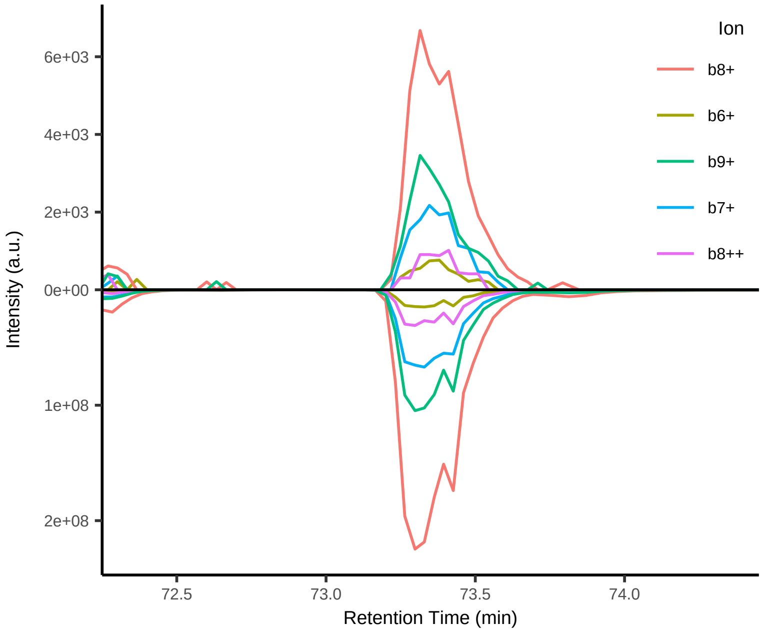


#14 YM[+15.994915]LDL[+7.017164]QPETT++

Peptide mix
Top 4 transitions: dotp = 0.93; l:h ppm = 21

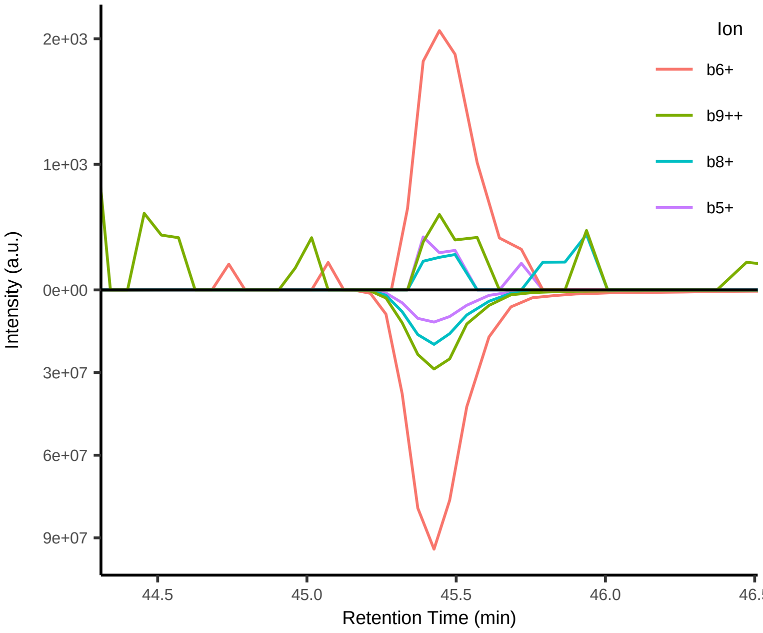


#15 LLMGTLGI[+7.017164]V+

Peptide mix
Top 4 transitions: dotp = 0.91; l:h ppm = 11

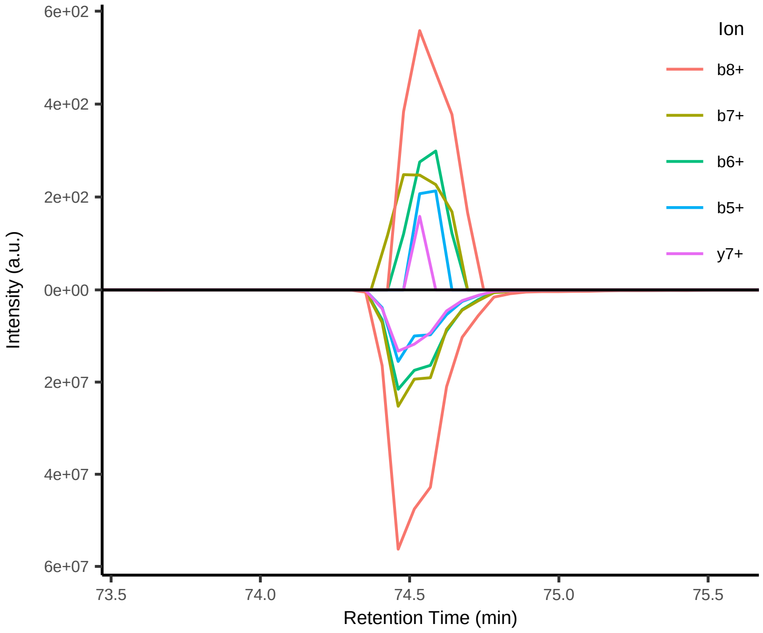


#16 DLLMGTLGI[+7.017164]V++

Peptide mix
Top 4 transitions: dotp = 0.73; l:h ppm = 9

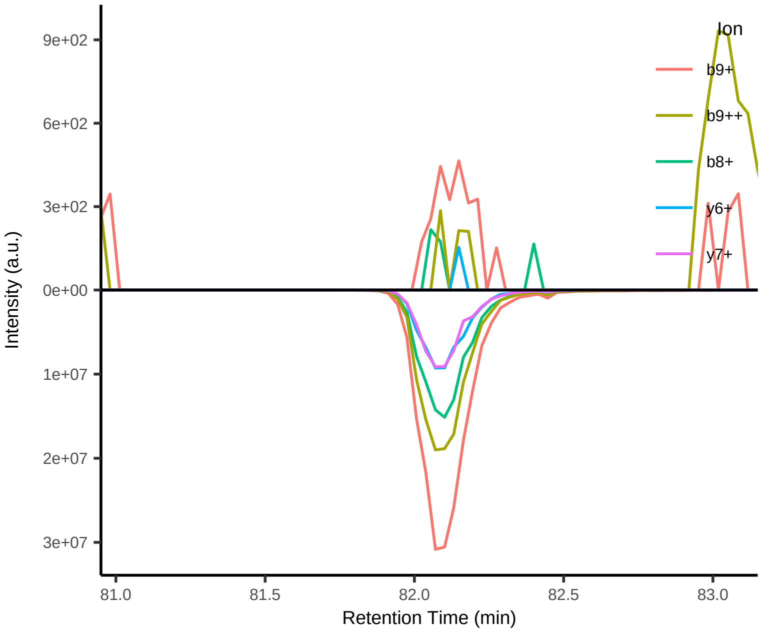


#17 ELQTTI[+7.017164]HDI++

Peptide mix
Top 4 transitions: dotp = 0.72; l:h ppm = 6

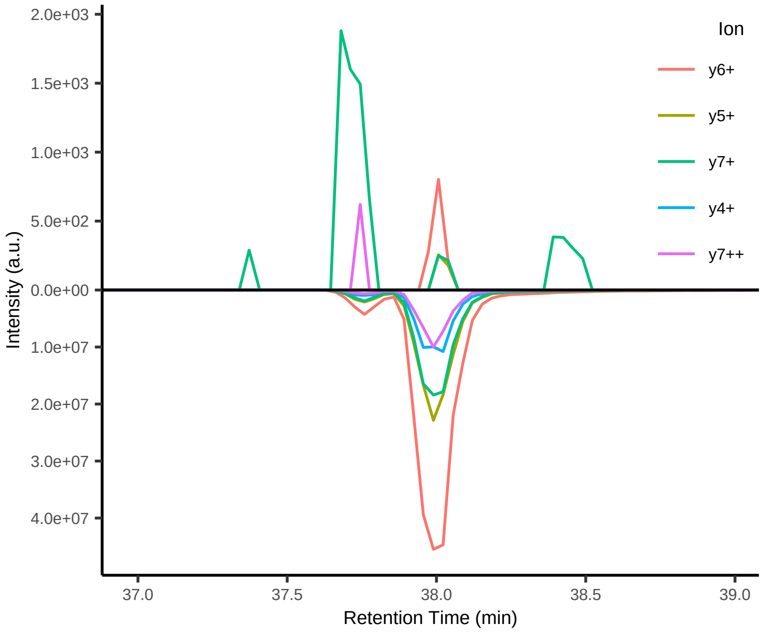

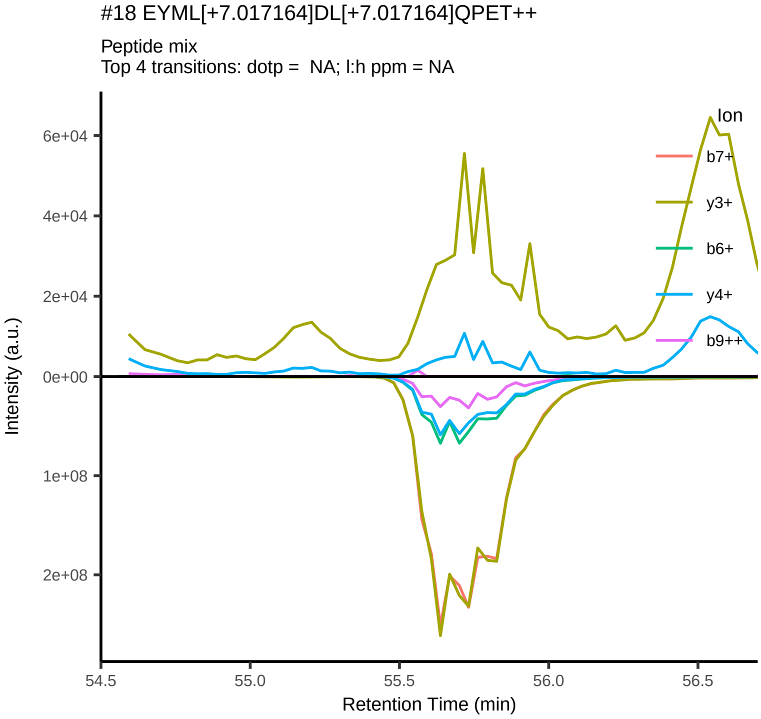
#18 EYML[+7.017164]DL[+7.017164]QPET++
Peptide mix
Top 4 transitions: dotp = NA; l:h ppm = NA
6e+04
4e+04
2e+04
0e+00
1e+08
2e+08
Intensity (a.u.)
54.5
55.0
55.5
56.0
56.5
Retention Time (min)
Ion
b7+
y3+
b6+
y4+
b9++

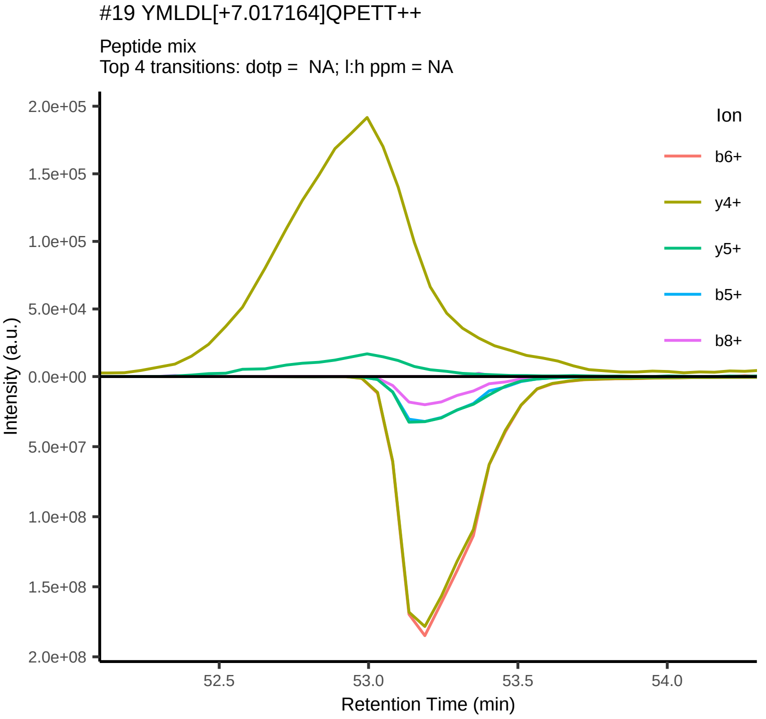
#19 YMLDL[+7.017164]QPETT++
Peptide mix
Top 4 transitions: dotp = NA; l:h ppm = NA
2.0e+05
1.5e+05
1.0e+05
5.0e+04
0.0e+00
5.0e+07
1.0e+08
1.5e+08
2.0e+08
Intensity (a.u.)
52.5
53.0
53.5
54.0
Retention Time (min)
Ion
b6+
y4+
y5+
b5+
b8+

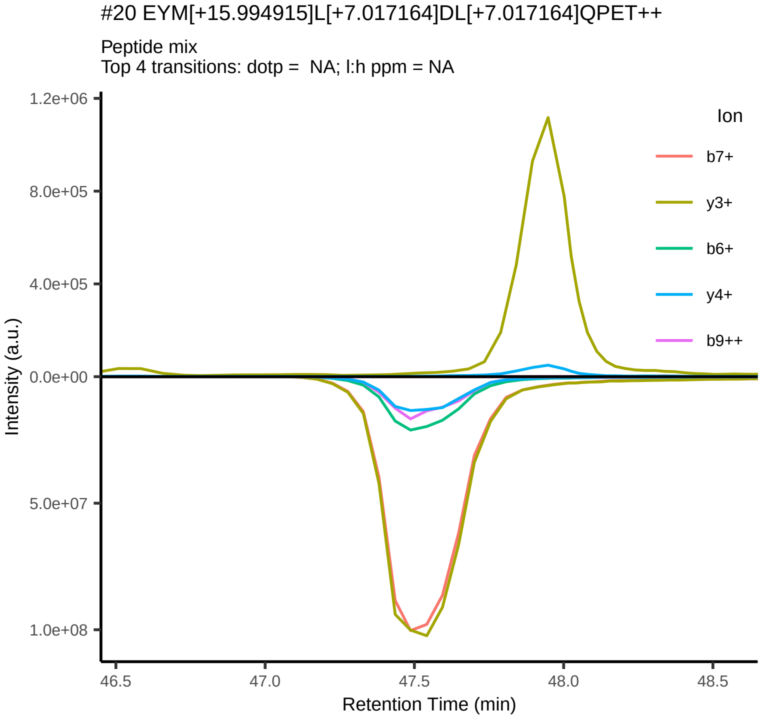
#20 EYM[+15.994915]L[+7.017164]DL[+7.017164]QPET++
Peptide mix
Top 4 transitions: dotp = NA; l:h ppm = NA
1.2e+06
8.0e+05
4.0e+05
0.0e+00
5.0e+07
1.0e+08
Intensity (a.u.)
46.5
47.0
47.5
48.0
48.5
Retention Time (min)
Ion
b7+
y3+
b6+
y4+
b9++

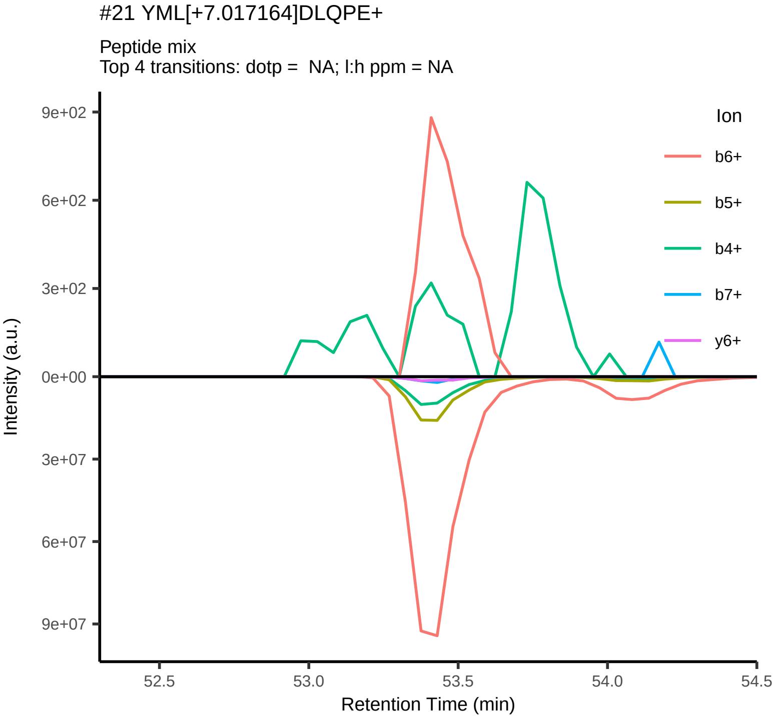
#21 YML[+7.017164]DLQPE+
Peptide mix
Top 4 transitions: dotp = NA; l:h ppm = NA
9e+02
6e+02
3e+02
0e+00
3e+07
6e+07
9e+07
Intensity (a.u.)
52.5
53.0
53.5
54.0
54.5
Retention Time (min)
Ion
b6+
b5+
b4+
b7+
y6+

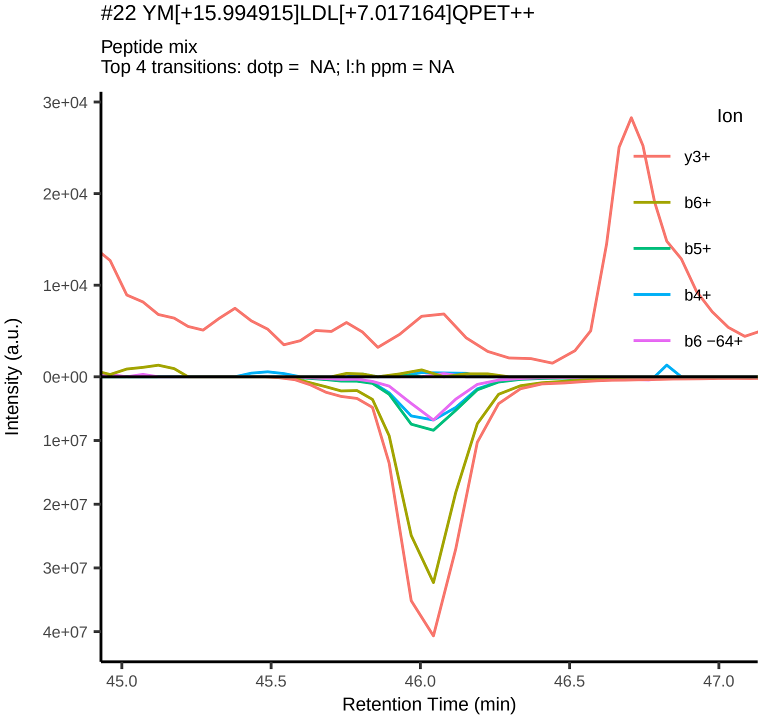
#22 YM[+15.994915]LDL[+7.017164]QPET++
Peptide mix
Top 4 transitions: dotp = NA; l:h ppm = NA
3e+04
2e+04
1e+04
0e+00
1e+07
2e+07
3e+07
4e+07
Intensity (a.u.)
45.0
45.5
46.0
46.5
47.0
Retention Time (min)
Ion
y3+
b6+
b5+
b4+
b6 −64+

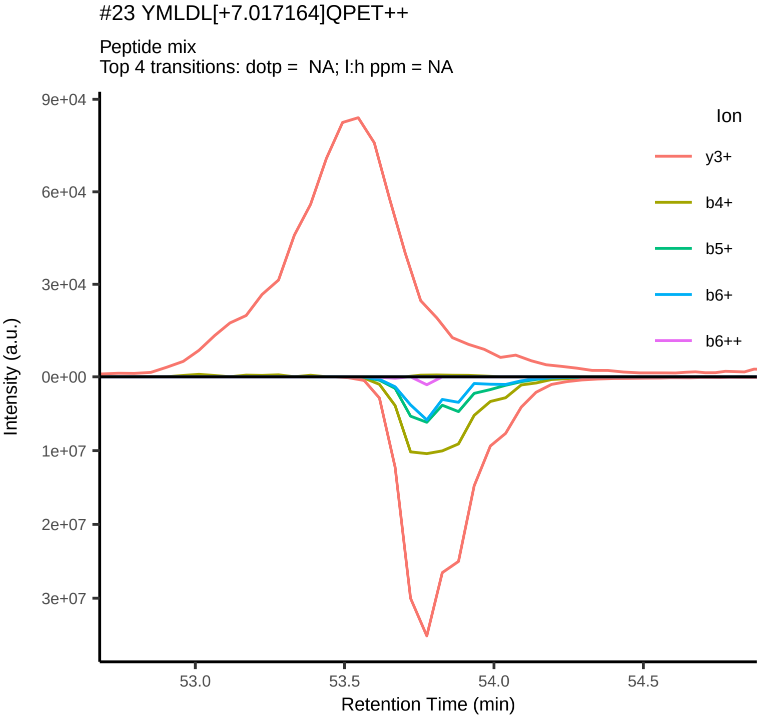
#23 YMLDL[+7.017164]QPET++
Peptide mix
Top 4 transitions: dotp = NA; l:h ppm = NA
9e+04
6e+04
3e+04
0e+00
1e+07
2e+07
3e+07
Intensity (a.u.)
53.0
53.5
54.0
54.5
Retention Time (min)
Ion
y3+
b4+
b5+
b6+
b6++

#24 MLDL[+7.017164]QPET++

Peptide mix
Top 4 transitions: dotp =  NA; l:h ppm = NA

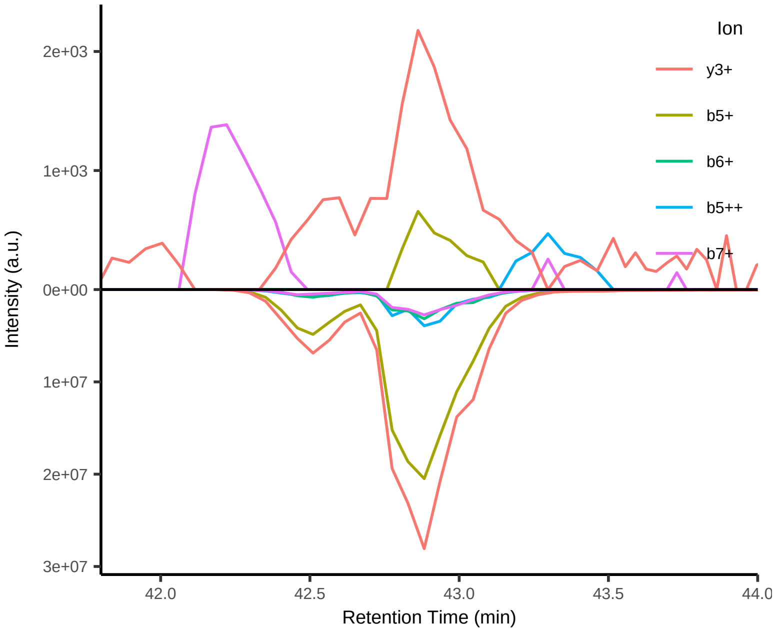


#25 LLM[+15.994915]GTLGI[+7.017164]V++

Peptide mix
Top 4 transitions: dotp =  NA; l:h ppm = NA

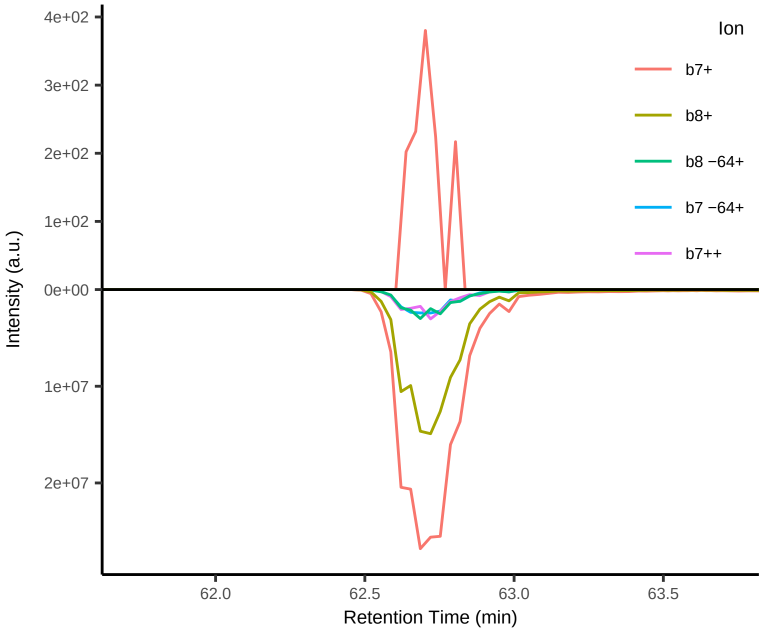


#26 EDLLMGTLGI[+7.017164]V++

Peptide mix
Top 4 transitions: dotp =  NA; l:h ppm = NA

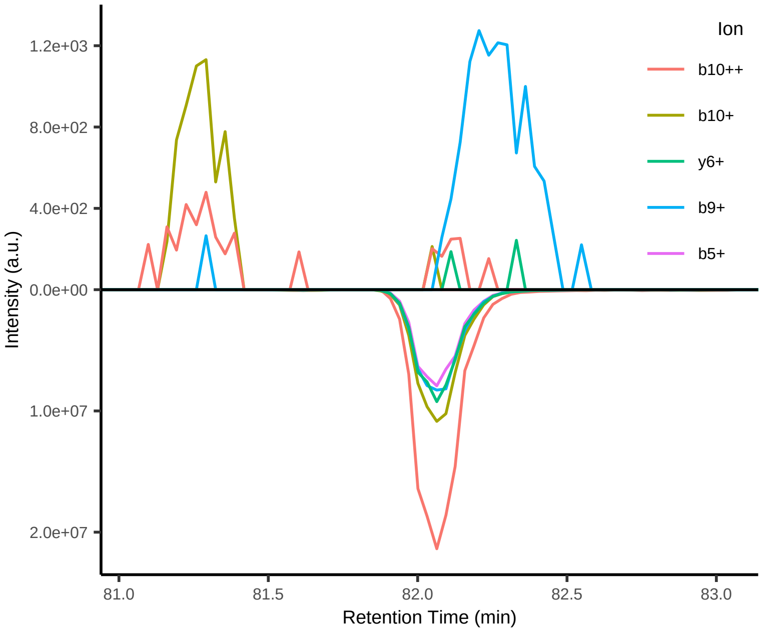


#27 DLLM[+15.994915]GTLGI[+7.017164]V++

Peptide mix
Top 4 transitions: dotp =  NA; l:h ppm = NA

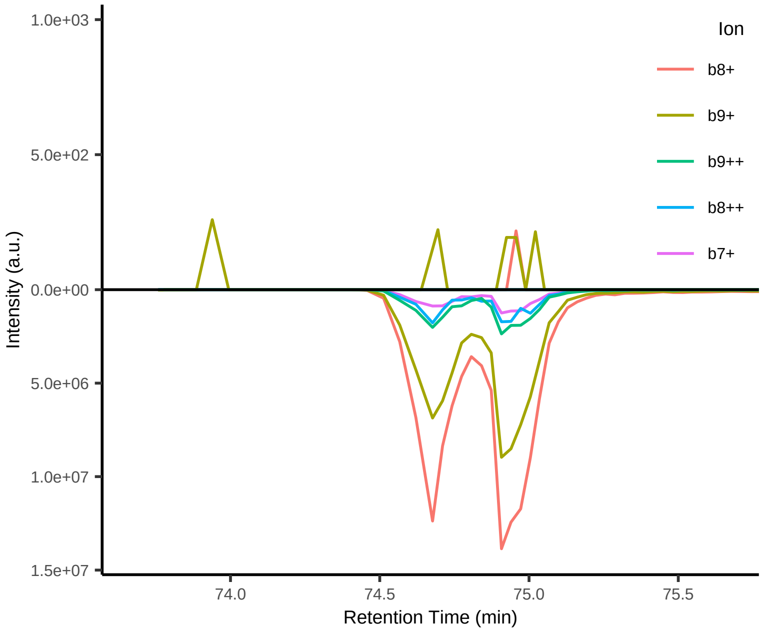


#28 MLDL[+7.017164]QPETT+

Peptide mix
Top 4 transitions: dotp =  NA; l:h ppm = NA

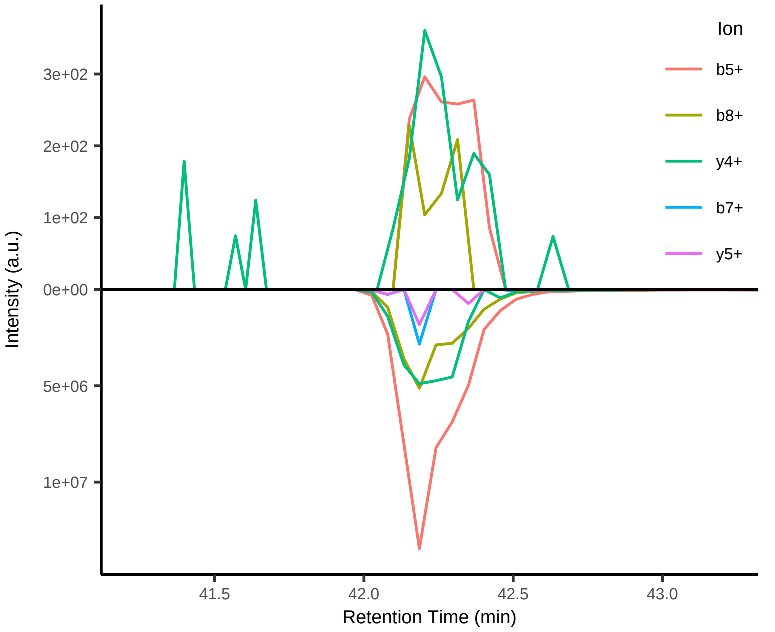


#29 EDLLM[+15.994915]GTLGI[+7.017164]V++

Peptide mix
Top 4 transitions: dotp =  NA; l:h ppm = NA

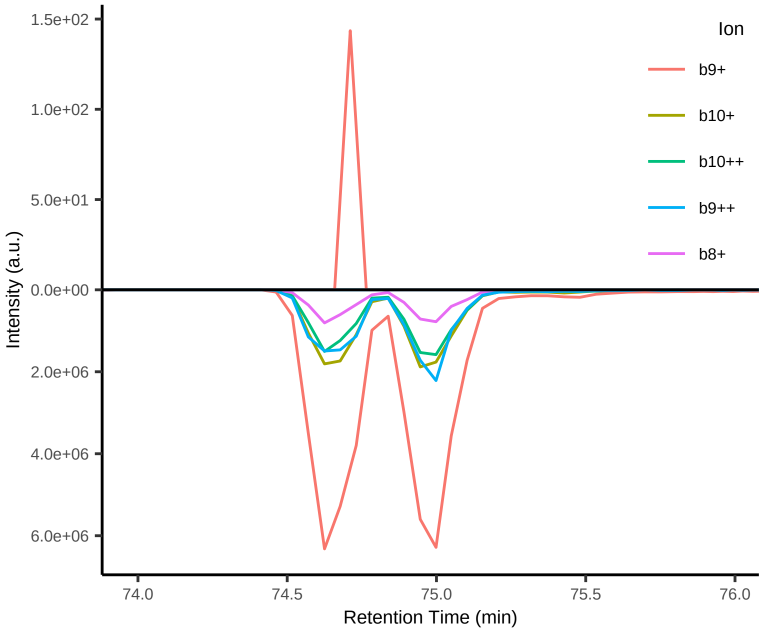

#30 M[+15.994915]LDL[+7.017164]QPET++

Peptide mix
Top 4 transitions: dotp = NA; l:h ppm = NA

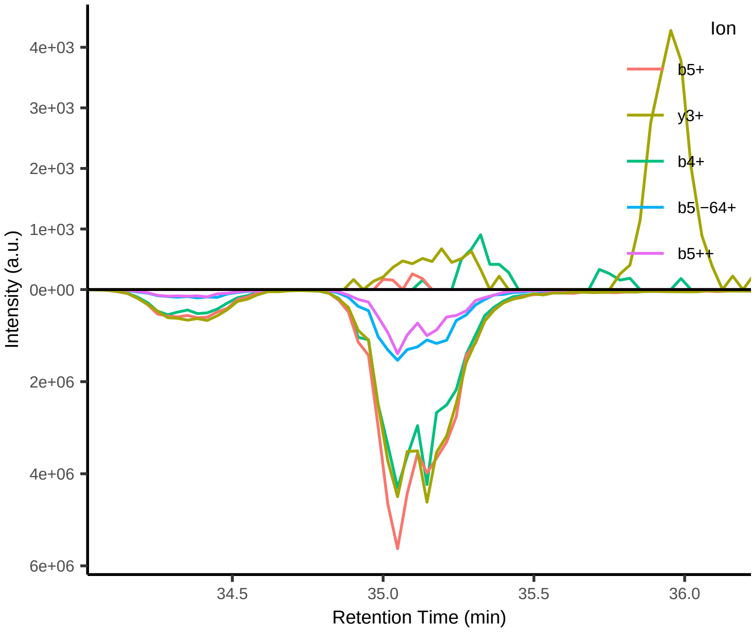


#31 YM[+15.994915]L[+7.017164]DLQPE++

Peptide mix
Top 4 transitions: dotp = NA; l:h ppm = NA

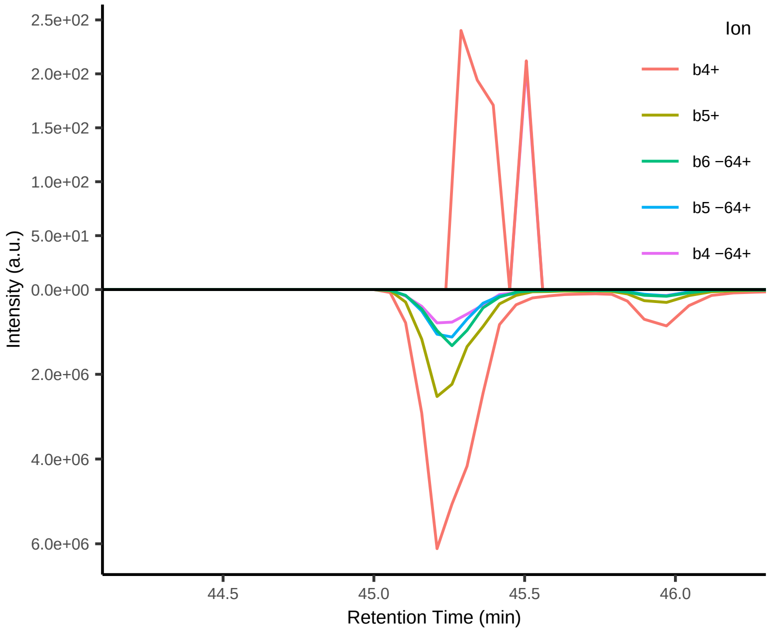


#32 TLHEYML[+7.017164]DLQP++

Peptide mix
Top 4 transitions: dotp = NA; l:h ppm = NA

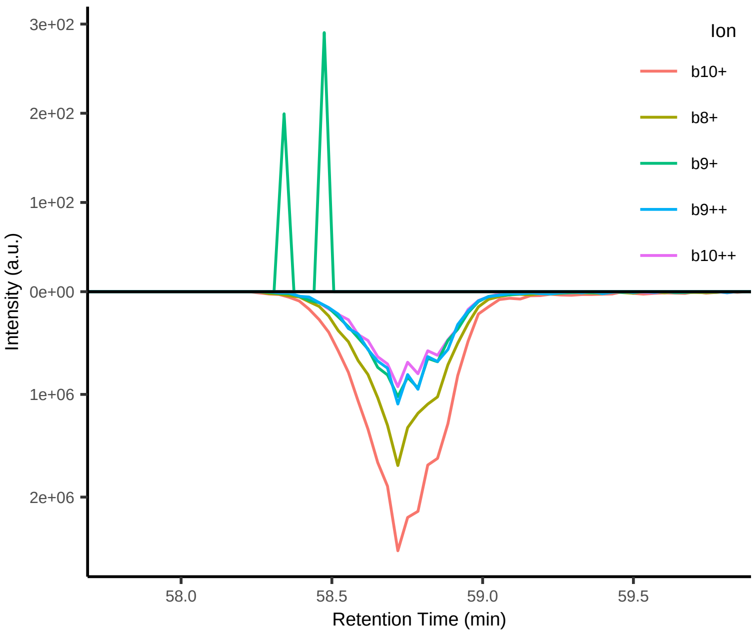


#33 TLHEYM[+15.994915]L[+7.017164]DLQP++

Peptide mix
Top 4 transitions: dotp = NA; l:h ppm = NA

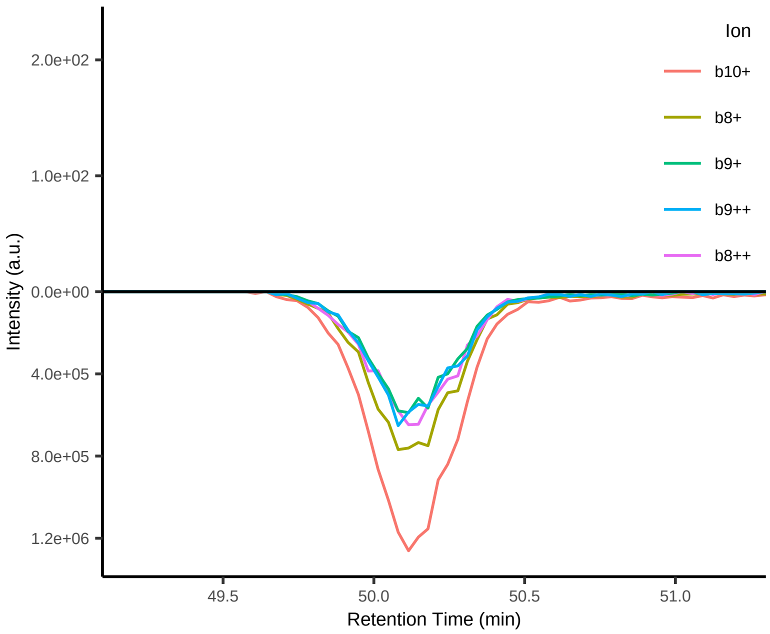


#34 AAL[+7.017164]ENTHLL++

Peptide mix
Top 4 transitions: dotp = NA; l:h ppm = NA

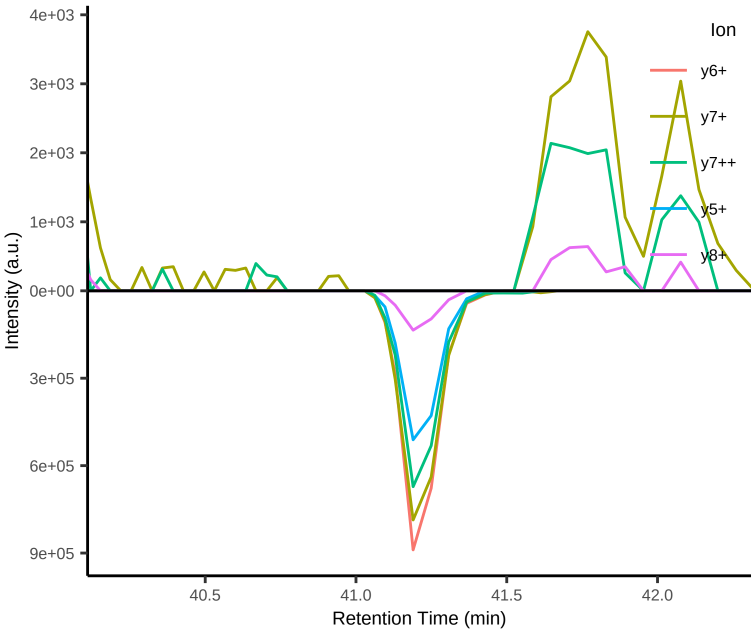


#35 TL[+7.017164]HEYMLDL++

Peptide mix
Top 4 transitions: dotp = NA; l:h ppm = NA

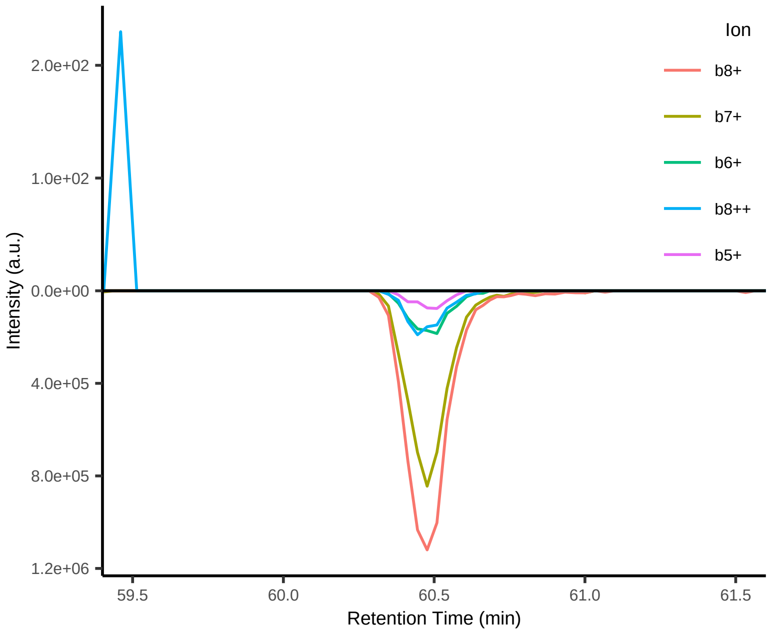

#36 FGPV[+6.013809]NHEEL++

Peptide mix
Top 4 transitions: dotp = NA; l:h ppm = NA

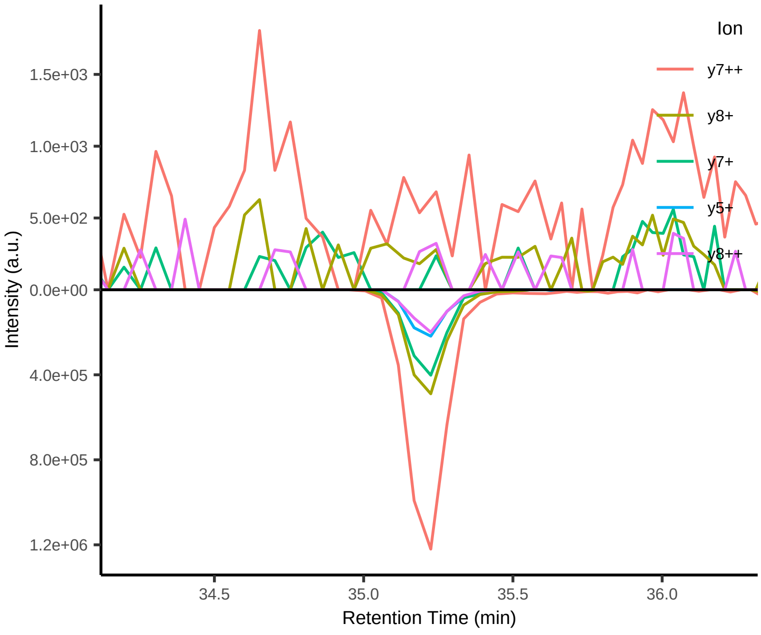


#37 AL[+7.017164]NEKLVNL++

Peptide mix
Top 4 transitions: dotp = NA; l:h ppm = NA

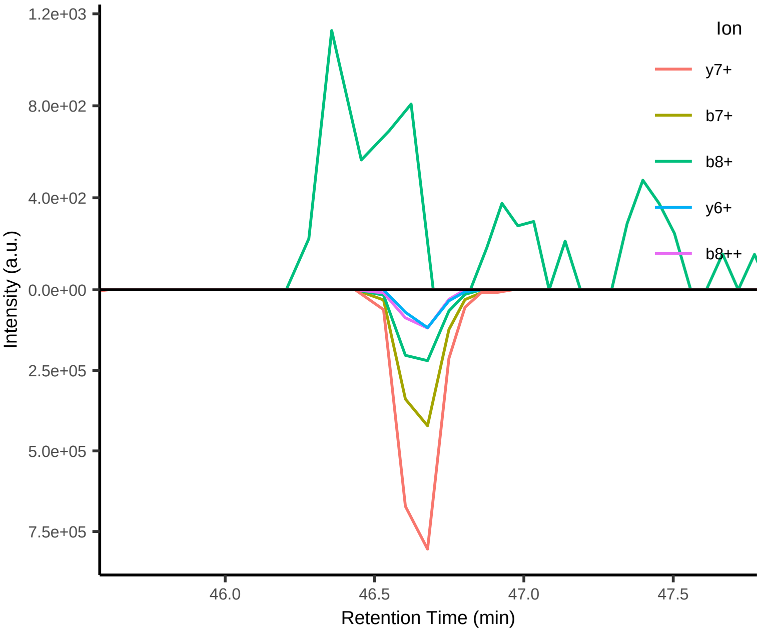


#38 KAL[+7.017164]INADEL++

Peptide mix
Top 4 transitions: dotp = NA; l:h ppm = NA

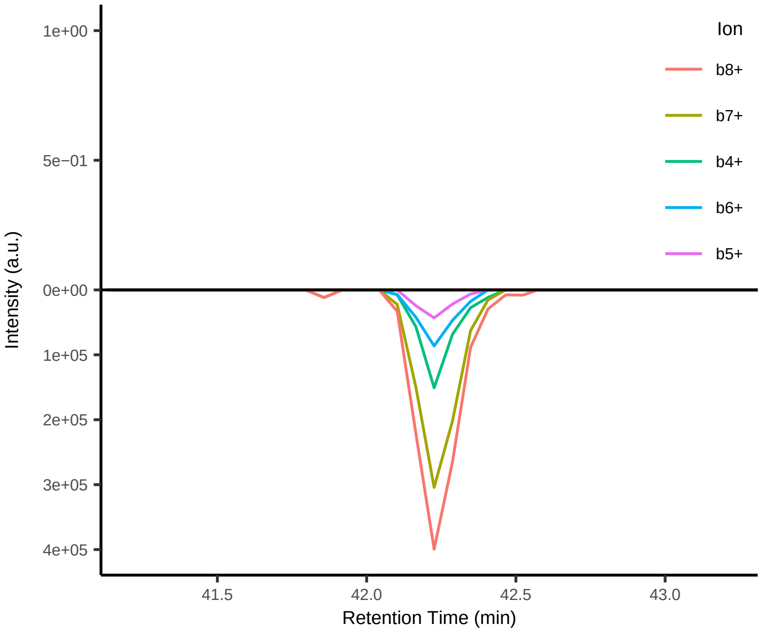


#39 TL[+7.017164]HEYM[+15.994915]LDL++

Peptide mix
Top 4 transitions: dotp = NA; l:h ppm = NA

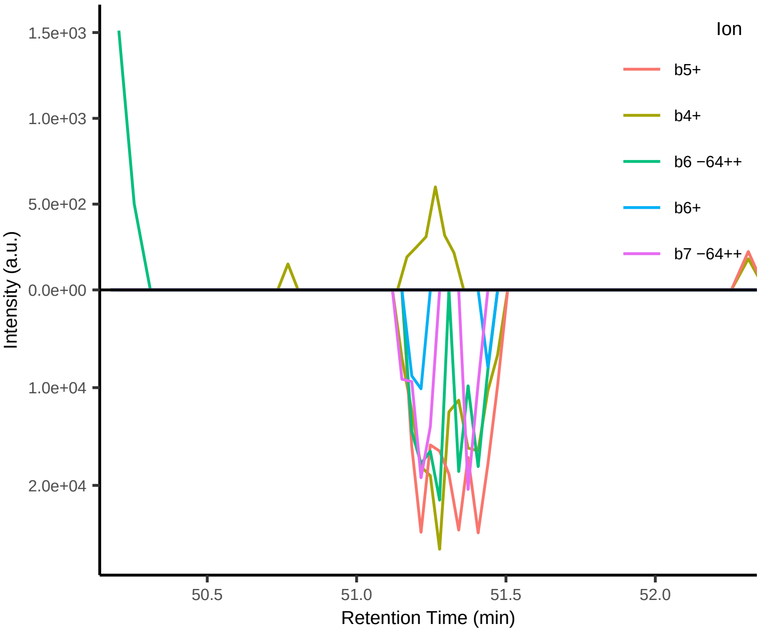


#40 TASEFDSAIAQDK[+8.014199]++

Peptide mix
Top 4 transitions: dotp = 0.99; l:h ppm = 44962

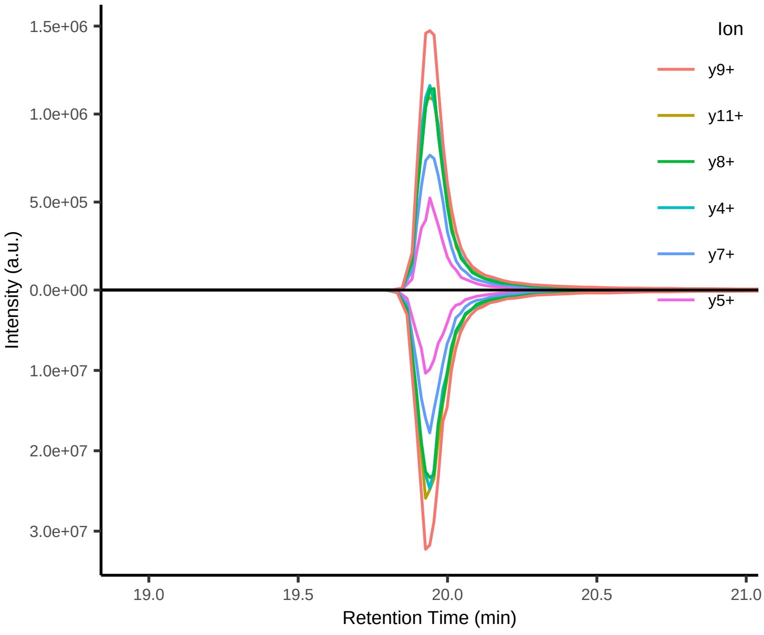


#41 DIPVPKPK[+8.014199]++

Peptide mix
Top 4 transitions: dotp = 0.99; l:h ppm = 835

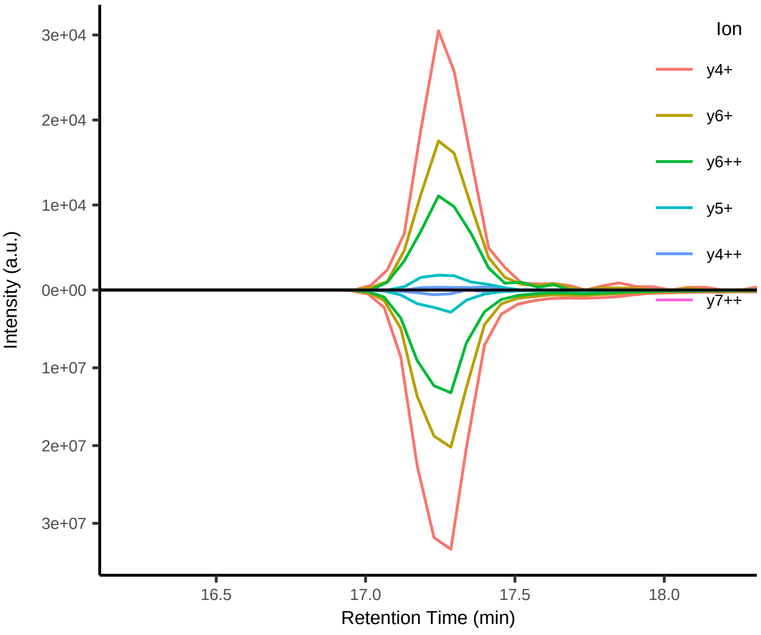

#42 ELGQSGVDTYLQTK[+8.014199]++

Peptide mix
Top 4 transitions: dotp = 0.97; l:h ppm = 285

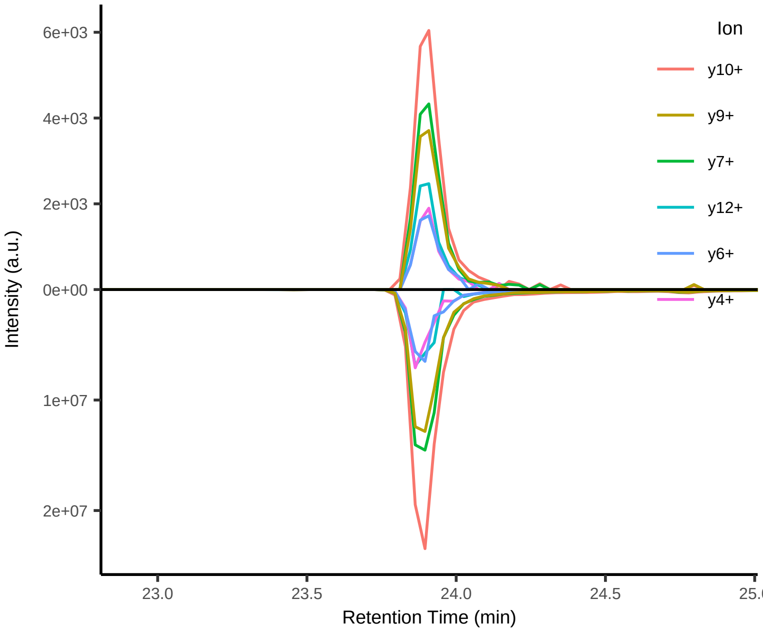


#43 GISNEGQNASIK[+8.014199]++

Peptide mix
Top 4 transitions: dotp = 0.98; l:h ppm = 221

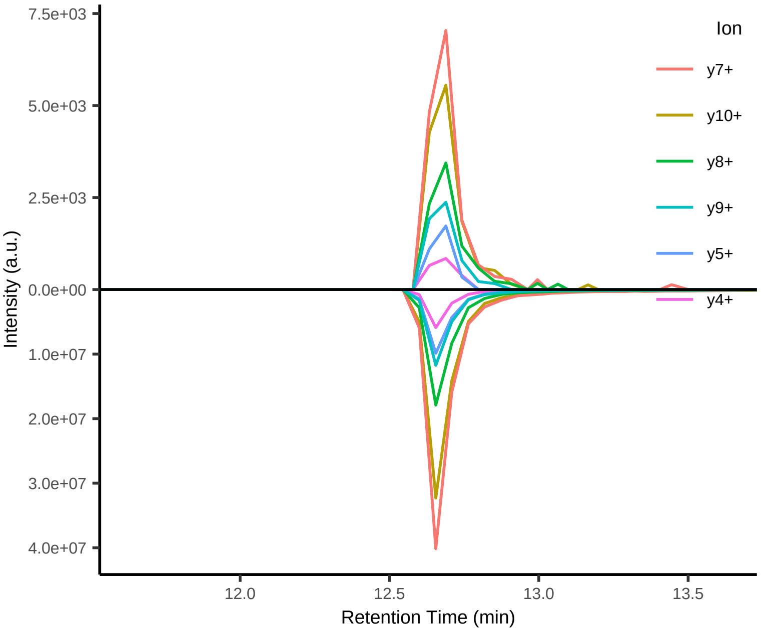


#44 NGFILDGFPR[+10.008269]++

Peptide mix
Top 4 transitions: dotp = 0.97; l:h ppm = 202

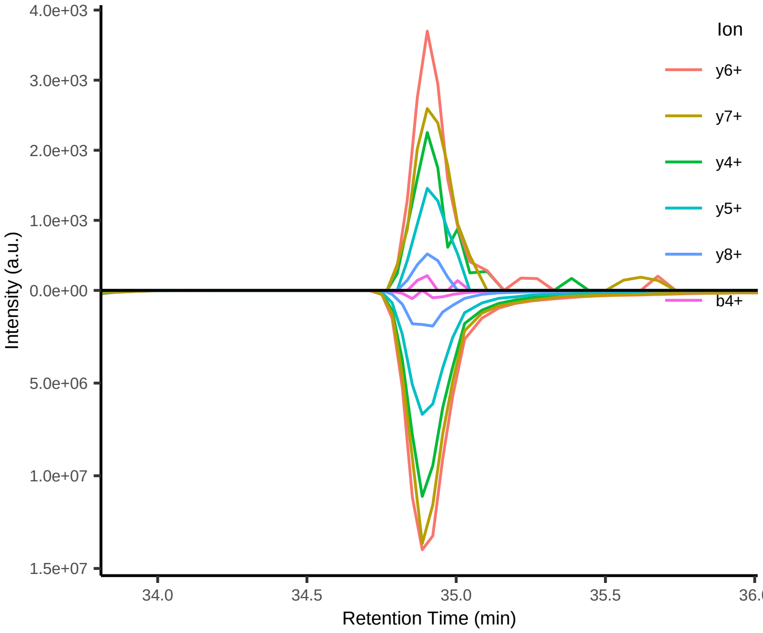


#45 HVLTSIGEK[+8.014199]++

Peptide mix
Top 4 transitions: dotp = 0.98; l:h ppm = 173

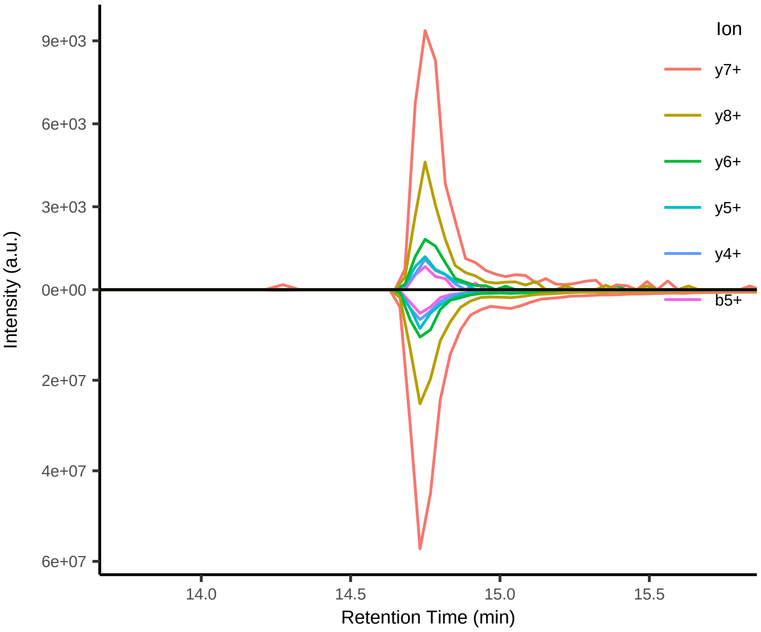


#46 IGDYAGIK[+8.014199]++

Peptide mix
Top 4 transitions: dotp = 0.98; l:h ppm = 150

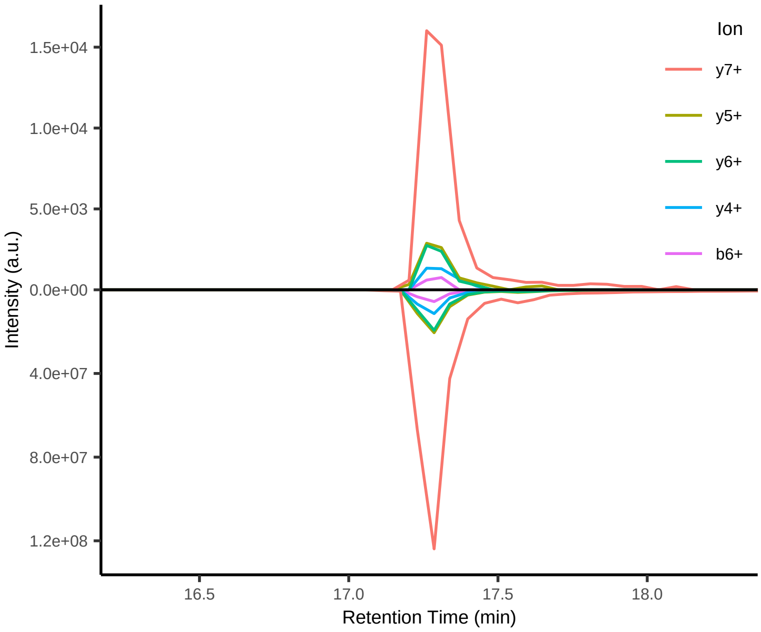


#47 SFANQPLEVVYSK[+8.014199]++

Peptide mix
Top 4 transitions: dotp = 0.94; l:h ppm = 145

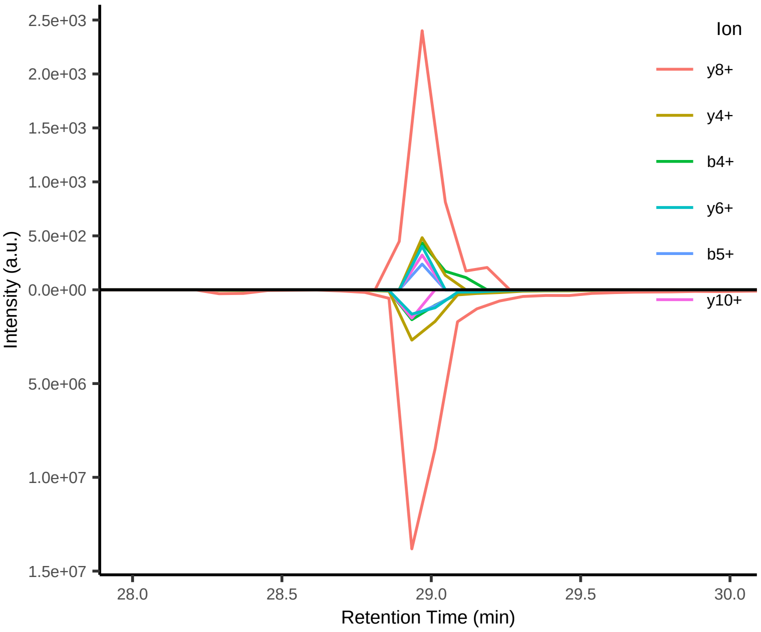

#48 LSSEAPALFQFDLK[+8.014199]++

Peptide mix
Top 4 transitions: dotp = 0.54; l:h ppm = 142

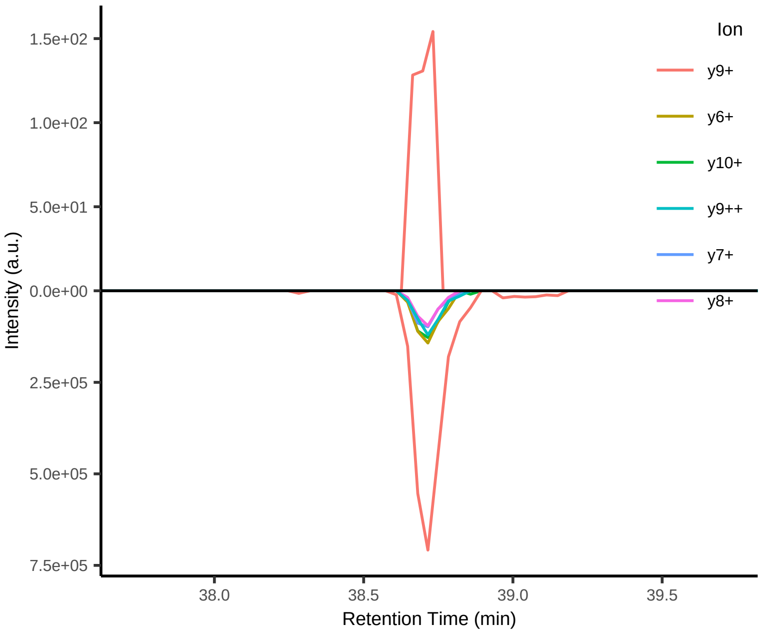


#49 SAAGAFGPELSR[+10.008269]++

Peptide mix
Top 4 transitions: dotp = 0.99; l:h ppm = 106

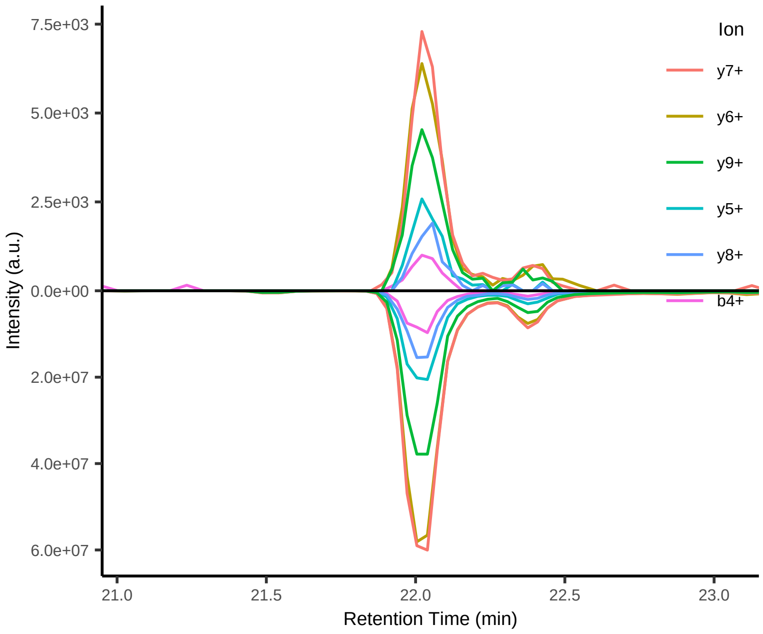


#50 ELASGLSFPVGFK[+8.014199]++

Peptide mix
Top 4 transitions: dotp = 0.80; l:h ppm = 104

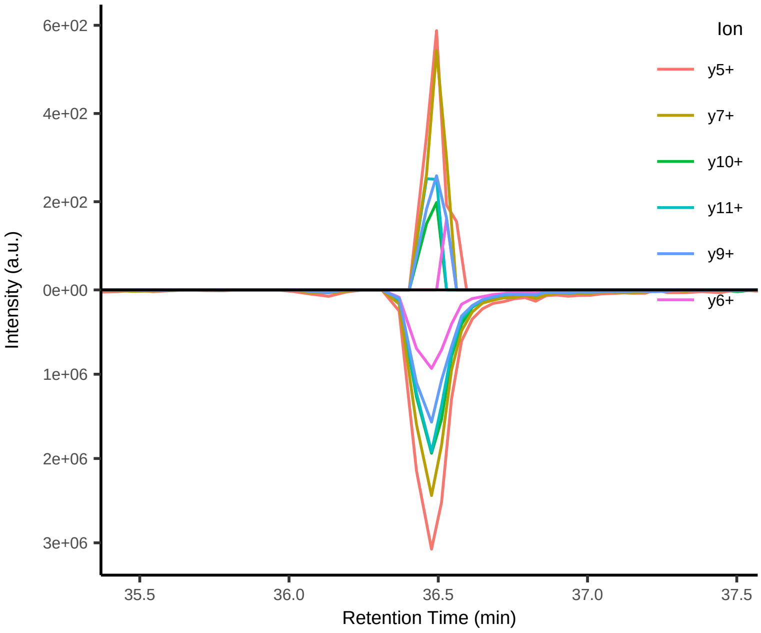


#51 GLILVGGYGTR[+10.008269]++

Peptide mix
Top 4 transitions: dotp = 0.86; l:h ppm = 95

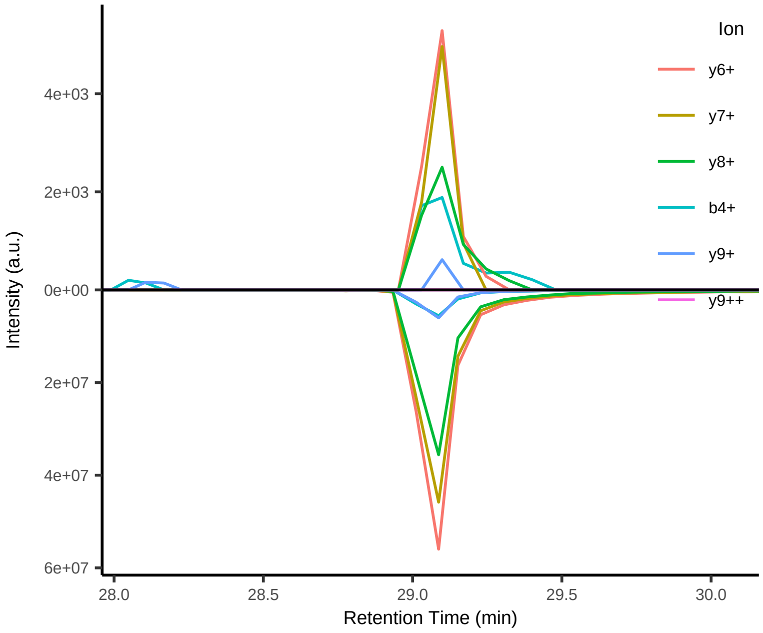


#52 GILFVGSGVSGGEEGAR[+10.008269]++

Peptide mix
Top 4 transitions: dotp = 0.95; l:h ppm = 89

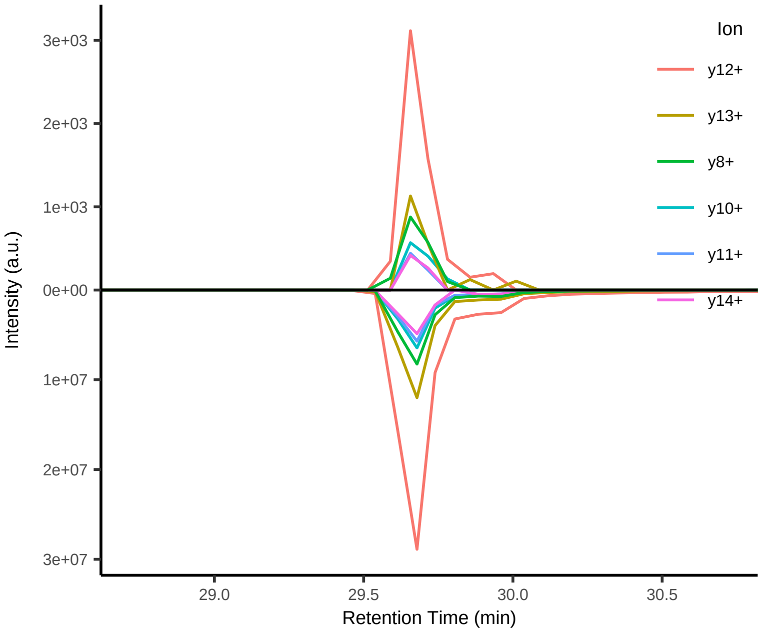


#53 LTILEELR[+10.008269]++

Peptide mix
Top 4 transitions: dotp = 0.90; l:h ppm = 25

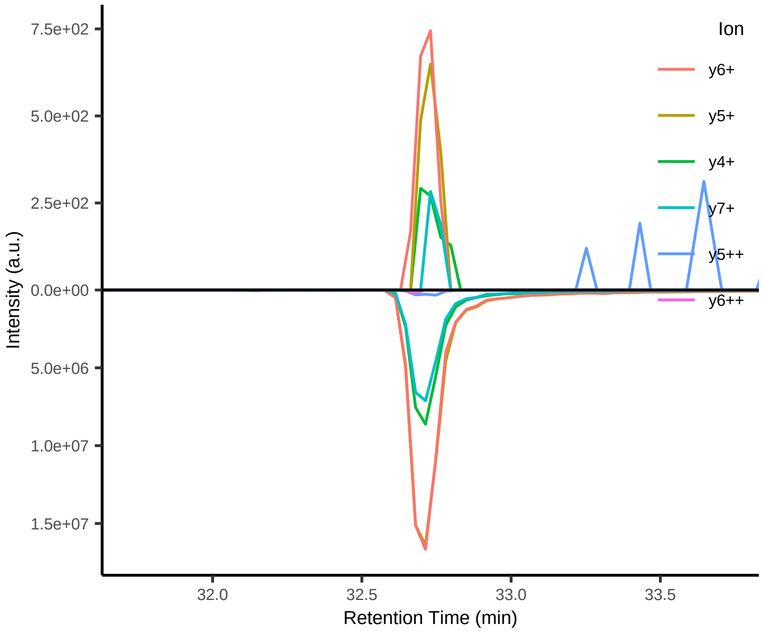

#54 SSAAPPPPPR[+10.008269]++

Peptide mix
Top 4 transitions: dotp = 0.93; l:h ppm = 25

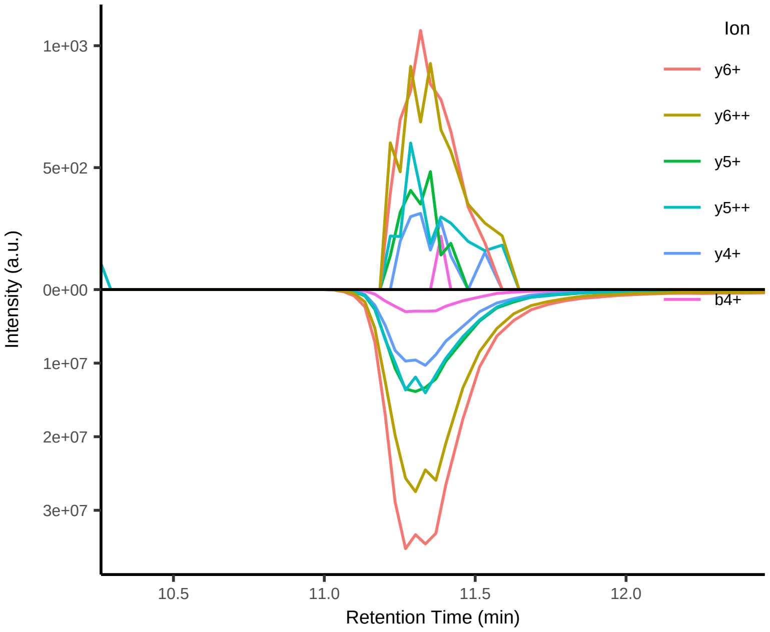


#55 IAESL[+7.017164]PVV+

Peptide mix
Top 4 transitions: dotp = 0.86; l:h ppm = 611

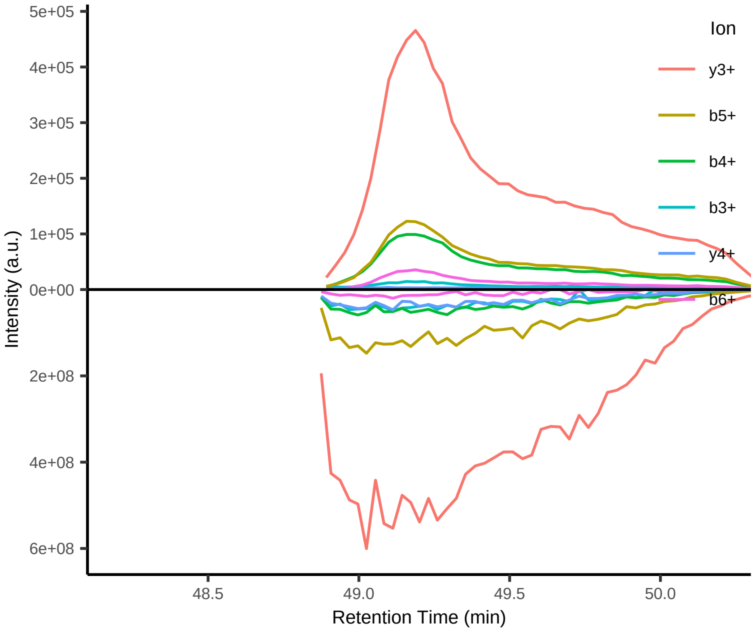


#56 SRFTGATII[+7.017164]++

Peptide mix
Top 4 transitions: dotp = 0.96; l:h ppm = 538

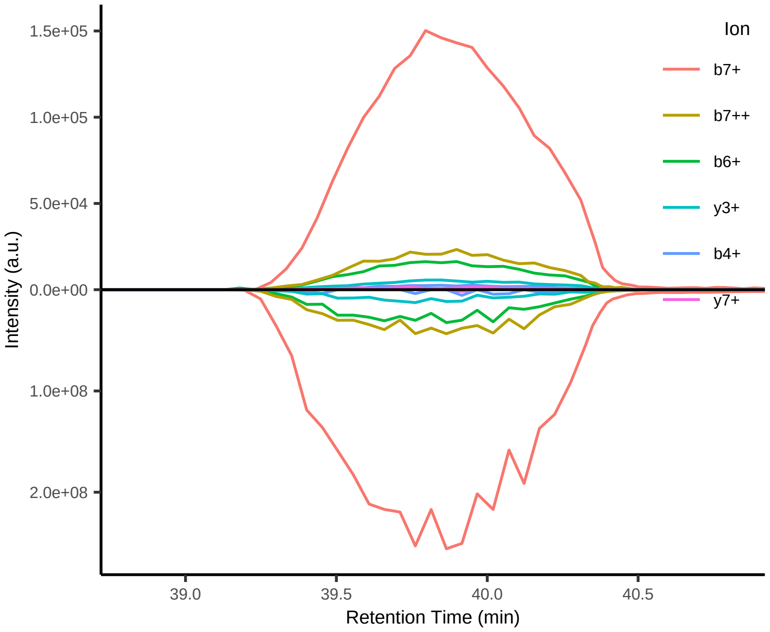


#57 KSTL[+7.017164]VRLLF++

Peptide mix
Top 4 transitions: dotp = 0.99; l:h ppm = 475

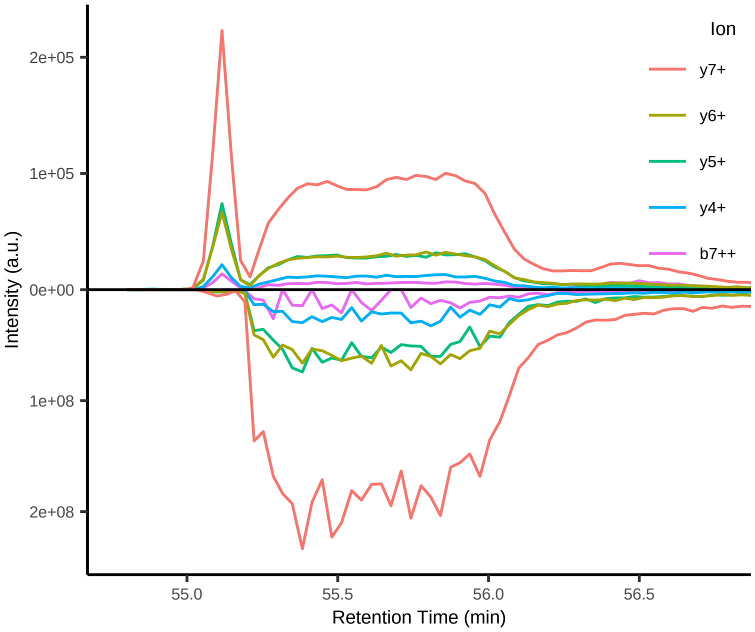


#58 L[+7.017164]NPNDPYHTYY++

Peptide mix
Top 4 transitions: dotp = 0.97; l:h ppm = 405

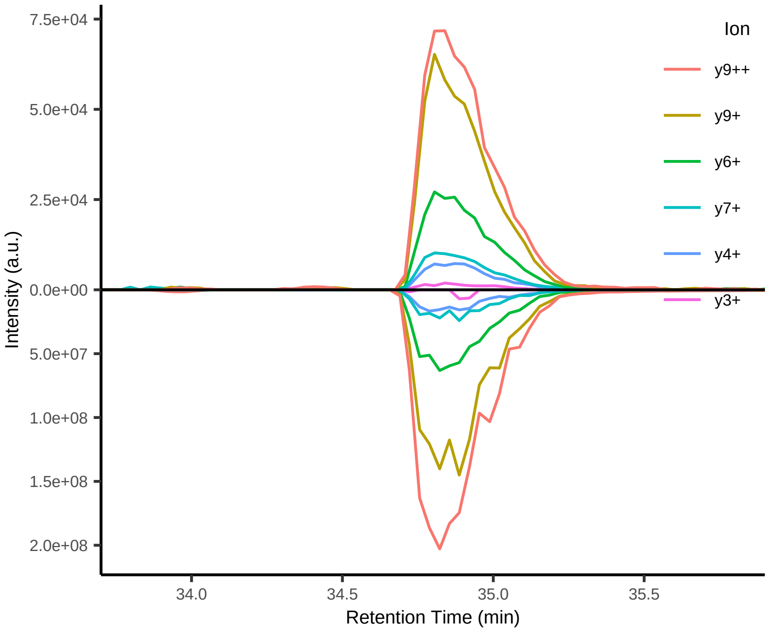


#59 HTYYRHK[+8.014199]V++

Peptide mix
Top 4 transitions: dotp = 0.96; l:h ppm = 381

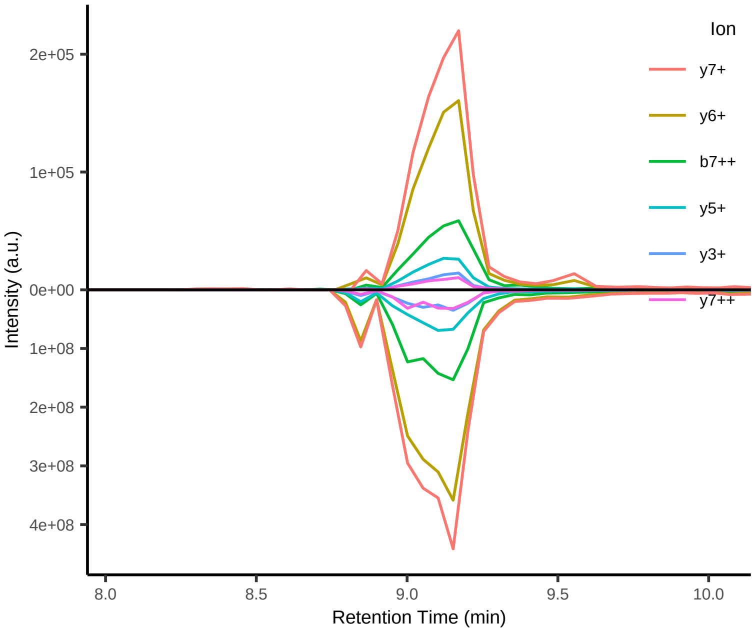

#60 SRL[+7.017164]SKEELI++

Peptide mix
Top 4 transitions: dotp = 0.99; l:h ppm = 331

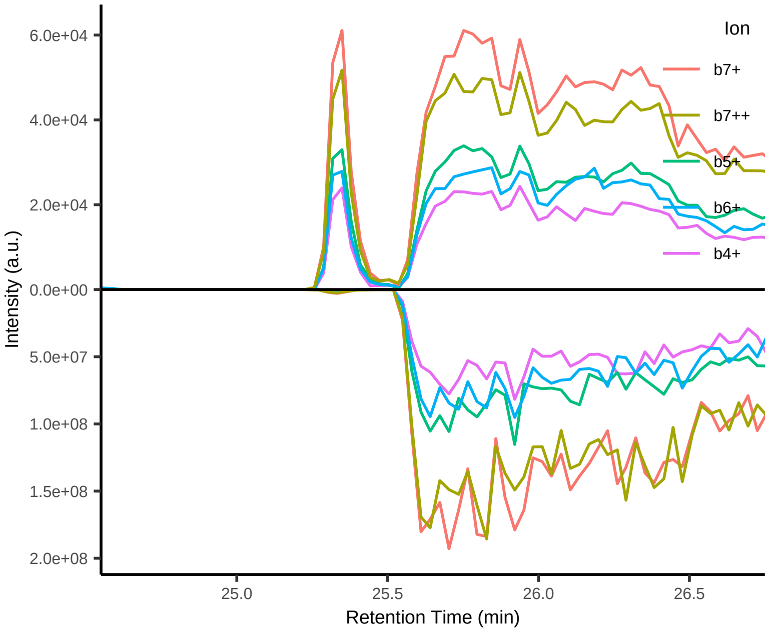


#61 M[+15.994915]LWLPHWGL[+7.017164]++

Peptide mix
Top 4 transitions: dotp = 0.94; l:h ppm = 310

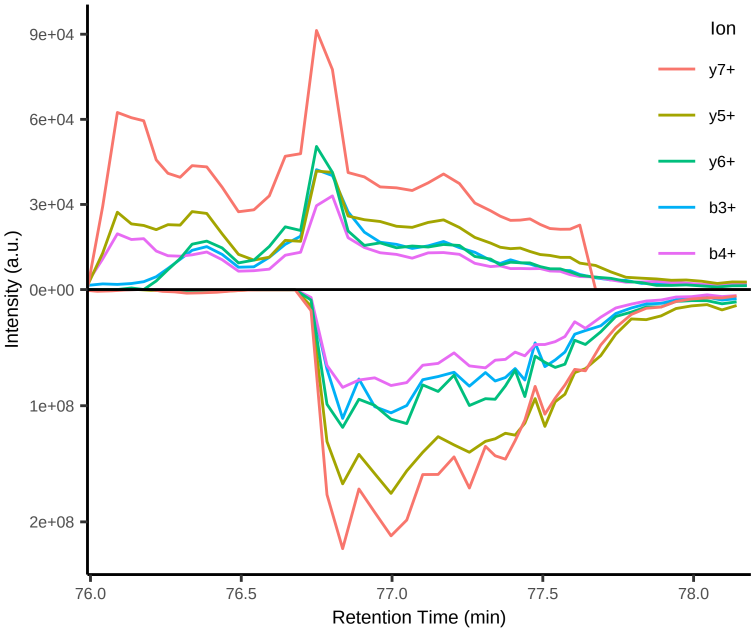


#62 MLWLPHWGL[+7.017164]++

Peptide mix
Top 4 transitions: dotp = 0.96; l:h ppm = 296

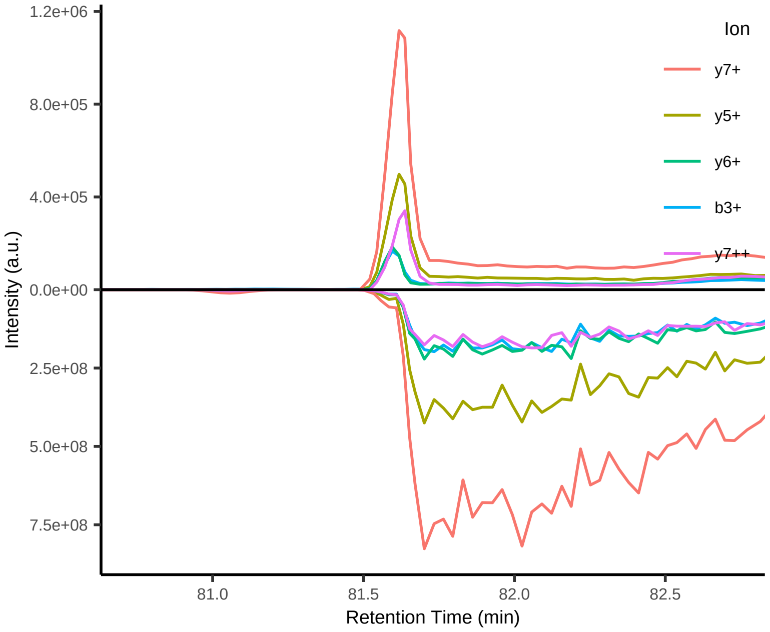


#63 RVRKELQEL[+7.017164]+++

Peptide mix
Top 4 transitions: dotp = 0.56; l:h ppm = 239

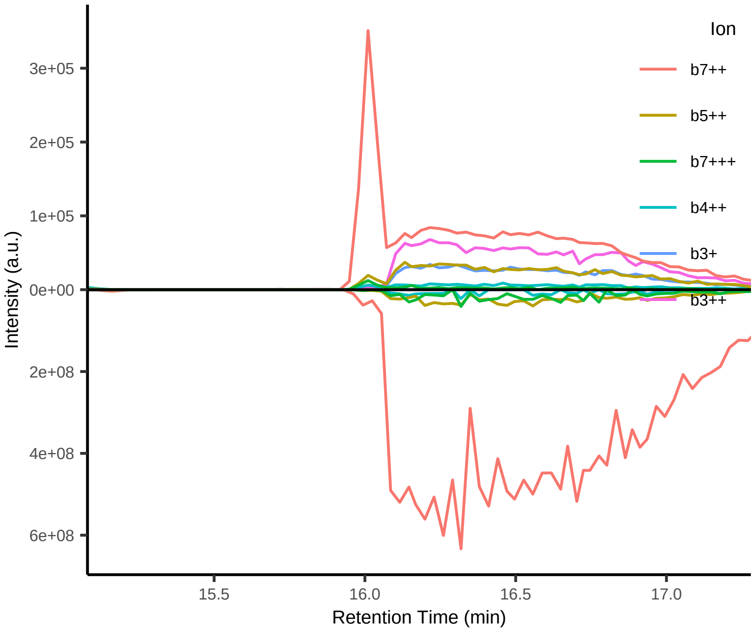


#64 KVAKGHMK[+8.014199]L++

Peptide mix
Top 4 transitions: dotp = 0.95; l:h ppm = 235

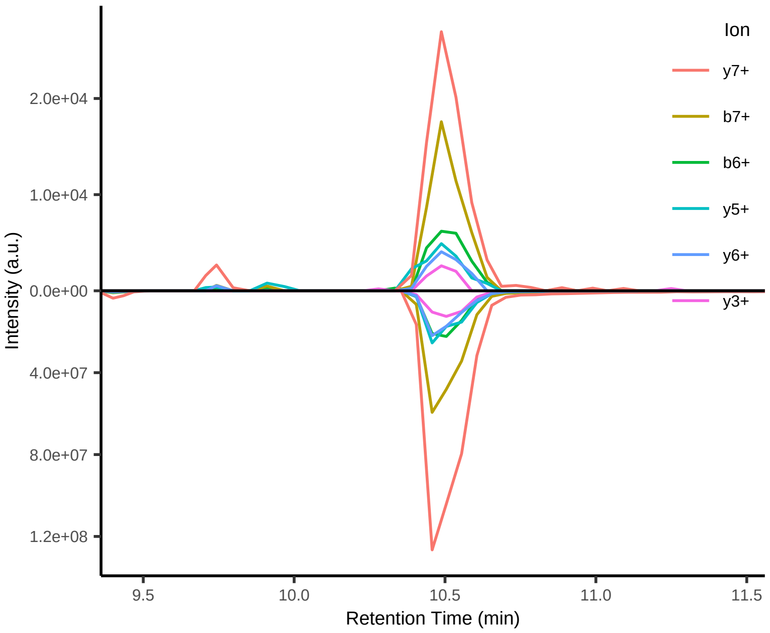


#65 FLM[+15.994915]LWL[+7.017164]PHW++

Peptide mix
Top 4 transitions: dotp = 0.87; l:h ppm = 208

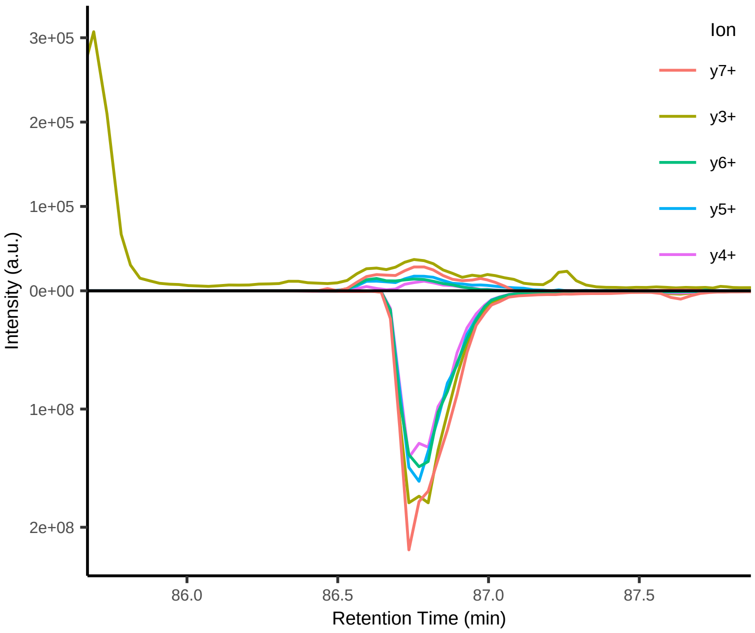

#66 YHTYYRHK[+8.014199]V++

Peptide mix
Top 4 transitions: dotp = 0.98; l:h ppm = 196

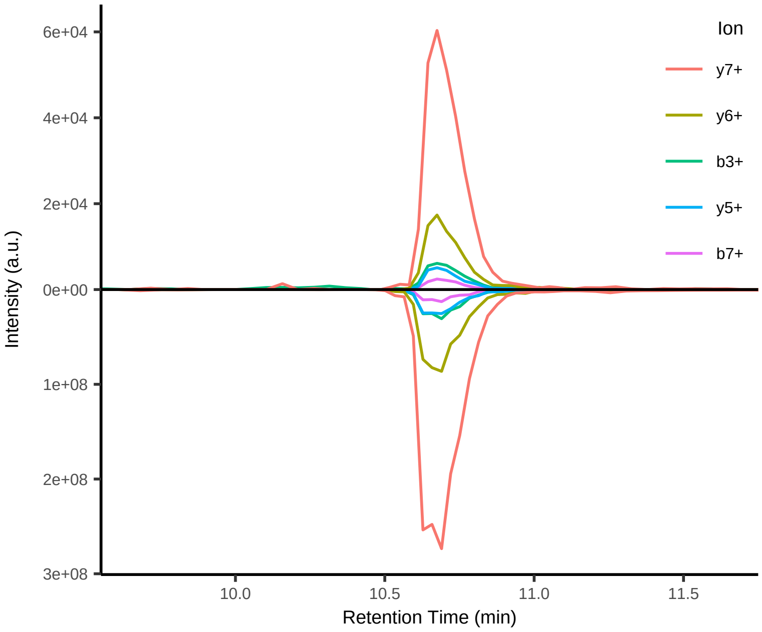


#67 GKSTL[+7.017164]VRLLF+++

Peptide mix
Top 4 transitions: dotp = 0.98; l:h ppm = 189

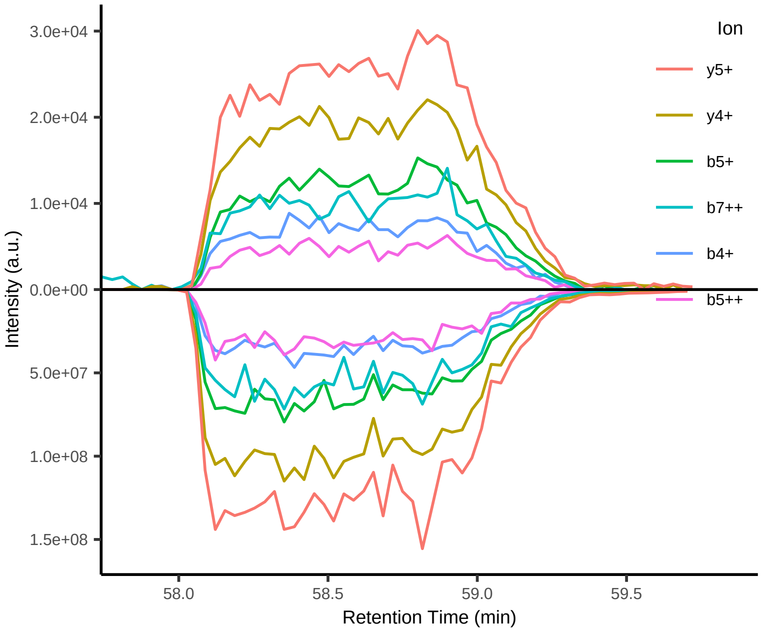


#68 RPMV[+6.013809]RSFTM[+15.994915]+++

Peptide mix
Top 4 transitions: dotp = 0.74; l:h ppm = 188

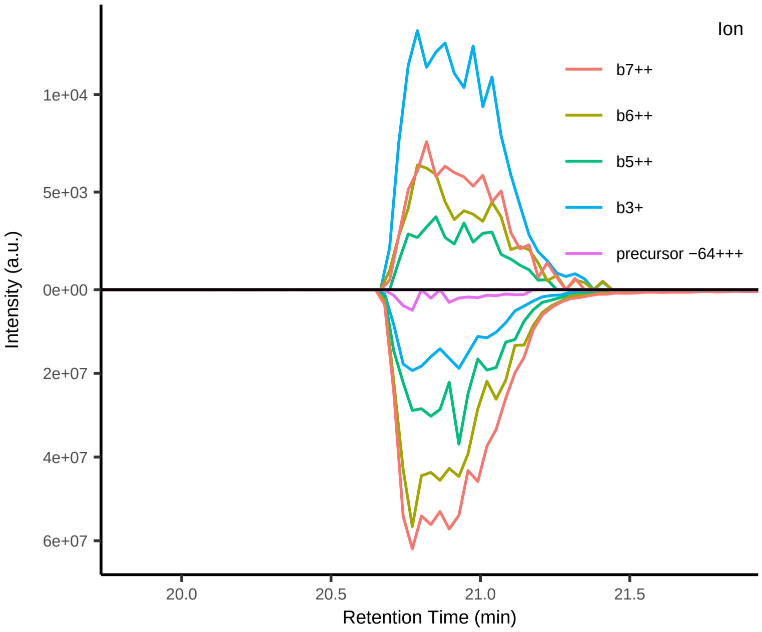


#69 KVAKGHM[+15.994915]K[+8.014199]L+++

Peptide mix
Top 4 transitions: dotp = 0.96; l:h ppm = 178

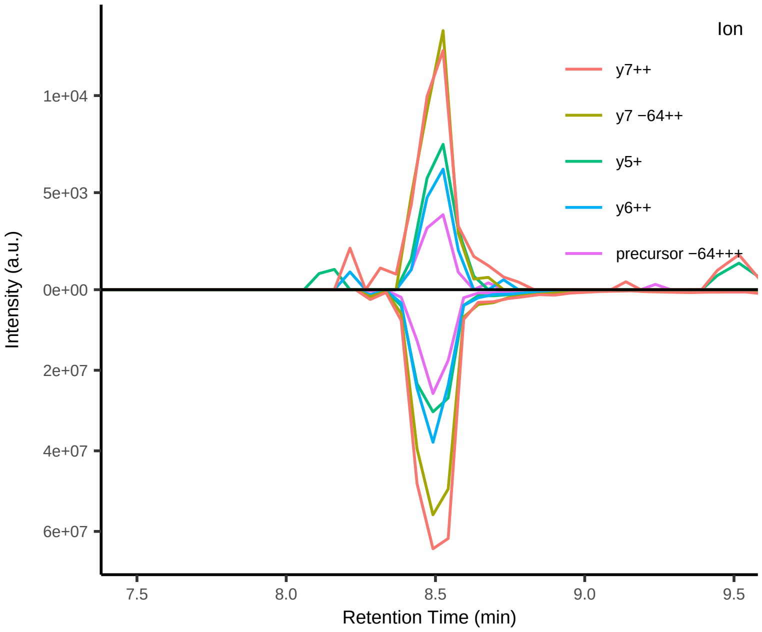


#70 KSTL[+7.017164]VRLL++

Peptide mix
Top 4 transitions: dotp = 0.96; l:h ppm = 178

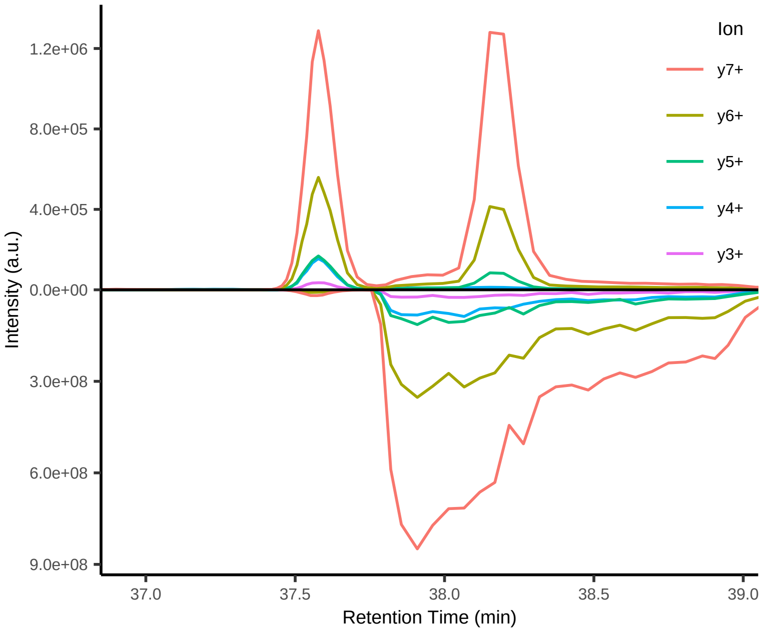


#71 RPMV[+6.013809]RSFTM+++

Peptide mix
Top 4 transitions: dotp = 0.82; l:h ppm = 177

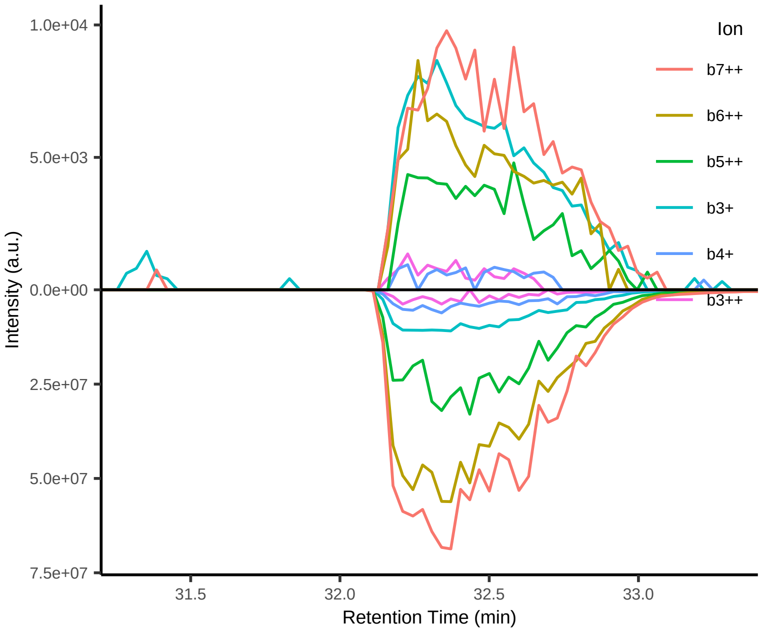

#72 MATELGIVL[+7.017164]+

Peptide mix
Top 4 transitions: dotp = 0.86; l:h ppm = 174

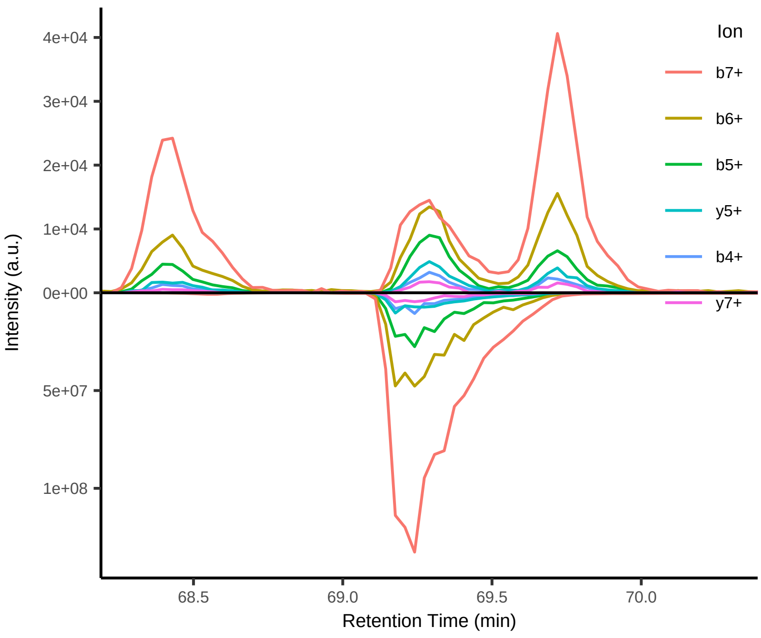


#73 FGTTV[+6.013809]PFTSW++

Peptide mix
Top 4 transitions: dotp = 0.92; l:h ppm = 170

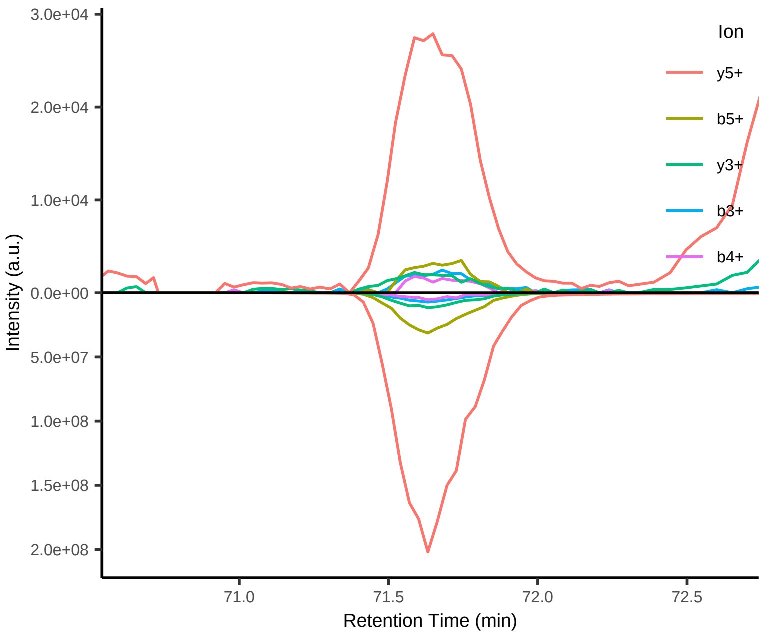


#74 FL[+7.017164]NPNDPYHTY++

Peptide mix
Top 4 transitions: dotp = 0.98; l:h ppm = 165

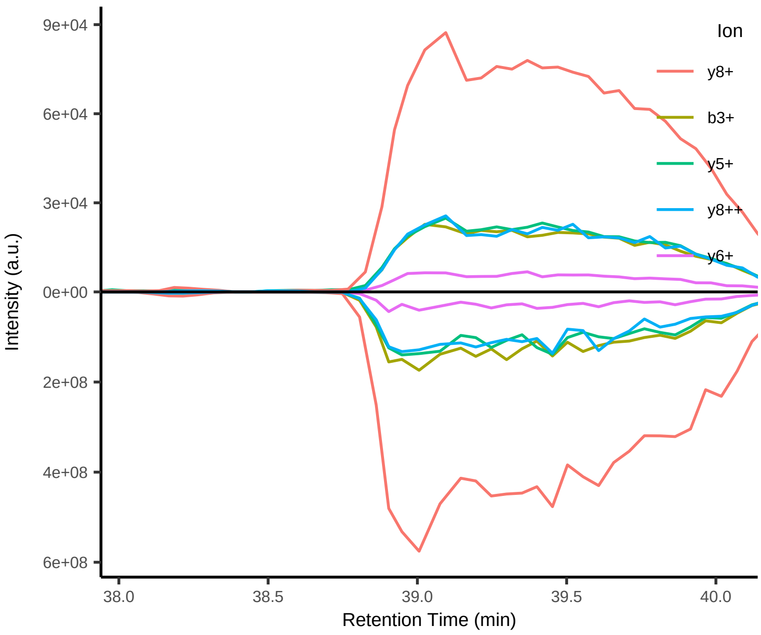


#75 TTV[+6.013809]PFTSW+

Peptide mix
Top 4 transitions: dotp = 0.96; l:h ppm = 161

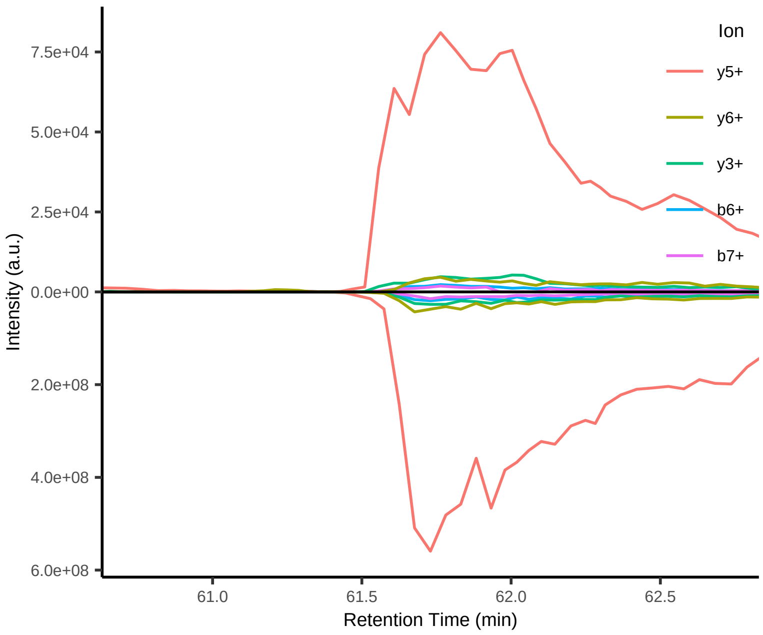


#76 GTTV[+6.013809]PFTSW++

Peptide mix
Top 4 transitions: dotp = 0.93; l:h ppm = 160

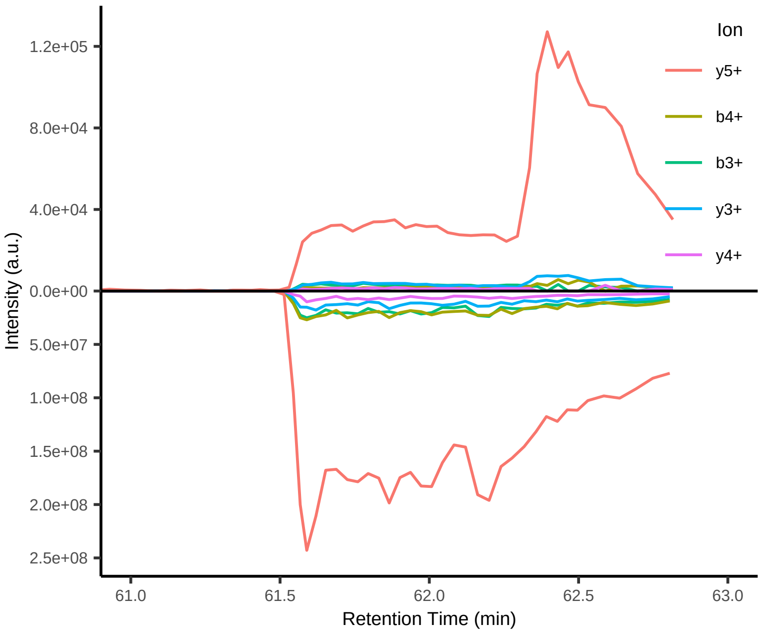


#77 SV[+6.013809]LTAFLM[+15.994915]L++

Peptide mix
Top 4 transitions: dotp = 0.99; l:h ppm = 146

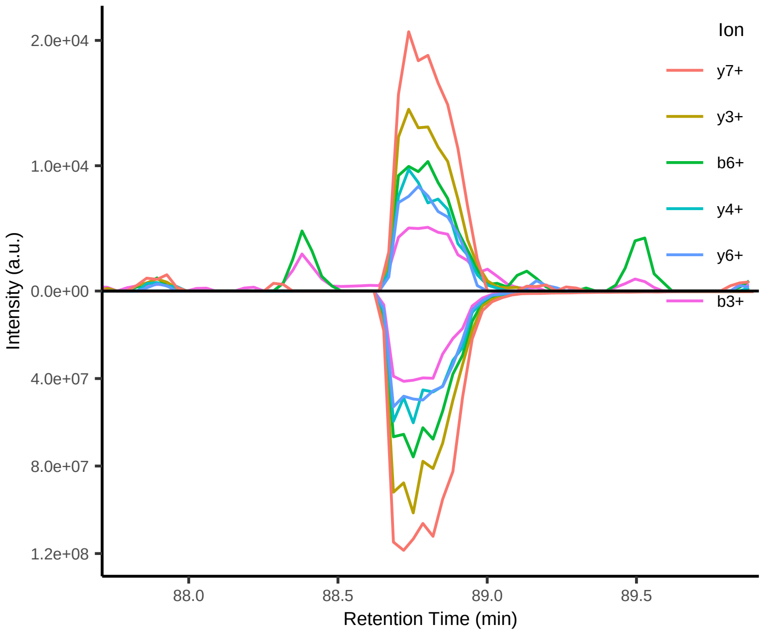

#78 RFGTTV[+6.013809]PFTSW++

Peptide mix
Top 4 transitions: dotp = 0.88; l:h ppm = 144

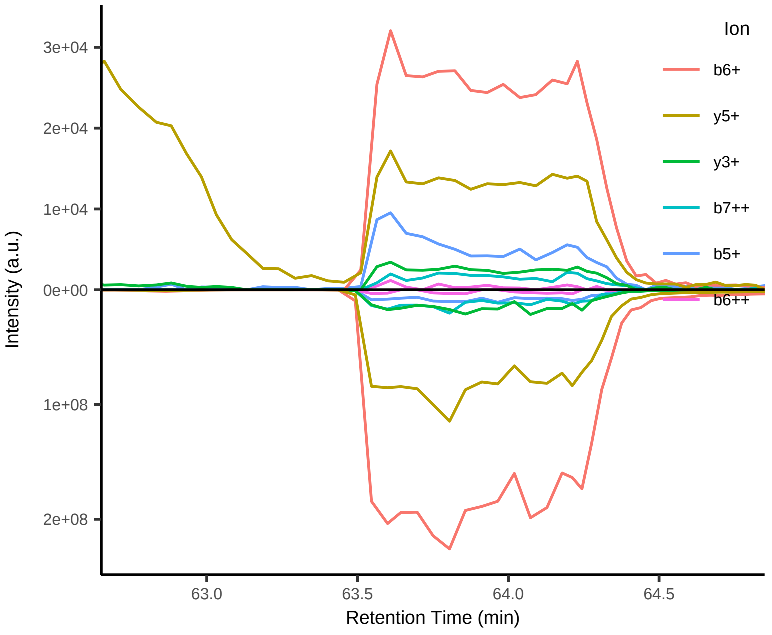


#79 LMLWL[+7.017164]PHW++

Peptide mix
Top 4 transitions: dotp = 0.95; l:h ppm = 140

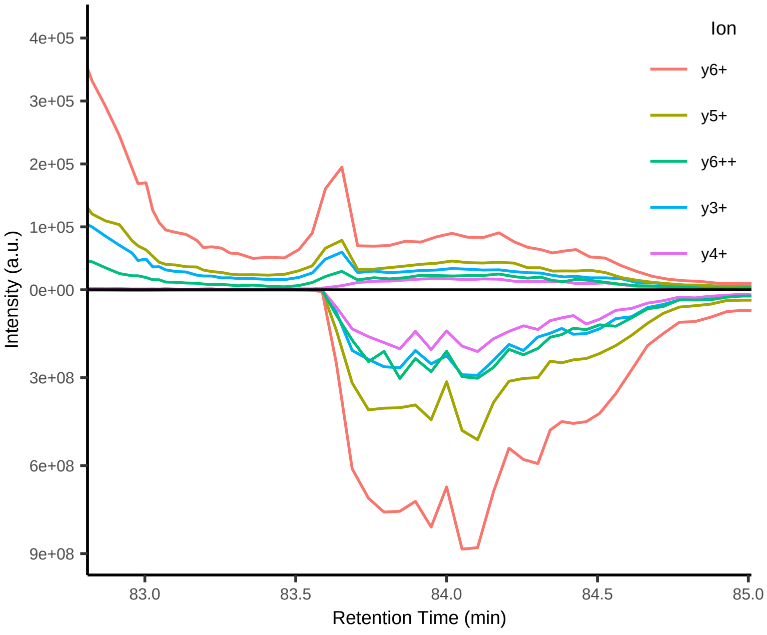


#80 MLWLPHWGL[+7.017164]Y++

Peptide mix
Top 4 transitions: dotp = 0.91; l:h ppm = 139

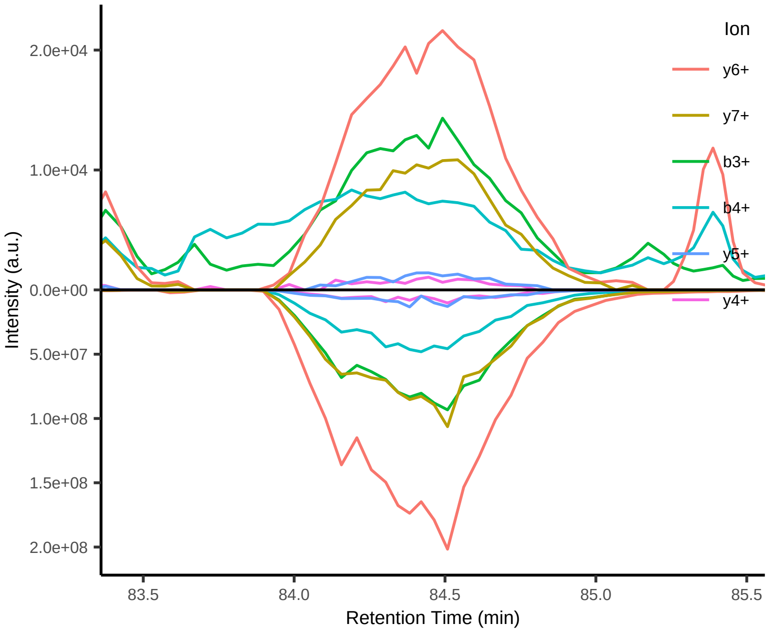


#81 KSV[+6.013809]LTAFLM[+15.994915]L++

Peptide mix
Top 4 transitions: dotp = 0.96; l:h ppm = 130

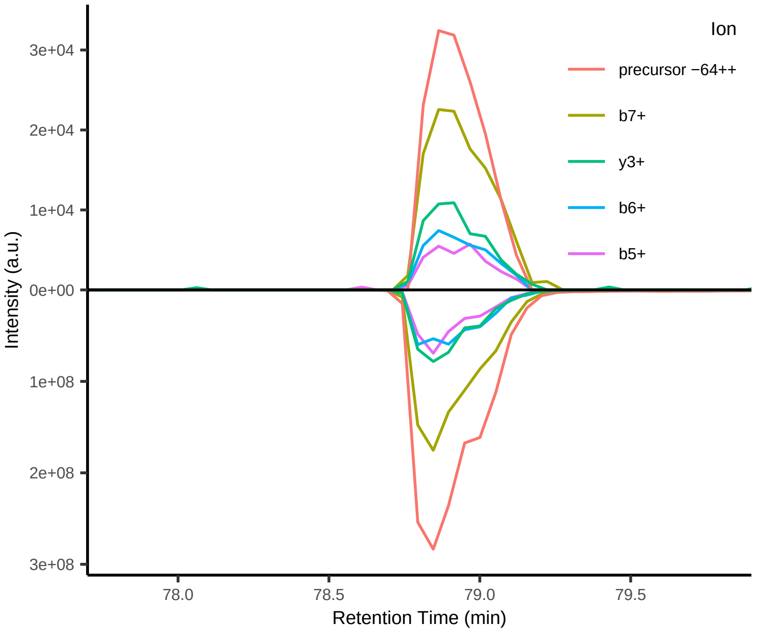


#82 M[+15.994915]LWLPHWGL[+7.017164]Y++

Peptide mix
Top 4 transitions: dotp = 0.94; l:h ppm = 127

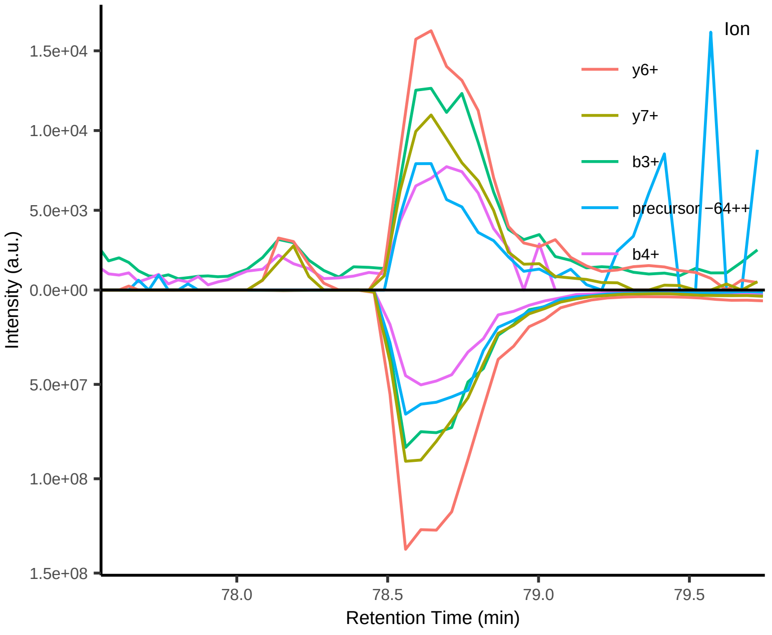


#83 GKSTL[+7.017164]VRLL+++

Peptide mix
Top 4 transitions: dotp = 0.97; l:h ppm = 131

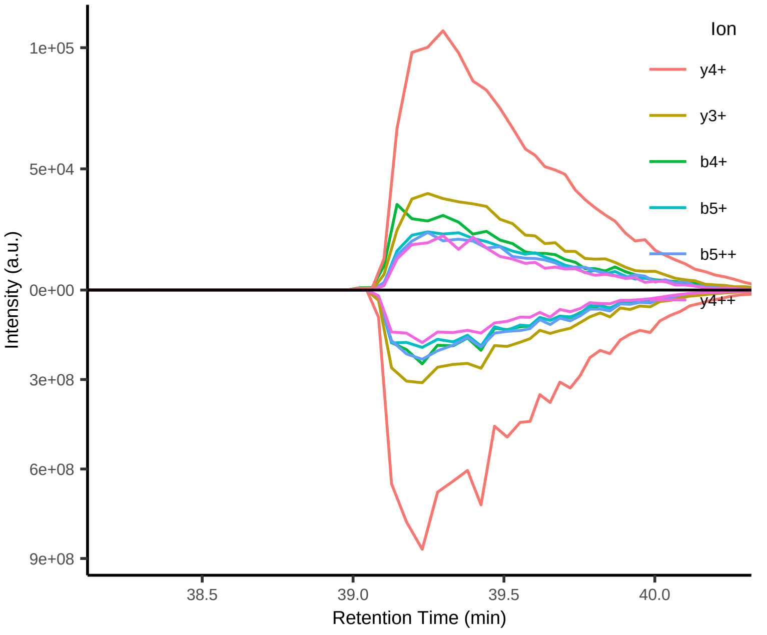

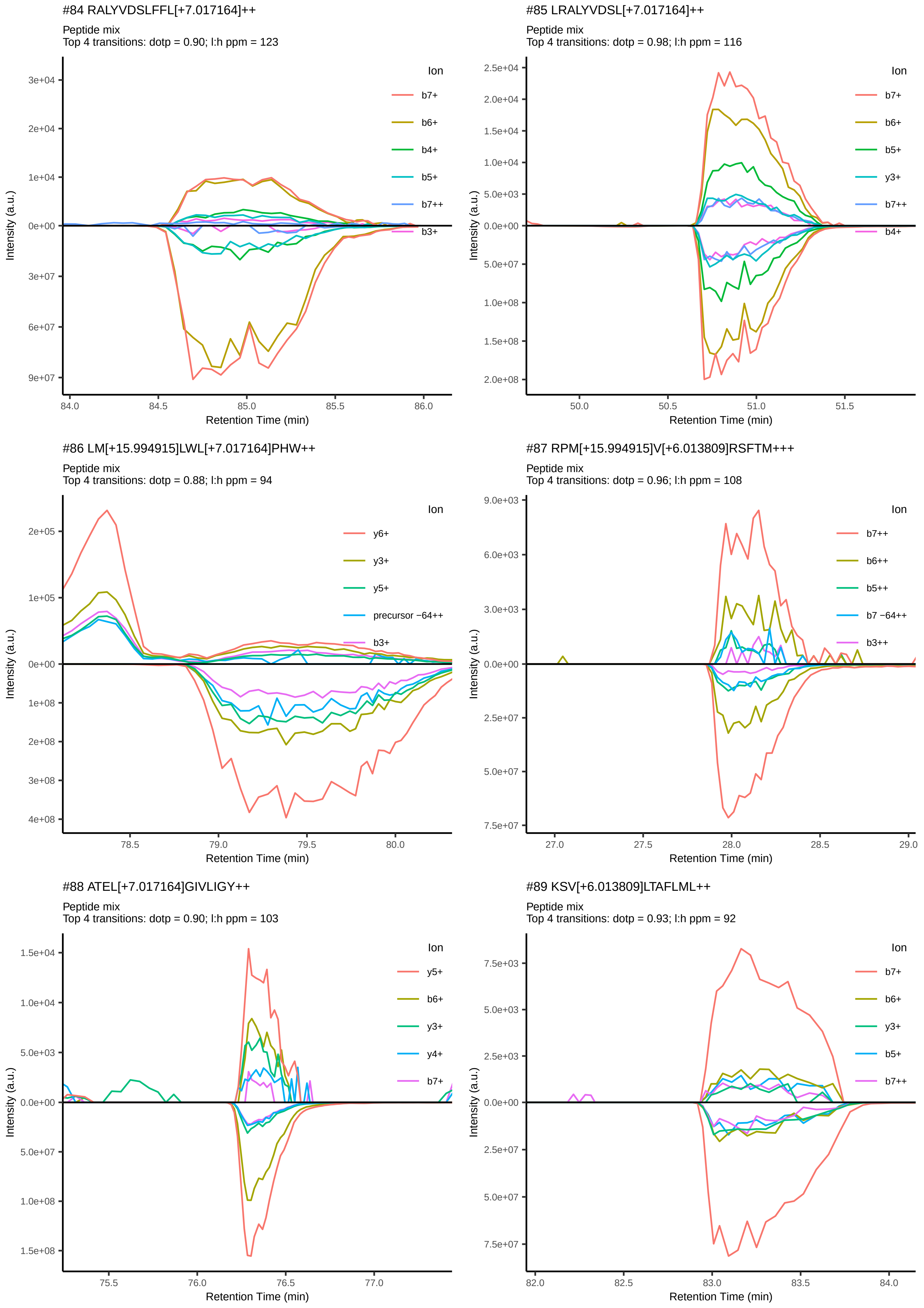
#84 RALYVDSLFFL[+7.017164]++
Peptide mix
Top 4 transitions: dotp = 0.90; l:h ppm = 123
Ion
b7+
b6+
b4+
b5+
b7++
b3+
Intensity (a.u.)
Retention Time (min)
#85 LRALYVDSL[+7.017164]++
Peptide mix
Top 4 transitions: dotp = 0.98; l:h ppm = 116
Ion
b7+
b6+
b5+
y3+
b7++
b4+
Intensity (a.u.)
Retention Time (min)
#86 LM[+15.994915]LWL[+7.017164]PHW++
Peptide mix
Top 4 transitions: dotp = 0.88; l:h ppm = 94
Ion
y6+
y3+
y5+
precursor −64++
b3+
Intensity (a.u.)
Retention Time (min)
#87 RPM[+15.994915]V[+6.013809]RSFTM+++
Peptide mix
Top 4 transitions: dotp = 0.96; l:h ppm = 108
Ion
b7++
b6++
b5++
b7 −64++
b3++
Intensity (a.u.)
Retention Time (min)
#88 ATEL[+7.017164]GIVLIGY++
Peptide mix
Top 4 transitions: dotp = 0.90; l:h ppm = 103
Ion
y5+
b6+
y3+
y4+
b7+
Intensity (a.u.)
Retention Time (min)
#89 KSV[+6.013809]LTAFLML++
Peptide mix
Top 4 transitions: dotp = 0.93; l:h ppm = 92
Ion
b7+
b6+
y3+
b5+
b7++
Intensity (a.u.)
Retention Time (min)

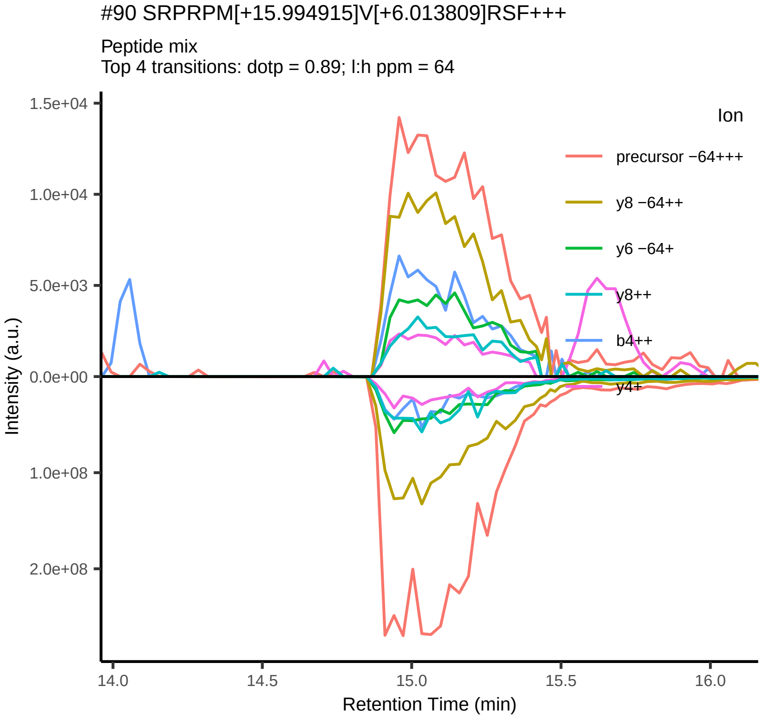
#90 SRPRPM[+15.994915]V[+6.013809]RSF+++
Peptide mix
Top 4 transitions: dotp = 0.89; l:h ppm = 64
Ion
precursor −64+++
y8 −64++
y6 −64+
y8++
b4++
y4+
Intensity (a.u.)
Retention Time (min)

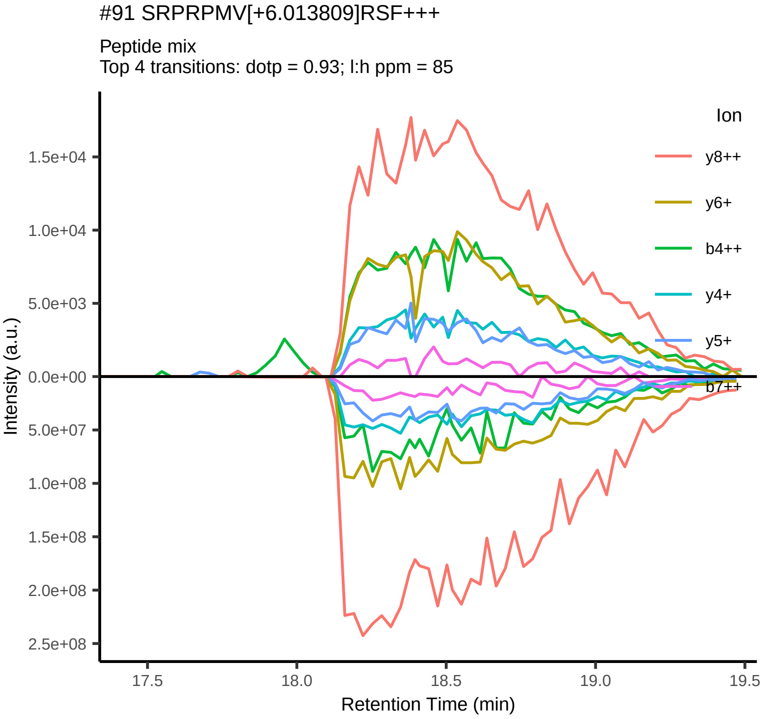
#91 SRPRPMV[+6.013809]RSF+++
Peptide mix
Top 4 transitions: dotp = 0.93; l:h ppm = 85
Ion
y8++
y6+
b4++
y4+
y5+
b7++
Intensity (a.u.)
Retention Time (min)

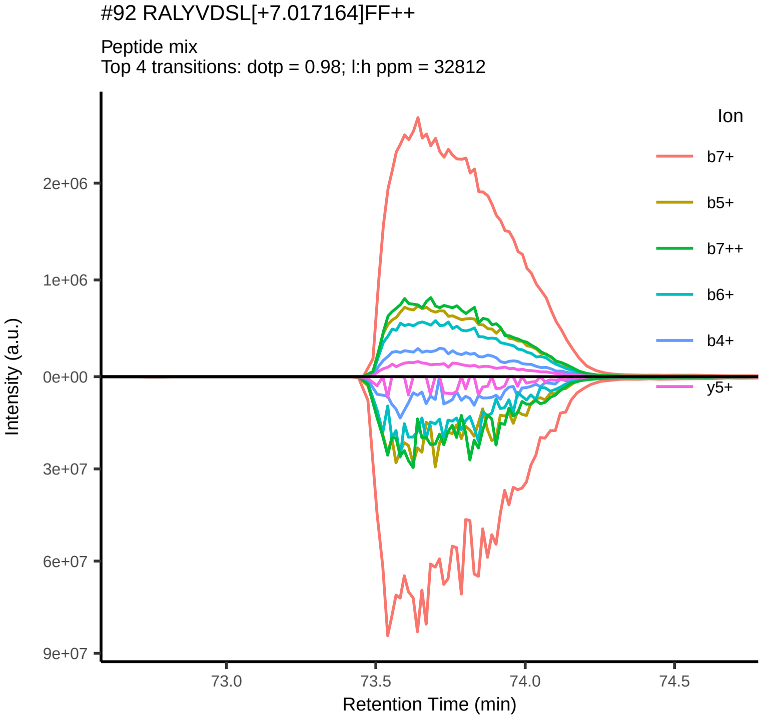
#92 RALYVDSL[+7.017164]FF++
Peptide mix
Top 4 transitions: dotp = 0.98; l:h ppm = 32812
Ion
b7+
b5+
b7++
b6+
b4+
y5+
Intensity (a.u.)
Retention Time (min)

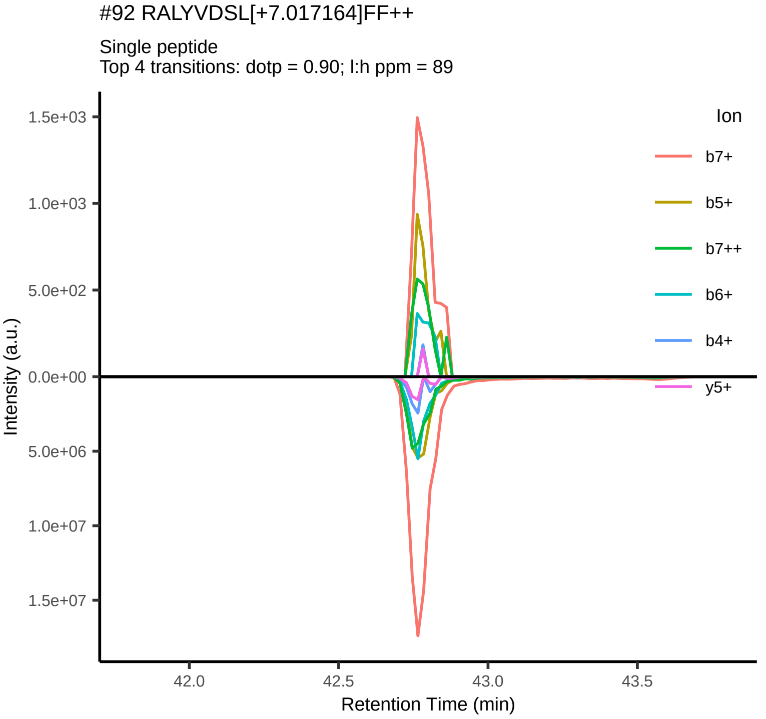
#92 RALYVDSL[+7.017164]FF++
Single peptide
Top 4 transitions: dotp = 0.90; l:h ppm = 89
Ion
b7+
b5+
b7++
b6+
b4+
y5+
Intensity (a.u.)
Retention Time (min)

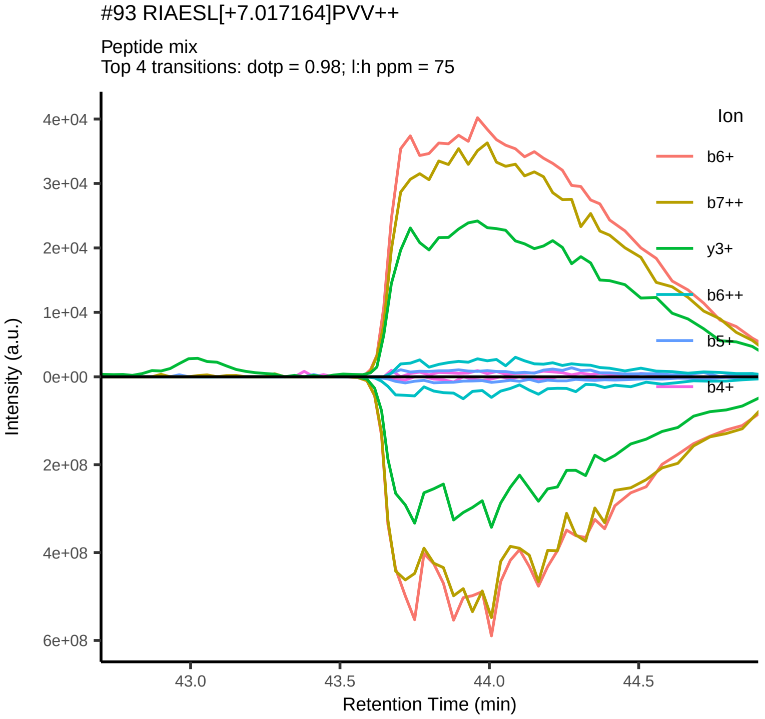
#93 RIAESL[+7.017164]PVV++
Peptide mix
Top 4 transitions: dotp = 0.98; l:h ppm = 75
Ion
b6+
b7++
y3+
b6++
b5+
b4+
Intensity (a.u.)
Retention Time (min)

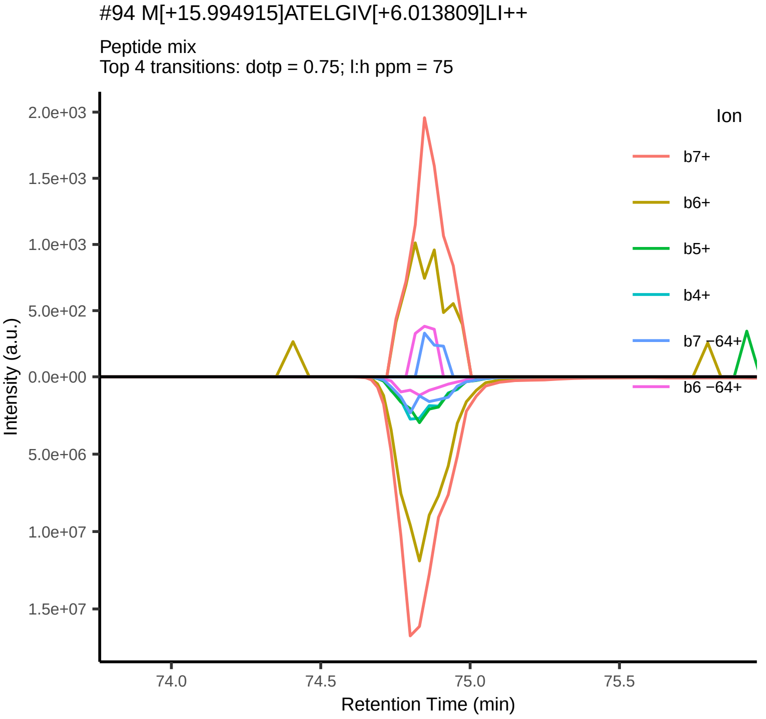
#94 M[+15.994915]ATELGIV[+6.013809]LI++
Peptide mix
Top 4 transitions: dotp = 0.75; l:h ppm = 75
Ion
b7+
b6+
b5+
b4+
b7 −64+
b6 −64+
Intensity (a.u.)
Retention Time (min)

#95 RALYVDSL[+7.017164]F++

Peptide mix
Top 4 transitions: dotp = 0.98; l:h ppm = 7025

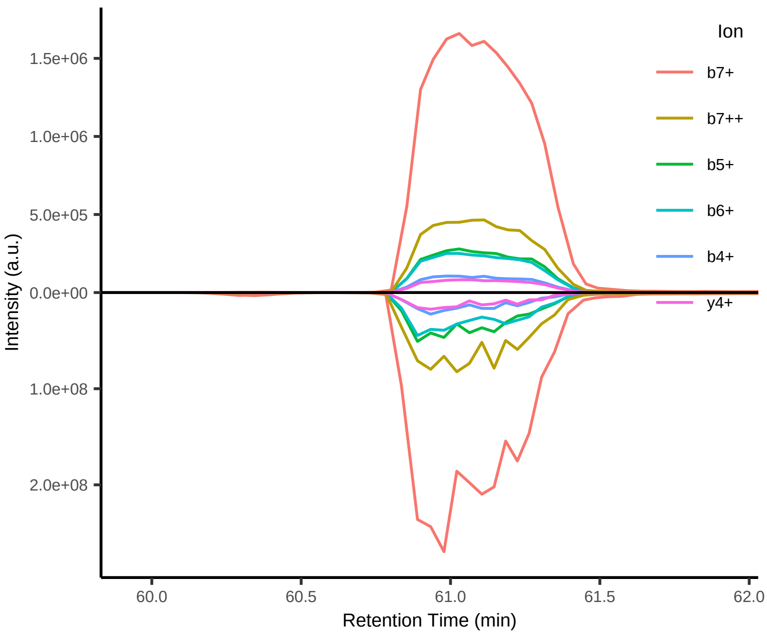


#95 RALYVDSL[+7.017164]F++

Single peptide
Top 4 transitions: dotp = 0.86; l:h ppm = 78

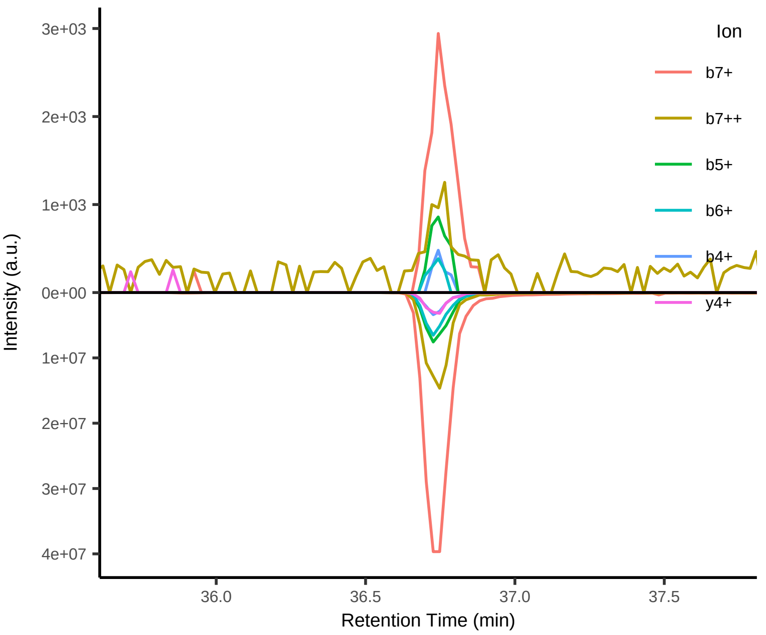


#96 ARIAESL[+7.017164]PV++

Peptide mix
Top 4 transitions: dotp = 0.96; l:h ppm = 68

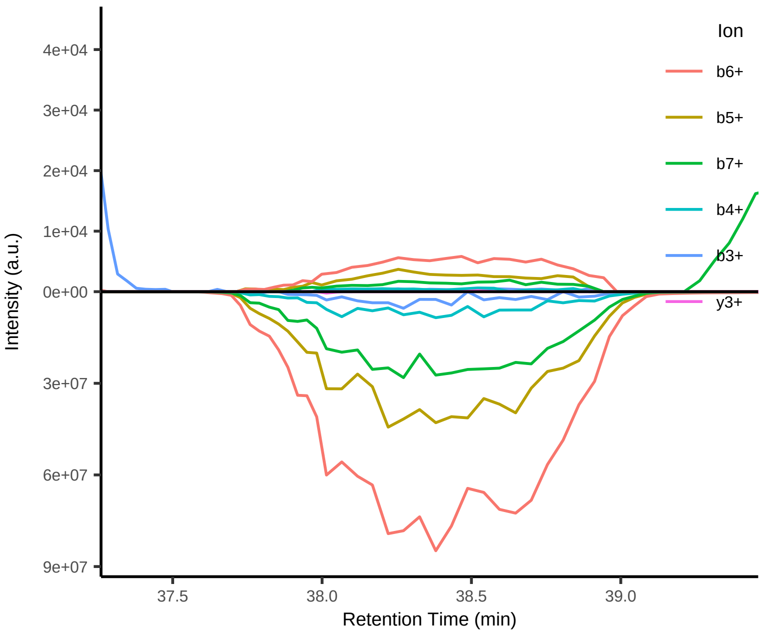


#97 MATELGIV[+6.013809]LI++

Peptide mix
Top 4 transitions: dotp = 0.93; l:h ppm = 68

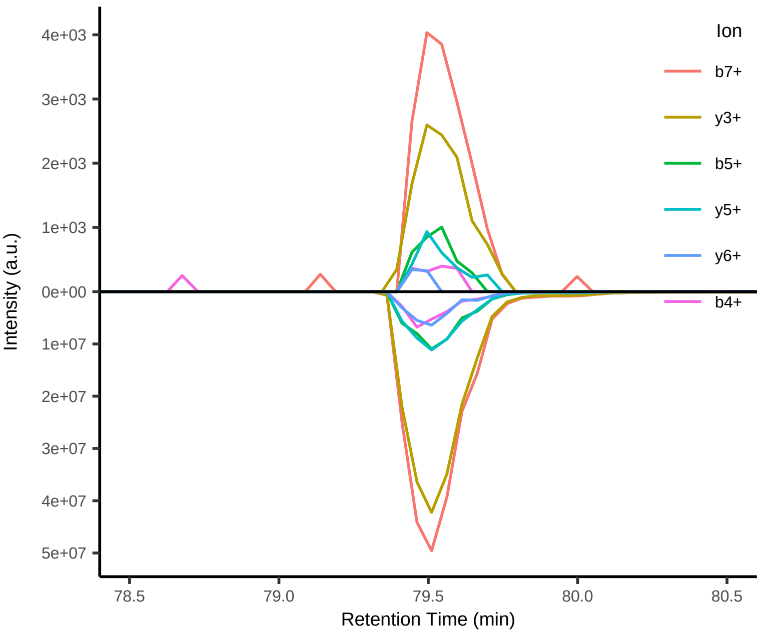


#98 RRGSRPR[+10.008269]PM[+15.994915]+++

Peptide mix
Top 4 transitions: dotp = 0.55; l:h ppm = 234

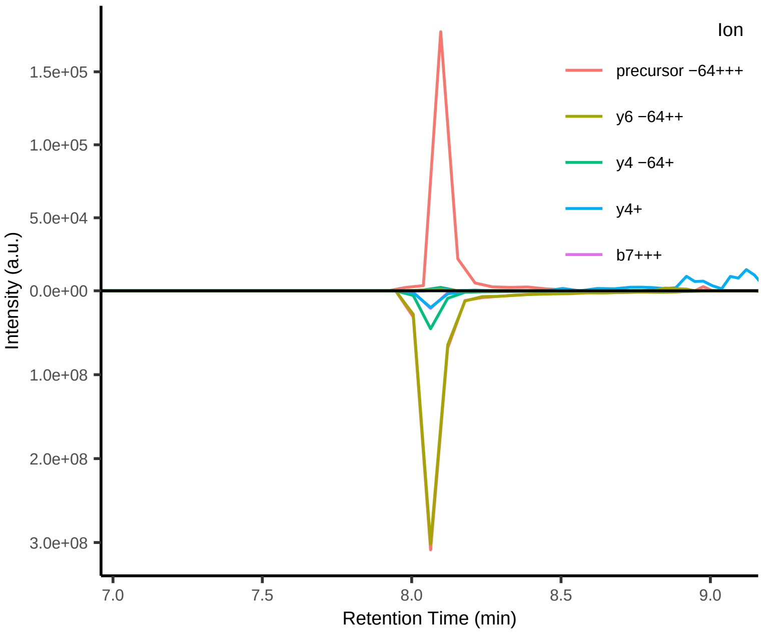


#99 M[+15.994915]ATELGI[+7.017164]V+

Peptide mix
Top 4 transitions: dotp = 0.98; l:h ppm = 11179

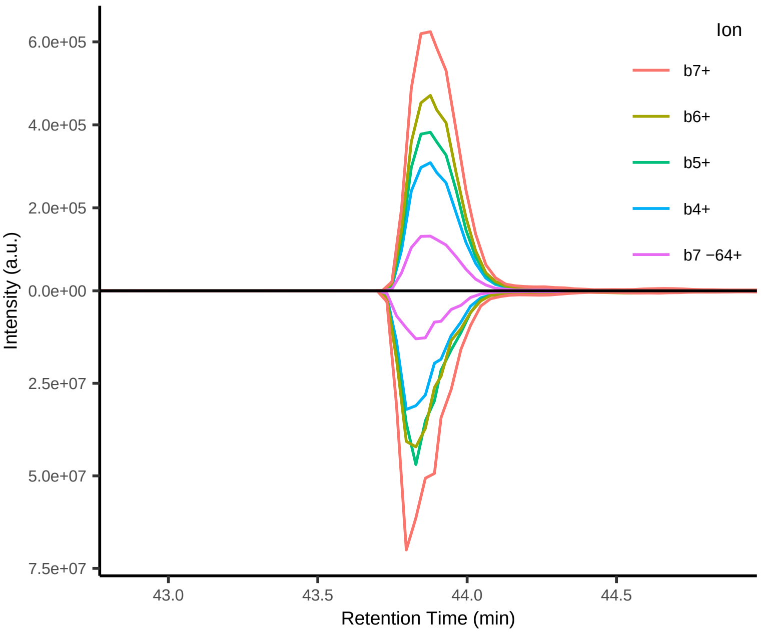

#99 M[+15.994915]ATELGI[+7.017164]V+

Single peptide
Top 4 transitions: dotp = 0.78; l:h ppm = 8

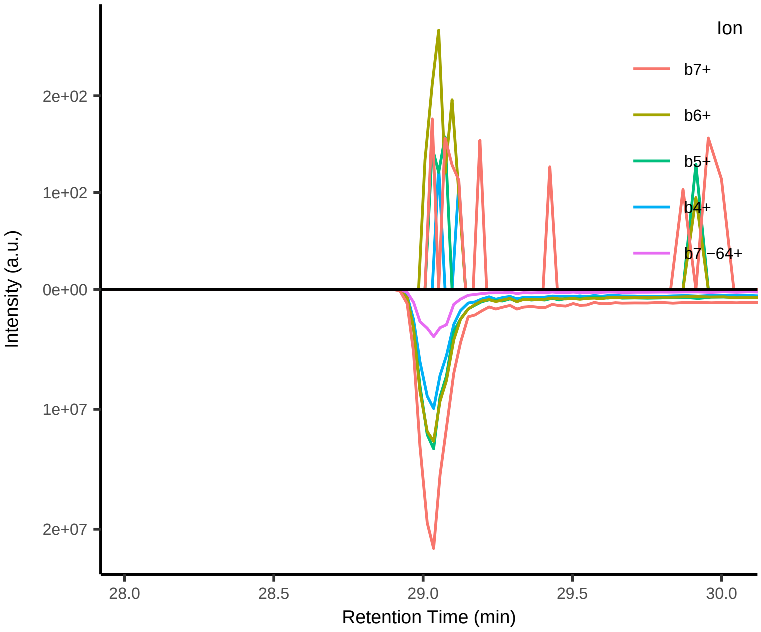


#100 MATELGI[+7.017164]V+

Peptide mix
Top 4 transitions: dotp = 0.97; l:h ppm = 7988

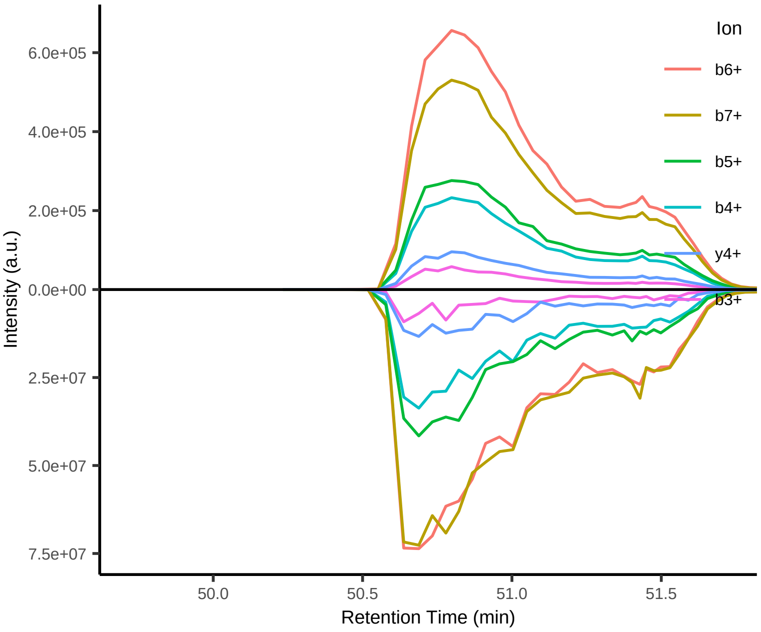


#100 MATELGI[+7.017164]V+

Single peptide
Top 4 transitions: dotp = 0.80; l:h ppm = 7

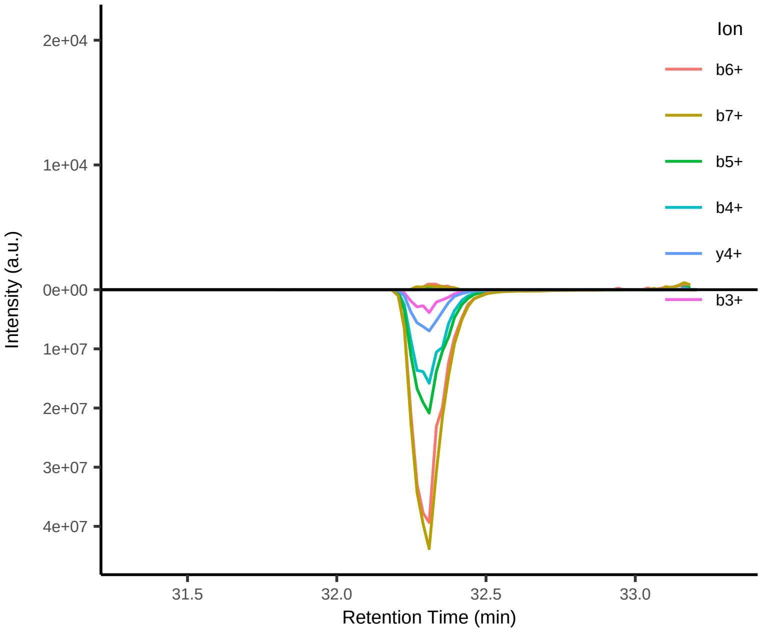


#101 GTTV[+6.013809]PFTSWK[+8.014199]++

Peptide mix
Top 4 transitions: dotp = NA; l:h ppm = NA

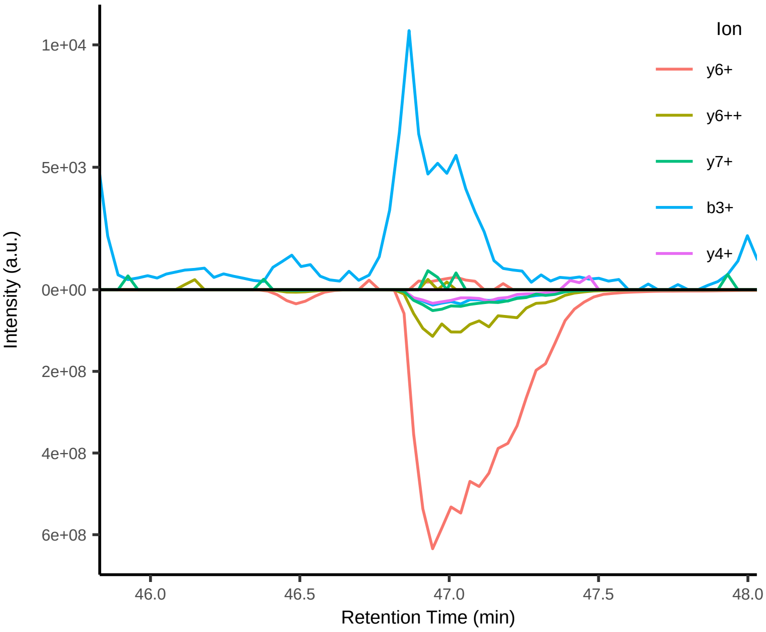


#102 FLM[+15.994915]LWL[+7.017164]PHWGL[+7.017164]++

Peptide mix
Top 4 transitions: dotp = NA; l:h ppm = NA

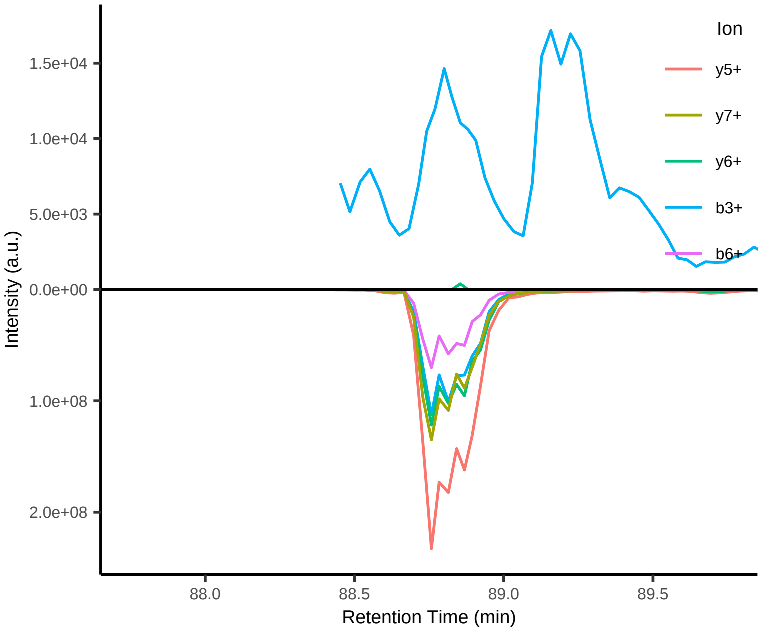


#103 KSV[+6.013809]LTAFL[+7.017164]M[+15.994915]++

Peptide mix
Top 4 transitions: dotp = 0.99; l:h ppm = 453

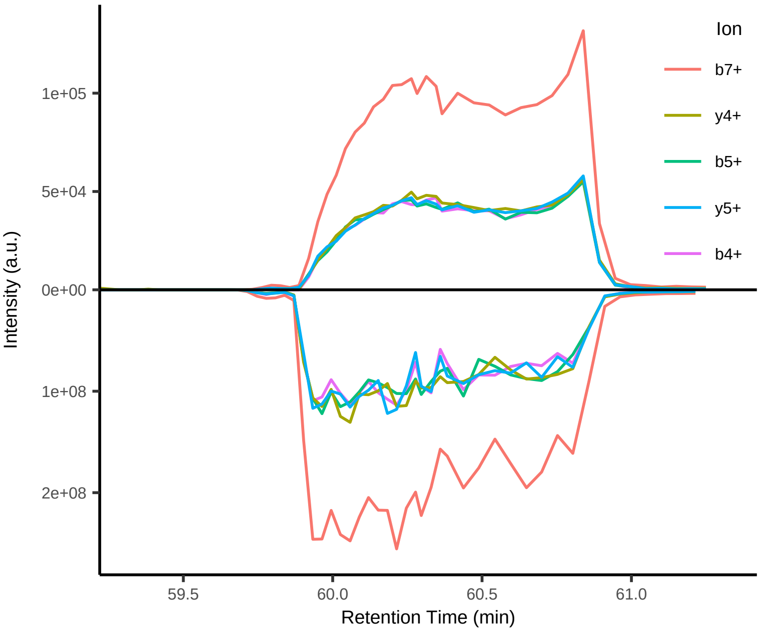

#103 KSV[+6.013809]LTAFL[+7.017164]M[+15.994915]++

Single peptide
Top 4 transitions: dotp = NA; l:h ppm = NA

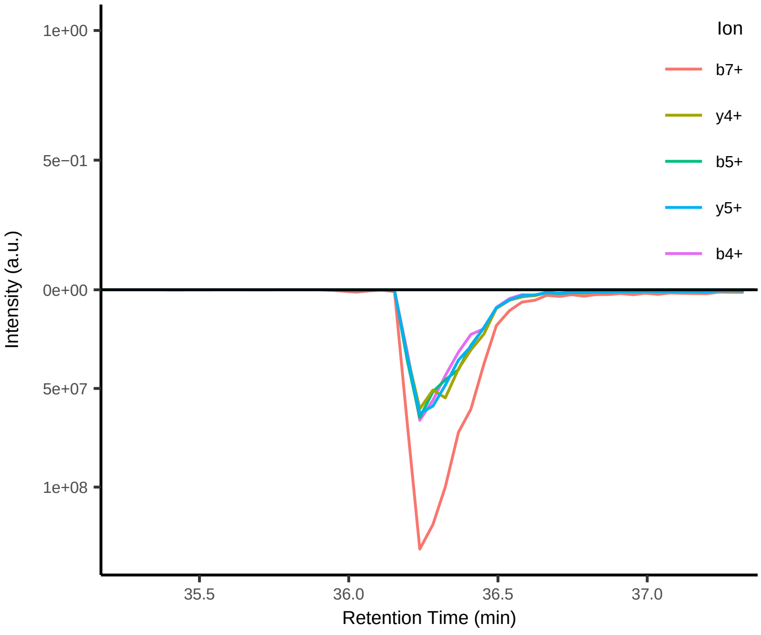


#104 STLVRLLFR[+10.008269]FY+++

Peptide mix
Top 4 transitions: dotp = NA; l:h ppm = NA

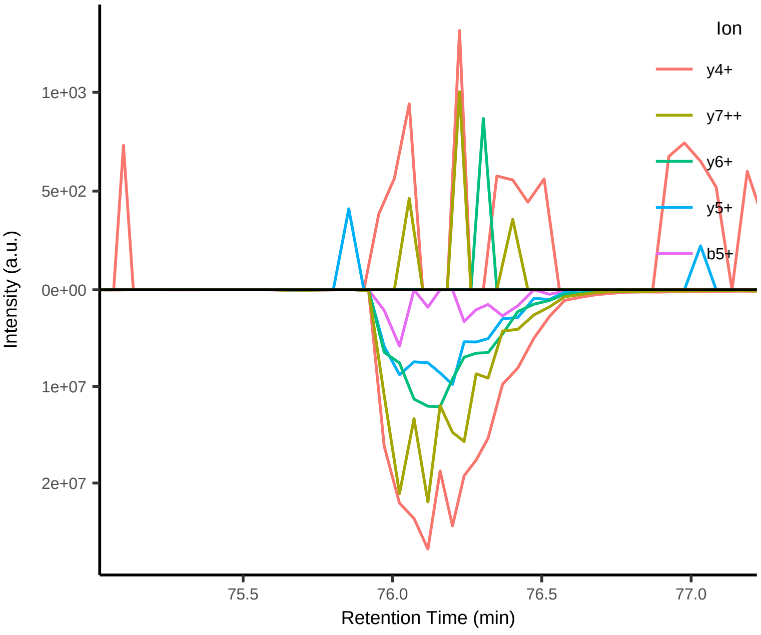


#105 STLV[+6.013809]RLLFRF+++

Peptide mix
Top 4 transitions: dotp = NA; l:h ppm = NA

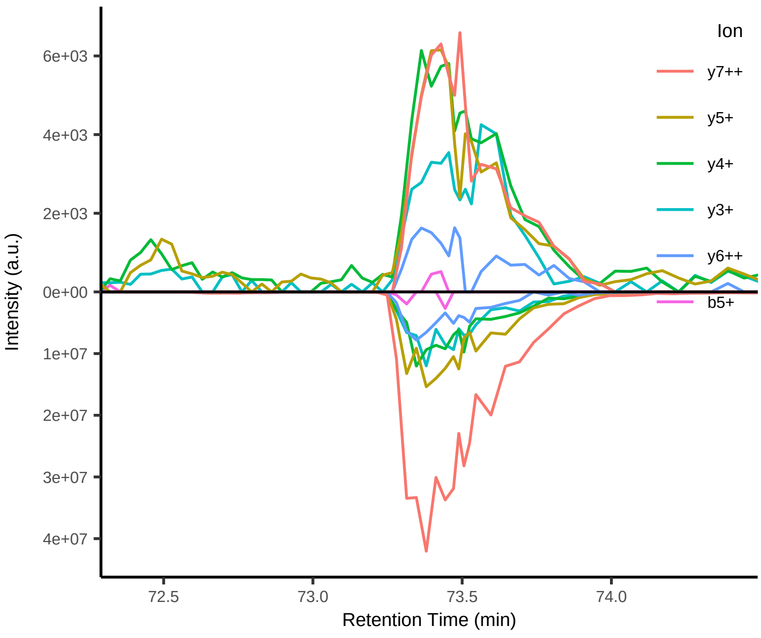


#106 RRGSRPR[+10.008269]PM+++

Peptide mix
Top 4 transitions: dotp = NA; l:h ppm = NA

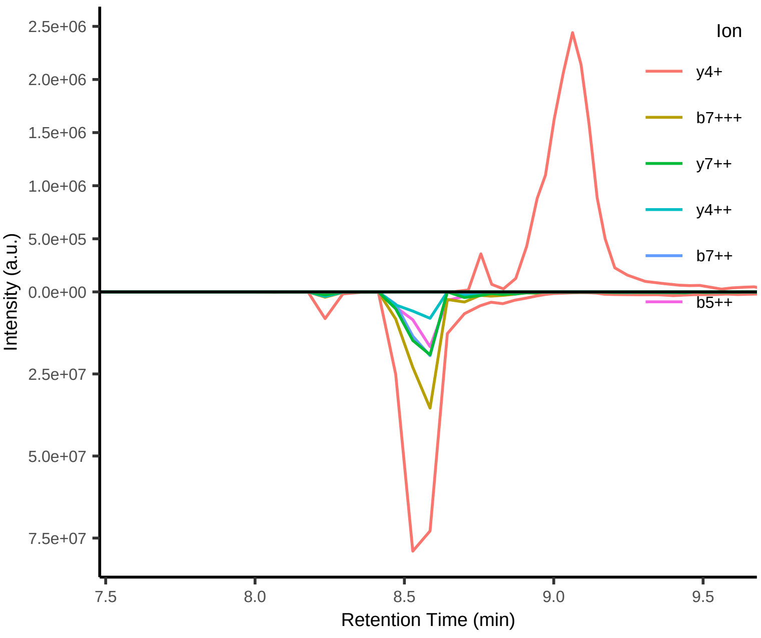


#107 VLIGYFTLV[+6.013809]++

Peptide mix
Top 4 transitions: dotp = NA; l:h ppm = NA

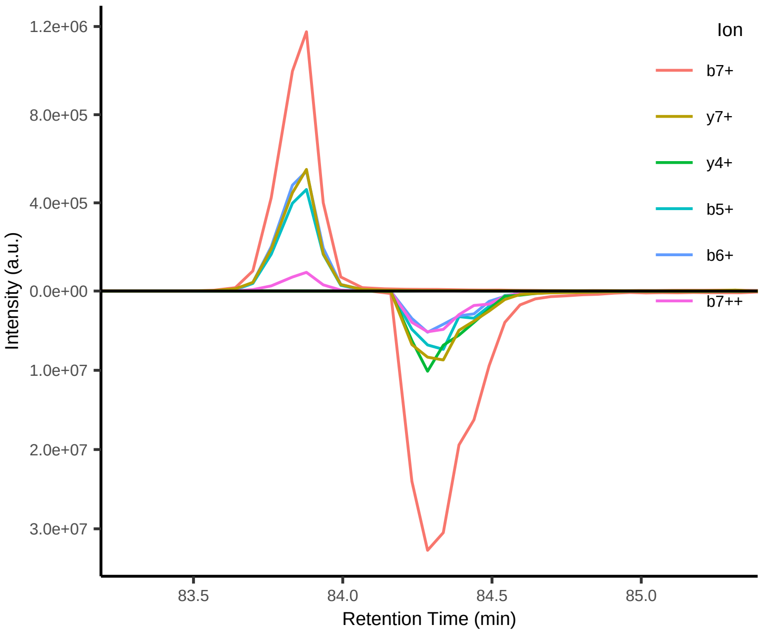


#108 M[+15.994915]ATELGIVL[+7.017164]+

Peptide mix
Top 4 transitions: dotp = NA; l:h ppm = NA

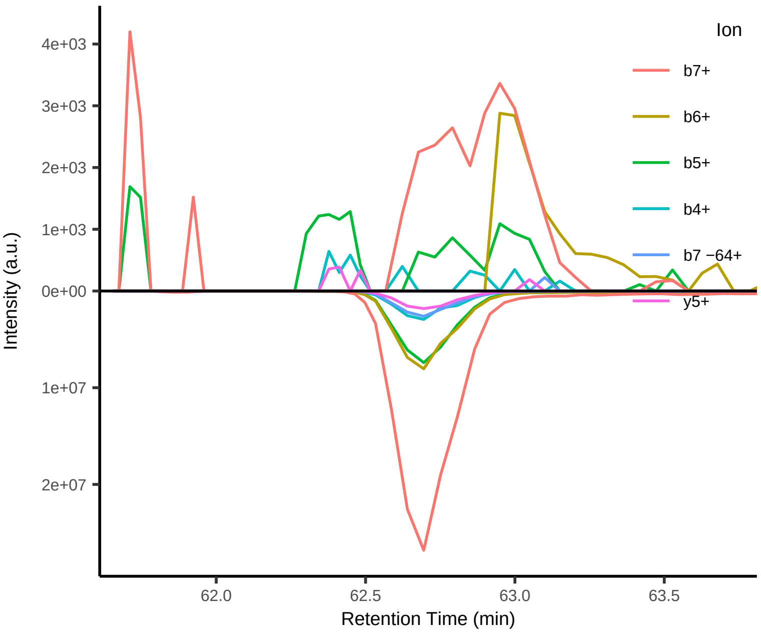

#109 ARIAESL[+7.017164]PVV++

Peptide mix
Top 4 transitions: dotp =  NA; l:h ppm = NA

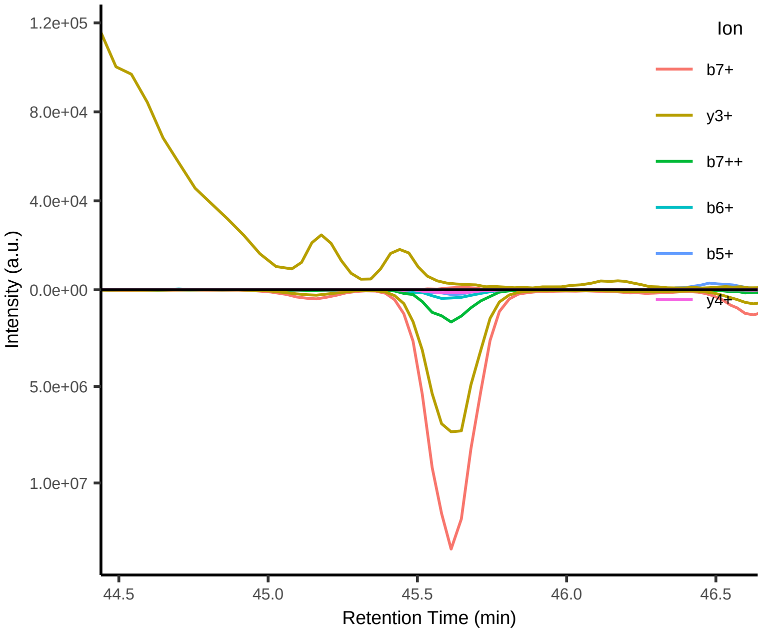


#110 FTGATII[+7.017164]EEY++

Peptide mix
Top 4 transitions: dotp =  NA; l:h ppm = NA

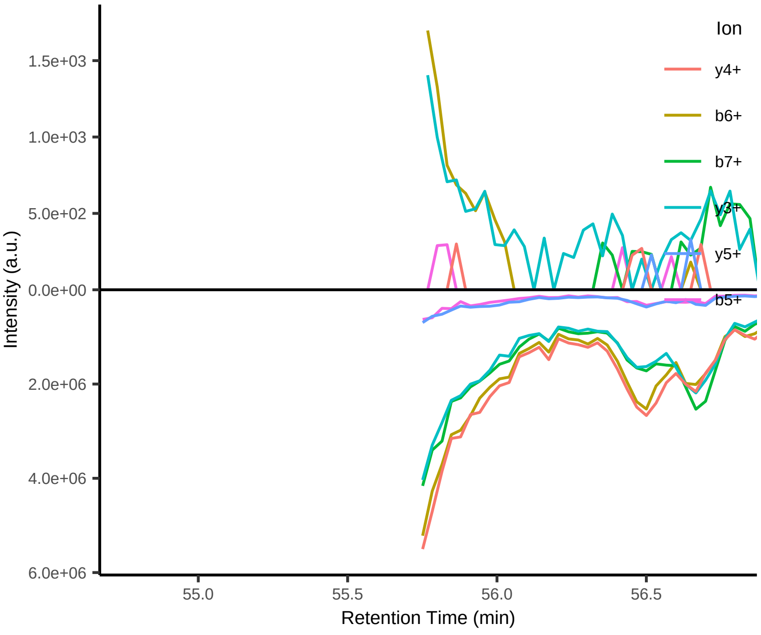


#111 SGSGKSTL[+7.017164]V++

Peptide mix
Top 4 transitions: dotp =  NA; l:h ppm = NA

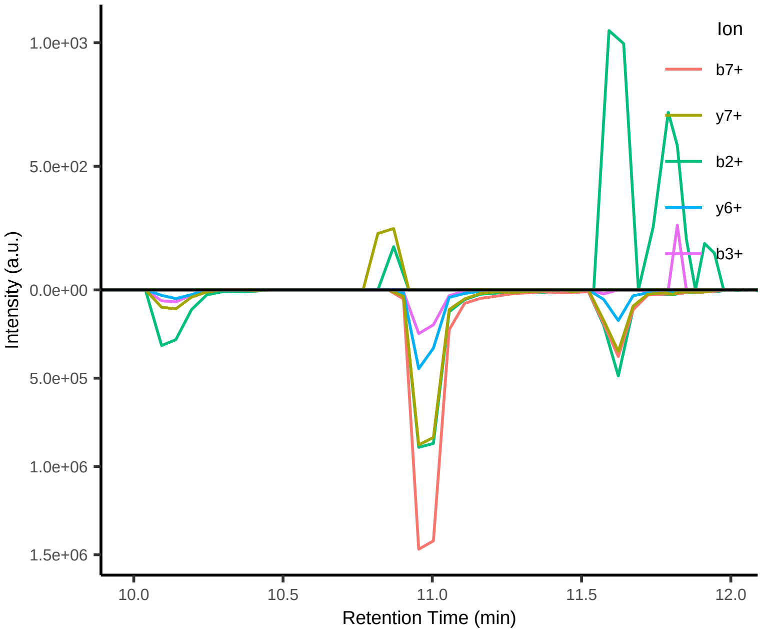


#112 SV[+6.013809]LTAFLML++

Peptide mix
Top 4 transitions: dotp =  NA; l:h ppm = NA

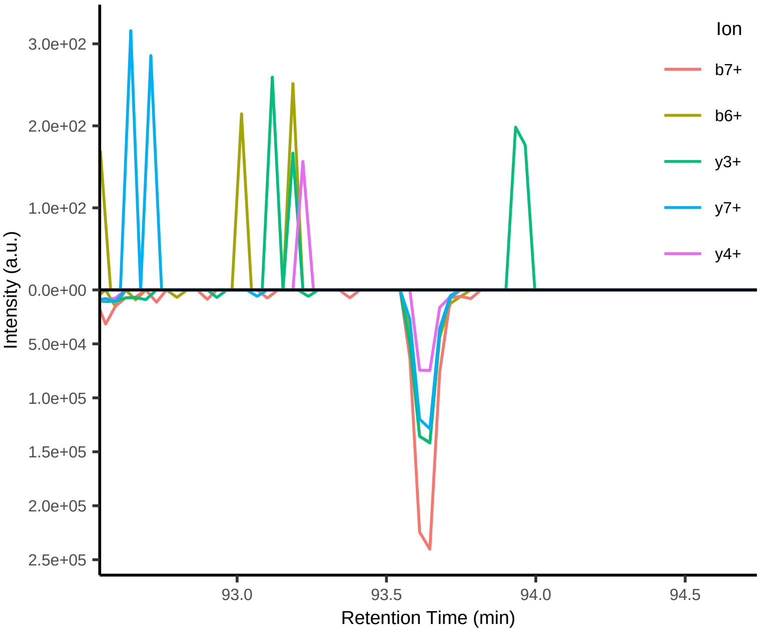


#113 KSV[+6.013809]LTAFL[+7.017164]M++

Peptide mix
Top 4 transitions: dotp = 0.98; l:h ppm = 1350

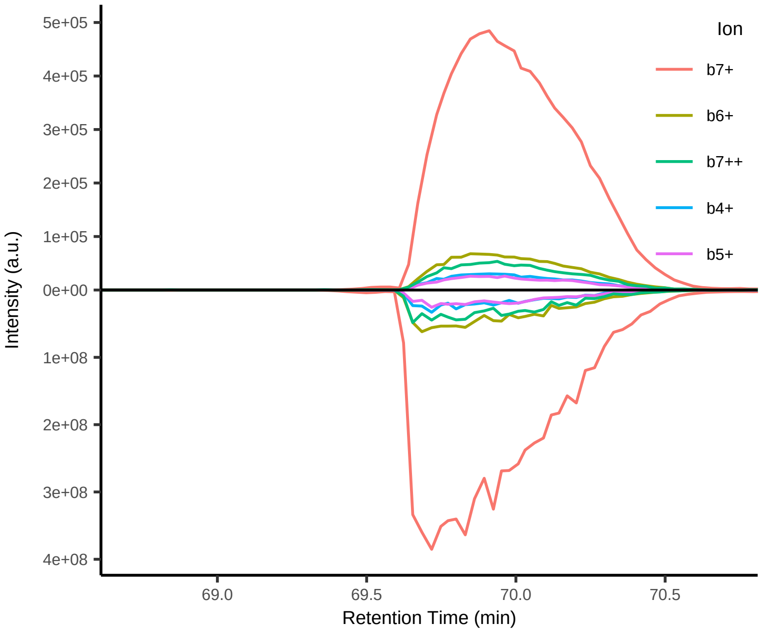


#113 KSV[+6.013809]LTAFL[+7.017164]M++

Single peptide
Top 4 transitions: dotp =  NA; l:h ppm = NA

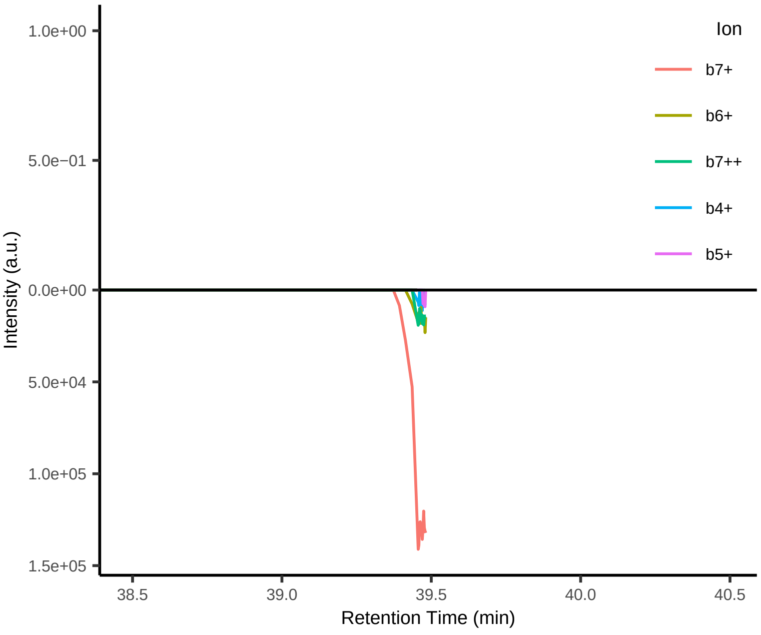
